# Structural basis of dual BACH1 regulation by SCF^FBXO22^ and SCF^FBXL17^

**DOI:** 10.1101/2024.06.03.596960

**Authors:** Benedikt Goretzki, Maryam Khoshouei, Patrick Penner, Christine Stephan, Dayana Argoti, Nele Dierlamm, Jimena Maria Rada, Sandra Kapps, Zacharias Thiel, Merve Mutlu, David Furkert, Catrin Swantje Müller, Felix Freuler, Simon Haenni, Laurent Tenaillon, Britta Knapp, Alexandra Hinniger, Philipp Hoppe, Sascha Gutmann, Grigory Ryzhakov, Enrico Schmidt, Mario Iurlaro, César Fernández

**Affiliations:** Discovery Sciences, Novartis Biomedical Research, Basel, Switzerland; Global Discovery Chemistry, Novartis Biomedical Research, Basel, Switzerland; Disease Area Oncology, Novartis Biomedical Research, Basel, Switzerland; Global Discovery Chemistry, Novartis Biomedical Research, Emeryville, USA; Disease Area Immunology, Novartis Biomedical Research, Basel, Switzerland

**Keywords:** BACH1, FBXO22, FBXL17, oxidative stress response, E3, Cullin-RING ligases, F-box, cysteine modification, S-nitrosylation, heme

## Abstract

BTB and CNC homolog 1 (BACH1) is a master transcriptional regulator of the cellular oxidative stress response and pro-metastatic oncogene. Post-translational stability of BACH1 is tightly regulated by distinct F-box ubiquitin ligases, including SCF^FBXO22^ and SCF^FBXL17^. However, the molecular details have been elusive. Here, we reveal a structural switch in FBXO22 that controls the recognition of a three-dimensional degron in the BACH1 BTB domain, thus explaining its specificity for dimeric BACH1. We describe how cancer-associated mutations in FBXO22 modulate binding and ubiquitylation of BACH1. Further, we reveal that cancer-related mutations or cysteine-modifications destabilize the BTB domain and redirect BACH1 to FBXL17, where it is recognized as a monomer. This explains how complementary ligases post-translationally regulate BACH1 depending on the state of its BTB domain. Our findings provide mechanistic insights into the regulation of the oxidative stress response and may spur therapeutic strategies to targeting oxidative stress-related disorders and metastatic cancers.

## Introduction

Cells have evolved intricate regulatory pathways to cope with oxidative stress under both physiological and pathological conditions such as chronic inflammation or cancer. These defensive mechanisms are critical in healthy or precancerous cells to safeguard against the progression to malignant states. However, the upregulation of antioxidant genes confers a growth advantage to cancer cells by acting as a countermeasure to maintain oxidative balance within the cellular environment ^1^.

The cellular response to oxidative stress is tightly regulated by the transcriptional activator nuclear factor erythroid 2-related factor 2 (NRF2) and the repressor BTB and CNC homolog 1 (BACH1). Both, NRF2 and BACH1 compete for the binding to antioxidant responsive elements (ARE), enhancer sites in the promotor regions of several cytoprotective genes^2, 3^. Under normal physiological conditions, NRF2 is inactivated in the cytoplasm through ubiquitylation by the Cullin3 ligase Kelch-like ECH-associated protein 1 (CUL3^KEAP1^) followed by proteasomal degradation and ARE sites are repressed by binding of BACH1^4^. Upon KEAP1 inactivation under oxidative stress, NRF2 translocation to the nucleus initiates the transcription of various ARE-dependent antioxidant genes, including heme oxygenase 1 (HO-1)^5^. During oxidative stress, hemoproteins can release their prosthetic heme groups, thus producing free heme, which may catalyze to the production of free radicals^6^. The binding of free heme to BACH1 has been shown to displace it from ARE sites^7^ and induce its nuclear export^8^ as well as degradation^9–11^. Notably, the NRF2-dependent expression of HO-1 leads to the degradation of heme molecules causing the stabilization of BACH1 at ARE sites thus repressing NRF2 activity. This negative feedback loop fine tunes NRF2 activity and BACH1 repression at ARE sites to maintain the cellular response to oxidative stress^3^.

The ubiquitin proteasome system is a critical layer regulating NRF2 and BACH1 activity under physiological and oxidative stress conditions. While the inactivation of NRF2 by CUL3^KEAP1^ is well understood^12^, the mechanism underlying BACH1 degradation is relatively underexplored. Two SKP1-CUL1-Fbox (SCF)-type E3 ligases have emerged to mediate the heme-dependent degradation of BACH1: SCF^FBXO22^ and SCF^FBXL17 9, 10^. However, it remains to be determined how BACH1 degradation is coordinated by these ligases and whether their roles are redundant or independent with regards to oxidative stress. Recent data suggest, that SCF^FBXO22^ is the dominant mediator for BACH1 degradation under oxidative stress and disruption of the FBXO22-BACH1 interaction leading to BACH1 stabilization in NRF2 activated cancer cells has been linked to metastasis^10^.

Here, we complement the current structural understanding of the cellular oxidative stress response with mechanistic insights into the regulation of BACH1 by SCF^FBXO22^. By combining nuclear magnetic resonance (NMR) spectroscopy and cryo-electron microscopy (cryo-EM) with hydrogen-deuterium-exchange mass spectrometry (HDX-MS) and molecular dynamics (MD) simulations, we elucidate how FBXO22 recognizes a three-dimensional degron in the Broad-Complex, Tramtrack and Bric-à-brac (BTB) domain of BACH1 through a conformational selection mechanism. FBXO22 binds the dimeric BTB domain in the BACH1 N-terminus while ubiquitylating the disordered C-terminal region. We show that BACH1 binding to FBXO22 and ubiquitylation is independent of heme which excludes a potential molecular glue mechanism for heme. Instead, our observations suggest another factor regulates the interaction of BACH1 with SCF^FBXO22^. Based on our structural insights, we identify cancer-associated mutations in FBXO22 which modulate binding and ubiquitylation of BACH1 and thus provide a rational for their potential contribution to cancer progression. Moreover, we show that destabilization of the BACH1 BTB domain through mutagenesis or cysteine-modification, including S-nitrosylation, redirects BACH1 from SCF^FBXO22^ to interact with SCF^FBXL17^ in its monomeric form. Thus, BACH1 degradation by dual ligases reflects regulation by a monomer-to-dimer switch and that FBXO22 and FBXL17 are complementary and regulate BACH1 stability under different cell states to ensure efficient BACH1 degradation under changing cellular conditions during oxidative stress.

## Results

### FBXO22 recognizes dimeric BACH1 through its BTB domain

Previous cell-based mutagenesis studies have identified residues ^9^**F**A**Y**E**S**^13^ in the BACH1 N-terminus as the degron sequence recognized by FBXO22^10^. The sequence mediates BTB dimerization through a domain swapped β-sheet (Figure 1a) and we thus anticipated that FBXO22 binds the dimeric BTB domain of BACH1.

**Figure 1:**
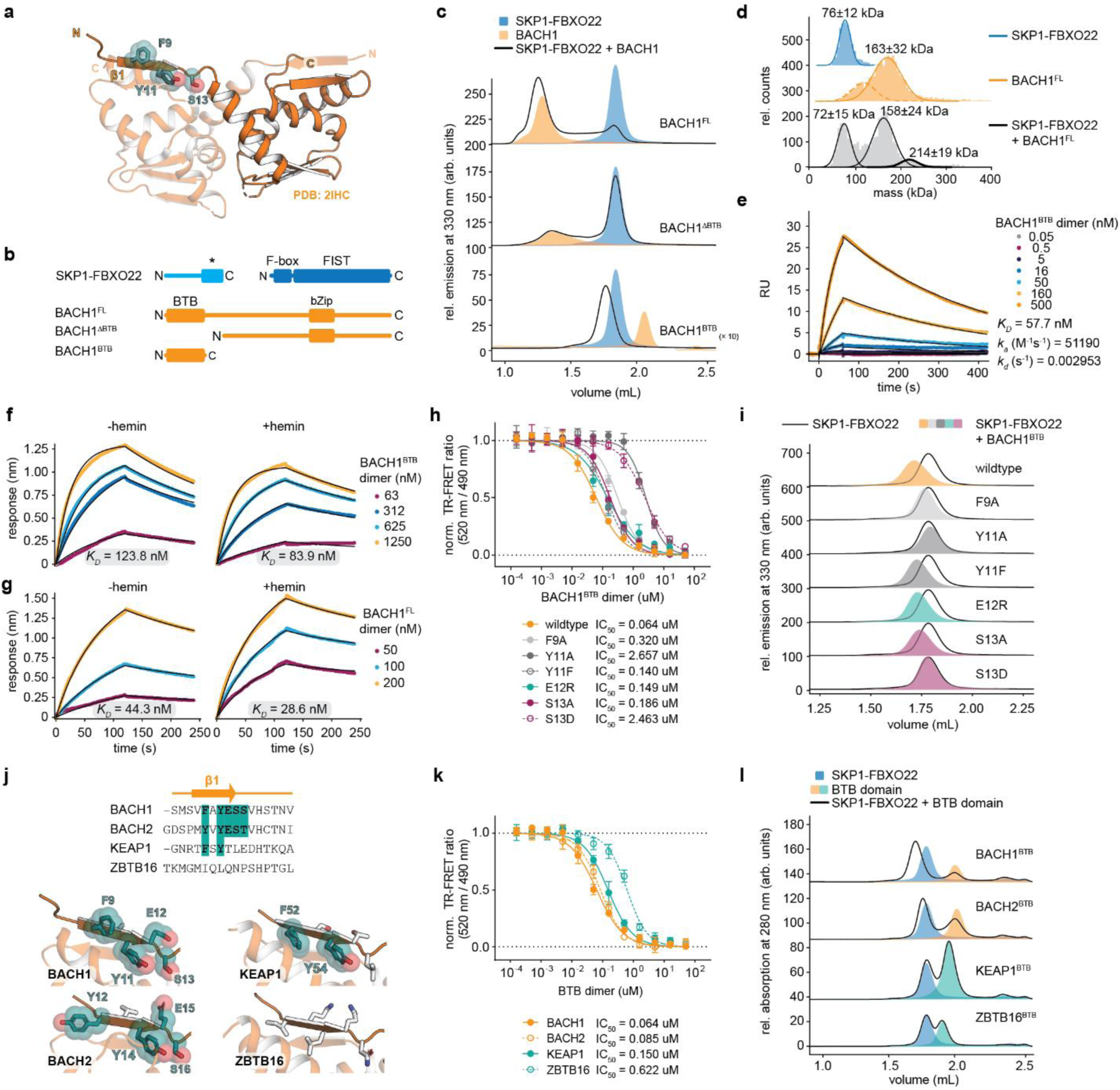
BACH1 binds to FBXO22 as a dimer via its BTB domain independent of heme. **a** X-ray crystal structure of the BACH1 BTB domain (PDB ID: 2IHC) highlighting the previously identified FBXO22-dependent degron ^9^**F**A**Y**E**S**^13^ in the domain-swapped N-terminal β-sheet^10^ **b** Overview of purified SKP1-FBXO22 and BACH1 constructs. FIST: short for F-box and intracellular signal transduction; *: F-box interacting motif; BTB: Broad-Complex, Tramtrack and Bric a brac domain; bZip: basic leucine zipper, heterodimerization with small musculoaponeurotic fibrosarcoma (Maf) proteins. **c** SEC analysis of complex formation between SKP1-FBXO22 and BACH1 constructs. Proteins were used in equimolar ratios, i.e., one SKP1-FBXO22 heterodimer per BACH1 homodimer. **d** Mass photometric (MP) analysis of SKP1-FBXO22 (calculated molecular weight, *M*_W_,_calc._=63 kDa; experimental molecular weight *M*_W_,_exp._=76 ± 12 kDa), BACH1^FL^ (*M*_W_,_calc._=180 kDa, 163 ± 32 kDa), and the SKP1-FBXO22/BACH1^FL^ complex (*M*_W_,_exp._=214 ± 19 kDa) suggests a complex stoichiometry of (SKP1)_1_(FBXO22)_1_(BACH1^FL^)_2_ (*M*_W_,_calc._=227 kDa). **e, f, g** SPR (**e**) and BLI (**f**, **g**) analysis of immobilized biotinylated ^Avi^SKP1-FBXO22 with BACH1 constructs. BLI measurements were carried out in the absence and presence of 5 µM hemin. All sensorgrams were analyzed with a kinetic fit assuming a 1:1 binding mode between SKP1-FBXO22 and BACH1 dimers. **h** TR-FRET competition assay evaluating the ability of BACH1^BTB^ mutants and BACH2, KEAP1, and ZBTB16 BTB domains to compete with wildtype BACH1^BTB^ (His-tagged, bound to Tb-conjugated Anti-His antibody) for binding to SKP1-FBXO22 (biotinylated, bound to AF488-conjugated Streptavidin). **i** Fluorescence-detected SEC assay (fSEC) to monitor complex formation between SKP1-FBXO22 and BACH1^BTB^ mutants. **j** Sequence and structural alignment of β1 strand in the BTB domains of BACH1, BACH2, KEAP1, and ZBTB16. Conserved residues in the FBXO22-dependent degron of the BACH1 BTB domain are highlighted in cyan. BACH1^BTB^ PDB ID: 2IHC, BACH2^BTB^ PDB ID: 3OHU; KEAP1^BTB^ PDB ID: 4XCI; ZBTB16^BTB^ PDB ID: 1BUO. Sequence similarities between BTB domains as indicated in the main text was calculated with the Sequence Manipulation Suite^75^. **k** TR-FRET competition assay evaluating the ability of the BACH2, KEAP1, and ZBTB16 BTB domains to compete with wildtype BACH1^BTB^ (His-tagged, bound to Tb-conjugated Anti-His antibody) for binding to SKP1-FBXO22 (biotinylated, bound to AF488-conjugated Streptavidin). **l** fSEC assay to monitor complex formation between SKP1-FBXO22 and the BACH2, KEAP1, and ZBTB16 BTB domains.

To investigate the interaction between FBXO22 and BACH1, we expressed and purified the complex of SKP1 (residues 2-163) and FBXO22 (residues 12-403), full-length BACH1 (residues 2-736, termed BACH1^FL^), a BACH1 mutant lacking the BTB domain (residues 174-736, BACH1^ΔBTB^) as well as the isolated BACH1 BTB domain (residues 7-128, termed BACH1^BTB^) (Figure 1b, Supplementary Table 1). Size-exclusion chromatography coupled to multi-angle light scattering (SEC-MALS) and circular dichroism (CD) spectroscopy confirmed the structural integrity of the SKP1-FBXO22 heterodimer and the BACH1 homodimers in solution (Supplementary Figure 1a-c). Fluorescence-detection size exclusion chromatography (fSEC) (Figure 1c) and native PAGE analysis (Supplementary Figure 1d) identified the BACH1 BTB domain as the minimal region required for an interaction with FBXO22 (Figure 1c and d). Furthermore, mass photometry (MP) and SEC coupled to multi-angle light scattering (SEC-MALS) suggests that BACH1 binds as a dimer to SKP1-FBXO22 (Figure 1e, Supplementary Figure 1b).

To determine the affinity and kinetics of the BACH1-FBXO22 interaction, we tested the binding of BACH1^BTB^ and BACH1^FL^ to biotinylated ^Avi^SKP1-FBXO22 via surface plasmon resonance (SPR) (Figure 1e) and biolayer interferometry (BLI) (Figure 1f). Kinetically fitting the sensorgrams with a 1:1 binding model (one BACH1^BTB^ dimer per SKP1-FBXO22) revealed an affinity of *K*_D_ =57.7 nM (SPR) and *K*_D_ =123.8 nM (BLI) and with comparable association and dissociation rate constants (*k*_a_=5.1×10^5^ M^-1^s^-1^ and *k*_d_=3.0 ×10^-3^ s^-1^ by SPR; *k*_a_=2.7×10^5^ M^-1^s^-1^ and *k*_d_=3.3×10^-3^ s^-1^ by BLI) (Figure 1f). Importantly, we determined a similar affinity and binding kinetics for BACH1^FL^ (*K*_D_=44.3 nM; *k*_a_=4.7×10^5^ M^-1^s^-1^; *k*_d_=2.1×10^-3^ s^-1^) (Figure 1g). Considering the role of heme as the physiological trigger for BACH1 degradation, we probed a potential influence of hemin on the FBXO22-BACH1 interaction. Importantly, analytical SEC and nuclear magnetic resonance spectroscopy (NMR) confirmed that recombinant BACH1^FL^ was free of heme and binds heme through CP motifs in its C-terminus but not in the BTB domain (Supplementary Figure 1e and f). We determined very similar affinities for BACH1^BTB^ and BACH1^FL^ to FBXO22 in the presence of 5 µM hemin (*K*_D_=83.9 nM for BACH1^BTB^/hemin and *K*_D_=28.6 nM for BACH1^FL^/hemin) (Figure 1f and g). These observations suggest that BACH1 binding to FBXO22 is mediated solely by the BTB domain and independent of heme bound to the BACH1 C-terminus.

We next validated the previously described binding epitope using a time-resolved fluorescence energy transfer (TR-FRET) assay in which we monitored the ability of untagged BACH1^BTB^ mutants with single point mutations in the BACH1 degron to compete with His_6_-tagged BACH1^BTB^ for binding to biotinylated ^Avi^SKP1-FBXO22 (Figure 1h). SEC-MALS confirmed that all mutant proteins still formed BTB dimers (Supplementary figure 2a) even though the thermal stability of several mutants was severely decreased as demonstrated by nano differential scanning fluorimetry (nano-DSF) (Supplementary figure 2b-d). While wildtype BACH1^BTB^ was able to compete the interaction at an IC_50_ value of 64 nM, the mutations F9A, Y11A, and S13D showed markedly increased IC_50_ values of 320 nM, 2657 nM, and 2463 nM, respectively, suggesting a compromised ability to interact with SKP1-FBXO22. Likewise, complex formation of the BACH1^BTB^ mutants with SKP1-FBXO22 was severely compromised in fSEC and nano-DSF experiments (Figure 1i, Supplementary figure 3a-e). By contrast, the mutation E12R only had a subtle impact on the BACH1-FBXO22 interaction in the TR-FRET (IC_50_=149 nM) and fSEC assay (Figure 1h and i). This agrees well with previous observations in cellular studies finding that mutations of F9, Y11, and S13A in BACH1 failed to co-immunoprecipitate with FBXO22 while mutations in E12 did not^10^. Notably, the comparably mild mutations Y11F and S13A had only subtle effects on the BACH1-FBXO22 interaction in the TR-FRET (IC_50_=140 nM and 183 nM, respectively) and fSEC assay (Figure 1h and i) while co-immunoprecipitation with FBXO22 was compromised in previous cellular studies^10^ This suggests that F9, Y11, and S13 in BACH1 are key residues for the interaction with FBXO22 and even perturbations with subtle effects on complex formation in vitro can abrogate the interaction on the cellular level.

The low tolerance to mutations in β1 suggests that FBXO22 is quite specific for BACH1 rather than being a universal receptor for BTB domains as previously observed for FBXL17^13, 14^. We thus tested the binding of FBXO22 to BTB domains with varying sequence similarity to BACH1^BTB^ (Figure 1j, Supplementary figure 4a and b). While BACH2^BTB^ (78.2% sequence similarity to BACH1, 64.1% similarity in β1) was still able to compete with BACH1^BTB^ at an IC_50_ of 85 nM, in the TR-FRET competition assay, KEAP1^BTB^ (42.4%/37.5%) and ZBTB16^BTB^ domains (44.0%/29.4%) competed with BACH1^BTB^ domain at increased IC_50_ values of 150 nM and 622 nM, respectively (Figure 1k). Likewise, BACH2^BTB^ showed complex formation in analytical SEC experiments while KEAP1^BTB^ and ZBTB16^BTB^ did not (Figure 1l). This highlights the specificity of FBXO22 towards the BTB domain found in BACH1.

### FBXO22 and BACH1^BTB^ interact through an intermolecular β-sheet

To investigate the FBXO22-BACH1 interaction in more detail, we employed NMR spectroscopy using uniformly ^15^N-labeled BACH1^BTB^ (U-^15^N BACH1^BTB^) (Supplementary Figure 5a and b). The (^1^H,^15^N)-HSQC NMR spectrum indicates a well folded and symmetric dimer. Upon addition of two equivalents unlabeled SKP1-FBXO22, severe peak broadening occurred throughout the spectrum due to the formation of a large 96 kDa complex (Supplementary Figure 5c). This precluded a straightforward chemical shift perturbation (CSP) analysis (Supplementary Figure 5d). However, we noticed that residues such as S17, I97, N117, E119 close to β1 split upon SKP1-FBXO22 binding (Figure 2a). A similar effect was observed in (^1^H,^13^C)-HMQC NMR experiments of ^13^CH_3_-labeled BACH1^BTB^ (U-(^2^H, ^15^N), (ILVMA)-^13^CH_3_ labeled BACH1^BTB^) (Figure 2b). The peak, corresponding to a non-native methionine residue in the BACH1^BTB^ N-terminus directly preceding the ^9^**F**A**Y**E**S**^13^ degron sequence split up in a concentration dependent manner with a saturation at one equivalent SKP1-FBXO22. This behavior reflects a symmetry break in the BTB domain, e.g., due to one SKP1-FBXO22 heterodimer binding to one subunit of the BTB dimer.

**Figure 2:**
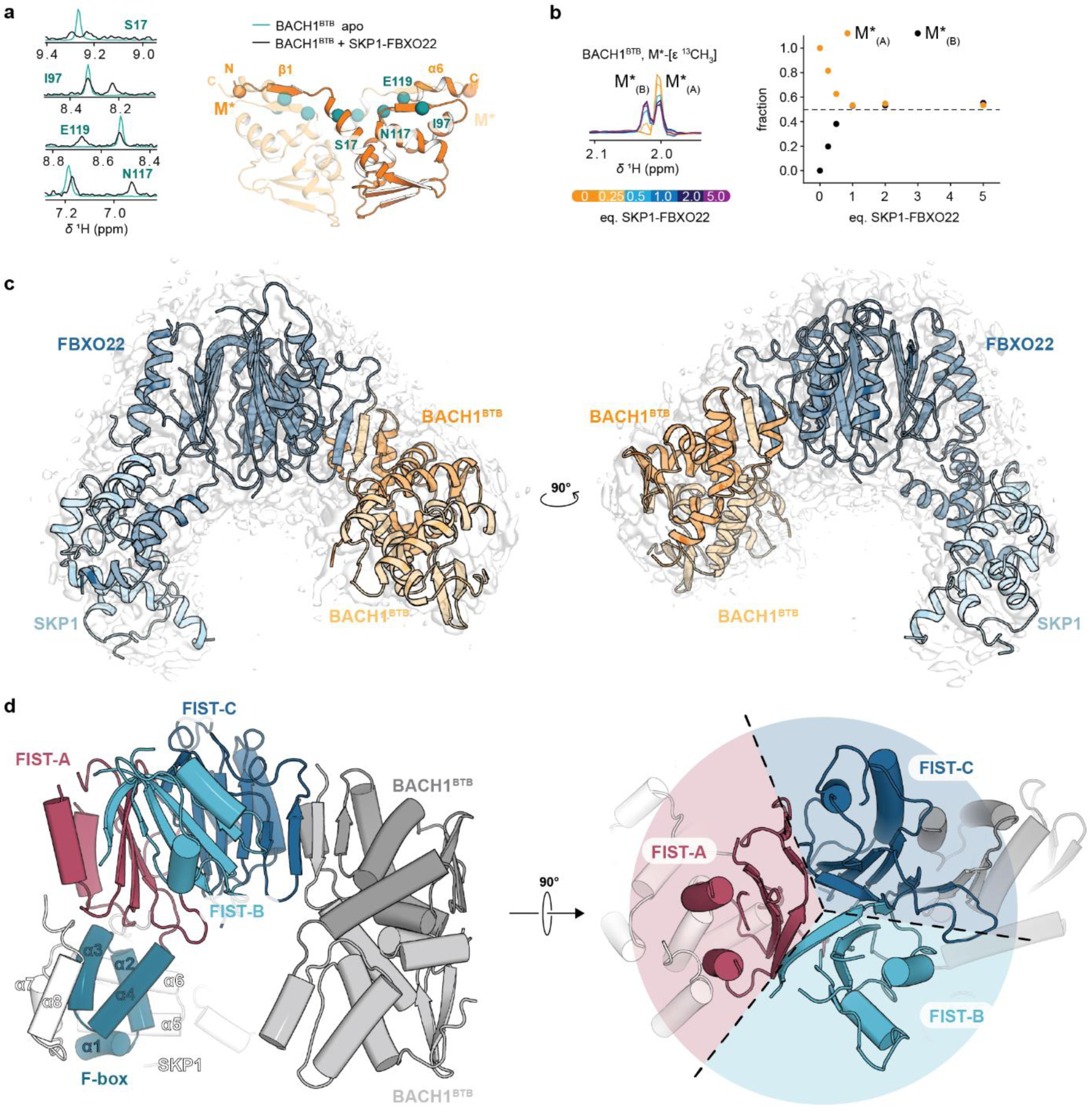
Structural analysis of the SKP1-FBXO22/BACH1^BTB^ complex by nuclear magnetic resonance spectroscopy and cryo-electron microscopy. **a** NMR analysis of the interaction between BACH1^BTB^ and SKP1-FBXO22. The NMR spectra represent 1D projections of the S17, I97, N117, E119 peaks in the (^1^H, ^15^N)-HSQC NMR spectra of U-^15^N BACH1^BTB^ (see supplementary figure 5b) in the absence (cyan) and presence of 5 equivalents SKP1-FBXO22 (black, ten-fold amplified signal). The peak splitting suggests a symmetry break in the BTB domain upon FBXO22 binding. The positions of S17, I97, N117, E119 in the BTB domain are highlighted as cyan spheres in an Alphafold2 structure of the BACH1 BTB domain. **b** 1D projection of the Met-[ε ^13^CH_3_] peak of the artificial methionine residue (M*) in the N-terminus of U- (^2^H, ^15^N), (ILVMA)-^13^CH_3_ labeled BACH1^BTB^ (see supplementary figure 5e) splitting into M*_(A)_ and M*_(B)_ in the presence of increasing equivalents of SKP1-FBXO22. The position of M* in the BACH1 BTB domain is highlighted as an orange sphere in the structure in (**a**). The plot of the peak fractions of M*_(A)_ and M*_(B)_ against the equivalents of SKP1-FBXO22 is shown on the right. **c** Cryo-EM density map superimposed onto the structural model of the SKP1-FBXO22/BACH1^BTB^ complex. **d** Domain architecture of FBXO22. The interaction with SKP1 is mediated by the N-terminal F-box domain while the interaction with BACH1^BTB^ occurs through the C-terminal FIST domain^76^. The FIST domain can be divided into three subdomains, FIST-A, FIST-B, and FIST-C which adopt a threefold pseudosymmetry. The interaction with BACH1^BTB^ occurs through the FIST-C subunit, in which the loop folds into a β-sheet.

To gain high-resolution insights into the BACH1-FBXO22 interaction, we determined a 3.2 Å single particle cryo-electron microscopy (cryo-EM) structure of SKP1-FBXO22 in complex with BACH1^BTB^ (Figure 2c, Supplementary Figure 6 and 7). Despite several surface exposed loops (Supplementary Figure 8a and b), we could confidently model SKP1-FBXO22 and BACH1^BTB^ into the EM density map allowing for a detailed analysis of the SKP1-FBOX22 architecture as well as the FBXO22/BACH1^BTB^ interface and binding mechanism. FBXO22 interacts through its N-terminal F-box domain with SKP1, as expected for SCF-type E3 ligases (Figure 2d)^14–17^, while the F-box and intracellular signal transduction (FIST) domain engages with the BACH1^BTB^ dimer. The low resolution in the periphery of SKP1 as well as in parts of BACH1^BTB^ indicates substantial protein flexibility in those regions (Supplementary Figure 8).

In line with our NMR analysis (Figure 2a and b), FBXO22 binds the BACH1^BTB^ dimer in an asymmetric manner through an interface containing the degron in β1 of one subunit and helix α6 of the other subunit (Figure 2c). Notably, the FIST domain adopts a threefold pseudo symmetric fold consisting of three subdomains termed FIST-A, FIST-B, and FIST-C hereafter (Figure 2d, Supplementary Figure 9a and b). The core of each FIST subunit is a β-α-β-α-β-loop-β motif in which the four β-strands form an antiparallel β-sheet packed against the two α-helices. The loop is largely unresolved in FIST-A and FIST-B, while in FIST-C, it forms a β-sheet that extends the subdomain’s core β-sheet by two additional β-strands. At the surface, the loop interacts through β16 with β1 of one BACH1^BTB^ subunit in an antiparallel manner resulting in an eight-stranded β-sheet comprising β14, β13, β17, β12, β15, β16 of FBXO22, β1 of one BACH1^BTB^ subunit, and β5 of the other subunit (Figure 3a). The binding epitope in BACH1 aligns well with the before described degron sequence ^9^**F**A**Y**E**S**^13 10^ and our mutagenesis data (Figure 1h and i). We further validated β16 in FBXO22 as the BACH1 BTB binding site using point mutation R376P to disrupt secondary structure formation in β16 and point mutation R376D to invert the side chain charge. Nano-DSF, CD spectroscopy and SEC-MALS confirmed the structural integrity of the R376P and R376D mutants (Supplementary Figure 10). However, both mutants showed a substantially reduced ability to compete with the SKP1-FBXO22/BACH1^BTB^ interaction (IC_50_=1350 nM and 900 nM, respectively) compared to wildtype SKP1-FBXO22 (IC_50_=0.5 µM) in our TR-FRET assay and did not form stable complexes with BACH1^BTB^ in fSEC experiments (Figure 3b and c). This confirms that β16 in FBXO22 is an essential feature for the interaction with BACH1.

**Figure 3:**
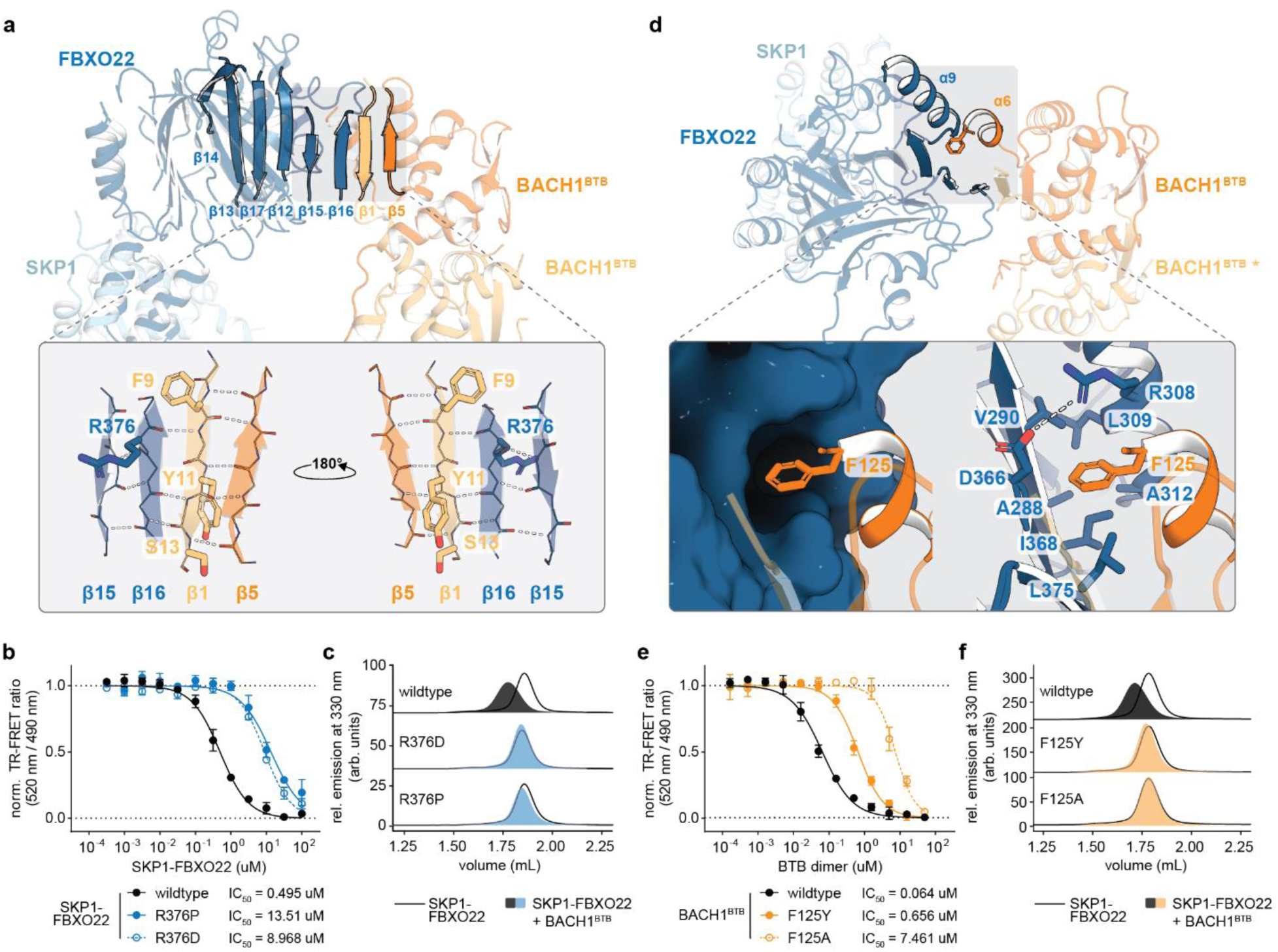
Structural features of the interaction between the FBXO22 FIST domain and the BACH1 BTB domain. **a** The BACH1 BTB domain interacts through its β1 strand with β16 in the FBXO22 FIST-C loop thus forming a seven-stranded antiparallel β-sheet involving (β14-β13-β17-β12-β15-β16)^FBXO22^-(β1-β5)^BACH1^. A close-up view of the β-sheet shows highlights the residues F9, Y11, and S13 of the FBXO22-dependent BACH1 degron and FBXO22 residue R376 at the binding interface. **b, c** Site-directed mutagenesis identifies R376 in the β16 strand of FBXO22 as a key residue in for the interaction with BACH1^BTB^. SKP1-FBXO22 R376P as well as R376D mutants exhibit a substantially reduced potency to compete with wildtype SKP1-FBXO22 for BACH1^BTB^ binding in the TR-FRET competition assay (**b**) and do not form complexes with BACH1^BTB^ in fSEC experiments (**c**). **d** In the SKP1-FBXO22/BACH1^BTB^ complex structure, β12, β15, β16 and α9 of the FIST-C domain form a hydrophobic pocket that accommodates residue F125 in the α6 helix of the BACH1 BTB domain. A close-up view shows how the F125 side chain penetrates the pocket formed by several aliphatic FBXO22 side chains. **e, f** Site-directed mutagenesis indicates that a phenylalanine side chain at residue 125 in BACH1 is crucial for the interaction with FBXO22. As demonstrated by TR-FRET (**e**) and fSEC (**f**) experiments, mutagenesis of F125 to alanine or tyrosine substantially perturbs the interaction of BACH1^BTB^ with FBXO22.

Beyond the intermolecular β-sheet between FBXO22 and BACH1^BTB^, the C-terminal helix α6 of one BACH1^BTB^ subunit forms extensive contacts with α9, β12, β15, and β16 of the FBXO22 FIST-C subunit (Figure 3d). The side chain of BACH1 residue F125 in α6 penetrates a hydrophobic pocket formed by FBXO22 residues A288, V290, A312, I314, I368, L375 and a salt-bridge between D366 and R308. Mutating F125 in BACH1^BTB^ to either tyrosine (F125Y) or alanine (F125A) substantially perturbed complex formation with FBXO22 in the TR-FRET (IC_50_=656 nM and 7461 nM, respectively, vs. IC_50_=64 nM for wildtype BACH1^BTB^) and the fSEC assay (Figure 3e and f, Supplementary Figure 2a-d). Thus, β-sheet formation in the FBXO22 FIST-C loop is not only necessary for interaction with the BACH1 β1 strand but also for forming a hydrophobic pocket with the FIST-C α6 helix for the F125 side chain of BACH1 helix α6.

### Folding of the FBXO22 FIST-C loop mediates BACH1 recognition

Next, we asked whether the BACH1-FBXO22 interaction requires conformational changes in either of the binding partners. The comparison of the previously published BACH1^BTB^ crystal structure (PDB: 2IHC)^18^ and our cryo-EM structure of the SKP1-FBXO22/BACH1^BTB^ complex suggests that the BACH1 BTB domain undergoes only very subtle structural changes upon binding to FBXO22 (RMSD=0.669 Å) (Figure 4a, b), with side chain reorientations of residues F9 in β1 and F125 in α6 being the most notable features (Figure 4b). F9 and F125 are crucial for BTB domain stability and binding to FBXO22 (Figure 1, Supplementary Figure 2c-e). When we probed the conformational flexibility of the BACH1 BTB domain by NMR, we observed a very rigid backbone throughout the protein on the picosecond to nanosecond timescale by {^1^H},^15^N-heteronuclear Overhauser effect (*het*NOE) NMR spectroscopy (Figure 4c), similar as previously observed for other BTB dimers^19^. Hydrogen deuterium exchange (HDX) NMR experiments suggest a very tight BTB dimer, as demonstrated by slow HDX in the dimer interface comprising α1, α2 and α5 (Figure 4d). However, we observed comparably fast HDX in β1 and α6 indicating protein motions on the microsecond to millisecond timescale within this part of the protein. These data suggest that the BACH1 BTB domain overall forms a rigid and tight dimer but exhibits flexibility on slow time scales in β1 and α6 to enable subtle structural changes necessary for the binding to FBXO22.

**Figure 4:**
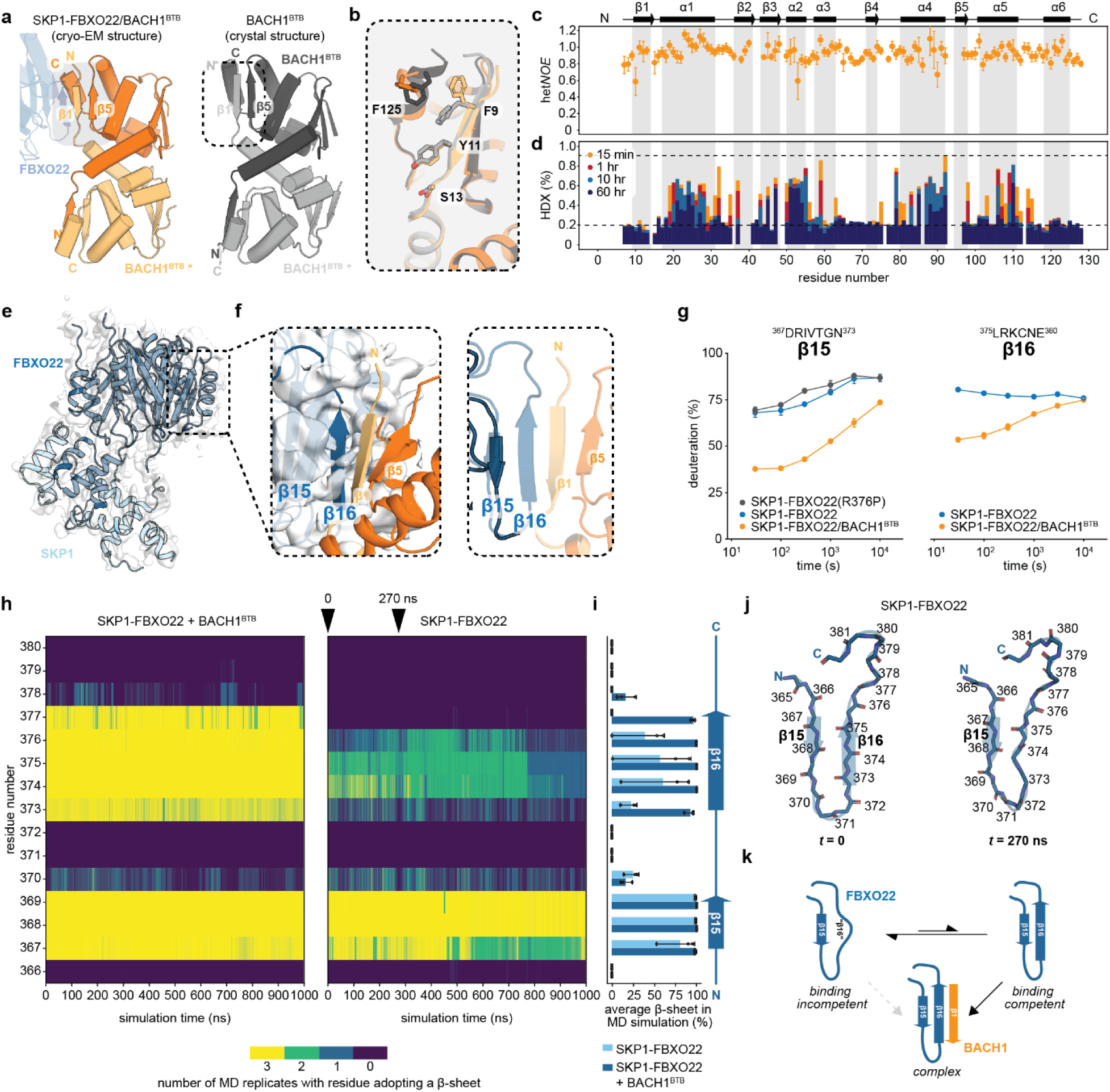
Structural dynamics underlying the FBXO22-BACH1 interaction. **a** BACH1^BTB^ conformation within the cryo-EM structure of the SKP1-FBXO22/BACH1^BTB^ complex and in the X-ray crystal structure (PDB ID: 2IHC) of the isolated BACH1 BTB domain. **b** Superposition of the BACH1 BTB domain bound to SKP1-FBXO22 in the cryo-EM structure with the BACH1 BTB domain X-ray crystal structure (PDB ID: 2IHC) suggests that only very subtle structural changes in BACH1 are required for binding to FBXO22. **c** {^1^H},^15^N-het*NOE* NMR analysis of U-^15^N labeled BACH1^BTB^ to probe fast backbone dynamics on the ns-ps time scale. Het*NOE* values close to 1 indicate a rigid protein backbone while regions exhibiting fast dynamics would be expected to show substantially decreased het*NOE* values. **d** Hydrogen deuterium exchange (HDX) NMR analysis of BACH1^BTB^ suggests that the dimerization domain containing the FBXO22-dependent degron exhibits substantial flexibility on the µs-ms time scale. Maximum deuteration occurs at a rel. HDX value of 0.2 while no deuteration is indicated by a rel. HDX value of 0.9. **e** Cryo-EM density map superimposed onto the structural model of the SKP1-FBXO22 heterodimer. **f** Close up view of the BACH1 binding site in the SKP1-FBXO22 cryo-EM structure highlights that β16 is not resolved, presumably due to protein dynamics, while it forms an intermolecular β-sheet with BACH1 upon complex formation. **g** Hydrogen deuterium exchange (HDX) MS deuterium uptake charts for peptides corresponding to β15 (^267^DRIVTGN^373^) and β16 (^375^LRKCNE^380^) in the BACH1 binding site in FBXO22 obtained from HDX analysis of SKP1-FBXO22 wildtype, SKP1-FBXO22(R376P), and SKP1-FBXO22/BACH1^BTB^. **h, I, j** Evolution (**h**) and average (**i**) β-sheet content for β15 and β16 in the SKP1-FBO22/BACH1^BTB^ as well as in SKP1-FBXO22 over the time of a 1 µs all-atom molecular dynamics simulation. Representative conformations of β15 and β16 at the start of the simulation, t=0, and after 270 ns showing the unfolding of the β-sheet are shown in (**j**). **k** Conformational selection mechanism for the FBXO22-BACH1 interaction. In the absence of BACH1, the FIST-C loop in FBXO22 is in equilibrium between an unfolded, binding incompetent state and a binding competent state in which a β-sheet forms between β15 and β16. BACH1 selects the FBXO22 subpopulation with a preformed β-sheet, thereby gradually shifting the equilibrium of FBXO22 to the BACH1 bound state.

To identify potential conformational changes in FBXO22 underlying the recognition of the BACH1 BTB dimer, we determined a 3.4 Å cryo-EM structure of SKP1-FBXO22 alone (Figure 4e, Supplementary figure 11a). Overall, the SKP1-FBXO22 apo state aligns well with the BACH1^BTB^ bound state (RMSD=0.813 Å). Strikingly, no EM density was observed for the β16 strand forming the BACH1^BTB^ binding site in the SKP1-FBXO22/BACH1^BTB^ complex (Figure 4f). Interestingly, we observed that the mutations R376D and R376P in β16 neither affected the overall secondary structure nor the thermal stability of the protein further corroborated a lack stable secondary structure in this region in the apo state (Supplementary figure 10a-d). Moreover, AlphaFold2 predicts the β15 and β16 region in the FIST-C subdomain of SKP1-FBXO22 alone with a comparably low confidence and high structural variability (Supplementary figure 11b), which may further suggest that this region does not adopt a stable structure^20^.

To further assess flexibility of this structural element, we probed the conformational dynamics of the β15 and β16 region in the FBXO22 FIST-C domain by hydrogen deuterium exchange mass spectrometry (HDX-MS) and all-atom molecular dynamics (MD) simulations (Figure 4g-j). In the apo state, β15 and β16 exhibited rapid hydrogen deuterium exchange (HDX) suggesting an unstable β-sheet in the absence of BACH1^BTB^ in the FIST-C loop (Figure 4g). Interestingly, secondary structure perturbation through the R376P mutation in β16 further increased the HDX rate in the loop by ∼5% which may indicate residual secondary structure content in this region in the wildtype protein (Figure 4g). In the presence of BACH1^BTB^, HDX was substantially reduced in the FIST-C loop consistent with the formation of a stable β-sheet in β15 and β16. Likewise, we observed that in the SKP1-FBXO22/BACH1^BTB^ complex, β15 and β16 formed a stable β-sheet over the time of a 1 µs all-atom MD simulation (Figure 4h). By contrast, after removal of the BACH1^BTB^ domain, β16 started to unfold and was disordered for 50% of the time during the 1 µs simulation. This effect was reduced in β15 which largely retained its β-strand conformation during the simulation (Figure 4i).

In conclusion, the cryo-EM, HDX-MS and MD simulation data suggest that the FIST-C loop in FBXO22 is responsible for BACH1 binding. The loop predominantly adopts an unstructured state incompatible with BTB binding, which is in equilibrium with a binding competent β-sheet consisting of β15 and β16. This observation is compatible with a conformational selection mechanism in which BACH1 requires β-sheet formation in the FBXO22 FIST-C loop prior to binding (Figure 4j).

### SCF^FBXO22^ ubiquitylates the BACH1 C-terminus independent of heme

Having established the structural basis for BACH1 binding to FBXO22 allowed us to further explore the molecular details underlying the ubiquitylation of BACH1 by SCF^FBXO22^. For this, we reconstituted the neddylated SCF^FBXO22^ complex to monitor modification of full-length BACH1 with fluorescently labeled ubiquitin in an in vitro ubiquitylation assay. A side-by-side comparison of wildtype ubiquitin (Ub) and a lysine-deficient ubiquitin mutant (Ub^0^) in our assay showed that BACH1 gets modified multiple times with both Ub and Ub^0^ (Figure 5a). This suggests that BACH1 is ubiquitylated by SCF^FBXO22^ at multiple sites and is not restricted to one specific lysine. Time course experiments indicated a relatively slow ubiquitylation of BACH1 by SCF^FBXO22^. This was demonstrated by the negligible ubiquitin consumption and the stable levels of non-ubiquitylated BACH1 over the time course of the experiment. A comparison of wildtype FBXO22 with the FBXO22 R376P mutant in the time course experiment confirmed that the loss of binding to BACH1 results in reduced ubiquitylation in vitro (Figure 5b).

**Figure 5:**
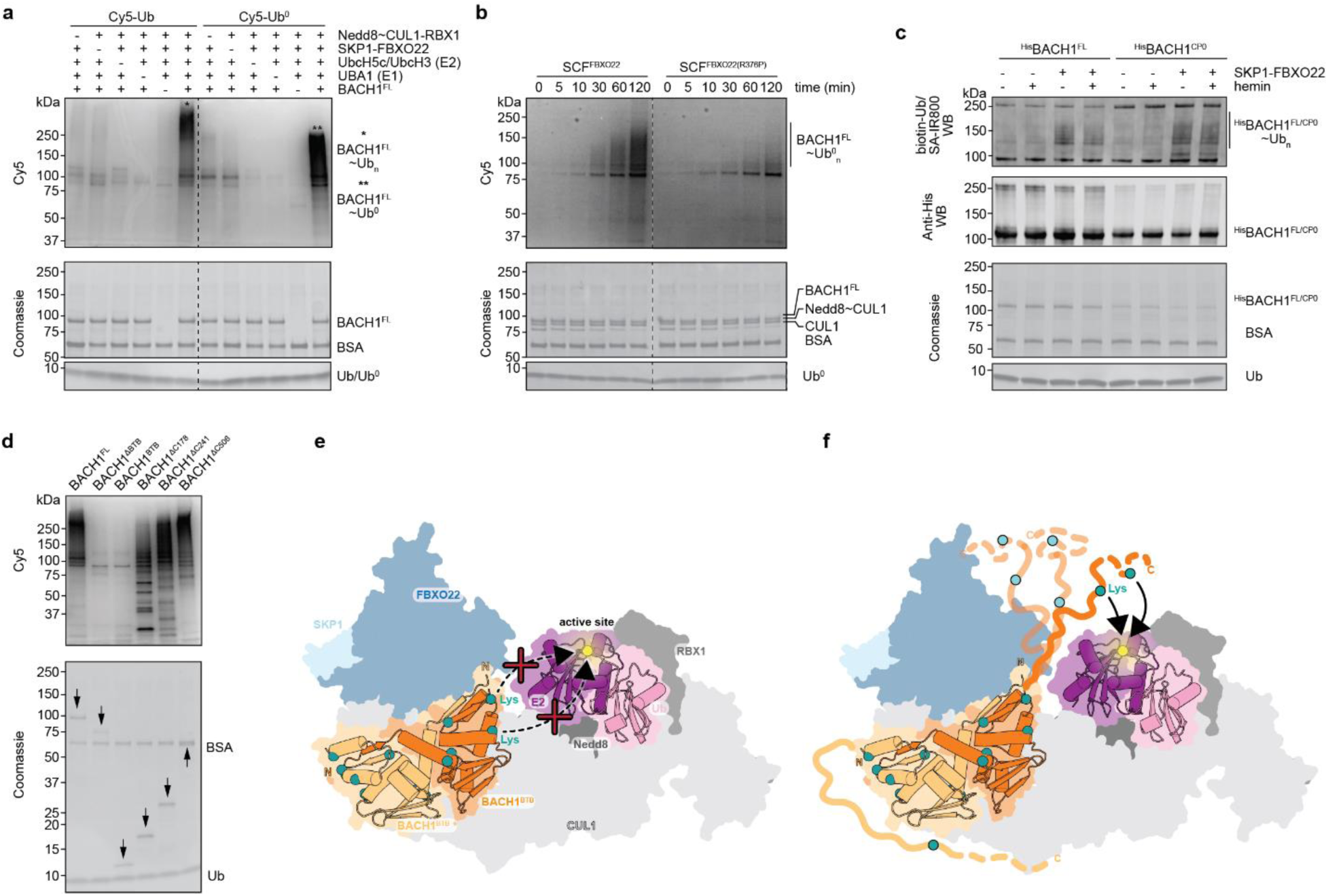
In vitro ubiquitylation of BACH1 by SCF^FBXO22^ in the disordered C-terminus. **a** Comparison of BACH1 ubiquitylation by SCF^FBXO22^ using Cy5-Ub and Cy5-Ub^0^. The upper panel shows the fluorescence scan, and the lower panel shows the Coomassie stained SDS-PAGE gel. The detection of ubiquitylated BACH1 at molecular weights >120 kDa with either Ub or Ub^0^ suggests that BACH1 is ubiquitylated at multiple sites. **b** Time course experiment comparing the ubiquitylation of BACH1 by wildtype SCF^FBXO22^ and an FBXO22 R376P mutant. **c** In vitro ubiquitylation assay of BACH1 in dependence of hemin. Due to the quenching properties of hemin, ubiquitylation was monitored using biotinylated-Ub followed by WB using a streptavidin-IR800 conjugate (upper panel). His-tagged BACH1^FL^ or BACH1^CP0^ were visualized by Anti-His WB (central panel) or Coomassie staining (lower panel). **d, e** Structural model explaining why the BTB domain is not ubiquitylated by SCF^FBXO22^. The model was generated by superimposing the cryo-EM structure of SKP1-FBXO22/BACH1^BTB^ along SKP1 onto the structure of a SCF^β-TRCP1^ trapped in a substrate ligation state (PDB: 6TTU)^22^. **e**: The model illustrates that lysine residues in the BACH1 BTB domain are too far away from the Ub∼E2 active site for efficient ubiquitin transfer. **f**: The model shows how the disordered BACH1 region extending from the BTB domain C-terminus can bring lysine residues to the Ub∼E2 active site through conformational sampling. **f** Identification of the BACH1 C-terminus as the ubiquitylation site using BACH1 deletion constructs. The upper panel shows the fluorescence scan, and the lower panel shows the Coomassie stained SDS-PAGE gel. Arrows indicate the bands of the respective BACH1 construct.

Despite the observation that heme did not influence the BACH1-FBXO22 interaction (Figure 1f and g), heme may modulate BACH1 ubiquitylation by other means, e.g. through structural reorganization of BACH1 around the heme binding sites to reposition lysine residues for a more efficient ubiquitin transfer. To test this, we monitored the ubiquitylation of BACH1^FL^ as well as BACH1^CP0^ by SCF^FBXO22^ in the absence and presence of hemin. Strikingly, there was no effect of hemin on the ubiquitylation of BACH1 (Figure 5c). This stands in contrast to what was described in a recent preprint^21^.

To rationalize how BACH1 is ubiquitylated by SCF^FBXO22^, we generated a structural model of the BACH1 BTB domain bound to SCF^FBXO22^ loaded with a Ub-charged E2 enzyme (Ub∼E2) based on a published cryo-EM structure of a SCF-type E3 ligase complex trapped in a substrate ligation step (PDB: 6TTU) (Figure 5e)^22^. The model suggests a significant distance between the BTB domain bound to SKP1-FBXO22 at the CUL1 N-terminus and the Ub∼E2 bound to NEDD8 and RBX1 at the CUL1 C-terminus. We thus questioned whether the BTB domain is the actual site where BACH1 is ubiquitylated by SCF^FBXO22^. In fact, it is a common theme that ubiquitylation sites reside in disordered regions often >20 residues distant from the degrons mediating the interaction with an E3 ligase^23^. Indeed, the structural model suggests that the extension by the disordered region downstream of the BTB domain may bring lysine residues close to the E2∼Ub active site and enable ubiquitin transfer (Figure 5f). We thus speculated that the disordered region C-terminal to the BTB domain contains the ubiquitylation sites instead of the BTB domain itself.

To test this hypothesis, we tested a panel of BACH1 mutants in which the C-terminus was gradually deleted in our in vitro ubiquitylation assay (Figure 5d). As a control, we used the FBXO22-binding deficient BACH1^ΔBTB^ mutant. SEC-MALS and fSEC experiments confirmed the structural integrity of the proteins and their ability to interact with SKP1-FBXO22 (Supplementary Figure 12). BACH1^FL^ was readily ubiquitylated, while no BACH1^ΔBTB^ ubiquitylation was observed as expected due to the lack of binding affinity for FBXO22. Strikingly, also BACH1^BTB^ was not ubiquitylated (Figure 1f). However, adding ∼50 C-terminal residues to the BACH1 BTB domain, termed BACH1^ΔC178^ (BACH1 residues 7-177), yielded a construct that was substantially ubiquitylated by SCF^FBXO22^. Ubiquitylation was enhanced by further extending the BACH1 C-terminus to residues 240 and 505 yielding the constructs BACH1^ΔC241^ and BACH1^ΔC506^, respectively (Figure 5e). This confirms that the BACH1 is ubiquitylated in the disordered region downstream of the BTB domain.

In conclusion, our data show that the ubiquitylation of BACH1 by SCF^FBXO22^ is not affected by hemin binding in the disordered BACH1 C-terminus even though this region contains the ubiquitylation sites.

### Cancer-associated mutations in FBXO22 modulate BACH1 ubiquitylation

BACH1 is an emerging drug target for cancer therapies^24^. However, its pathological functions seem multifaceted and cancer-type dependent suggesting that upregulation as well as downregulation of BACH1 may contribute to disease progression^25^. Considering that BACH1 levels are directly regulated by the ubiquitin proteasome system (UPS) through SCF^FBXO22^, and that FBXO22 knockout has recently been shown to stabilize BACH1 in lung cancer cells^10^, we hypothesized that cancer cells may develop genetic alterations that modulate the interaction between FBXO22 and BACH1. We identified the mutations Q307E, Q307R, R367W, R367L in the FBXO22 FIST-C subdomain in the TCGA database^26^ that we deemed capable of modulating the interaction with BACH1 (Figure 6a). In the SKP1-FBXO22/BACH1^BTB^ complex, Q307 in helix α9 is proximal to BACH1 residue Q124. Interestingly, Q124 of BACH1 is neighboring F125, a key residue for FBXO22 binding. R367 is part of the β15 strand in the FIST-C loop close to R376 in β16, another key residue for BACH1 binding. Therefore, mutation of the arginine side chain to tryptophan or leucine may sufficiently perturb the local β-sheet structure and abrogate BACH1 binding, similar to the R367P and R376D mutations.

**Figure 6:**
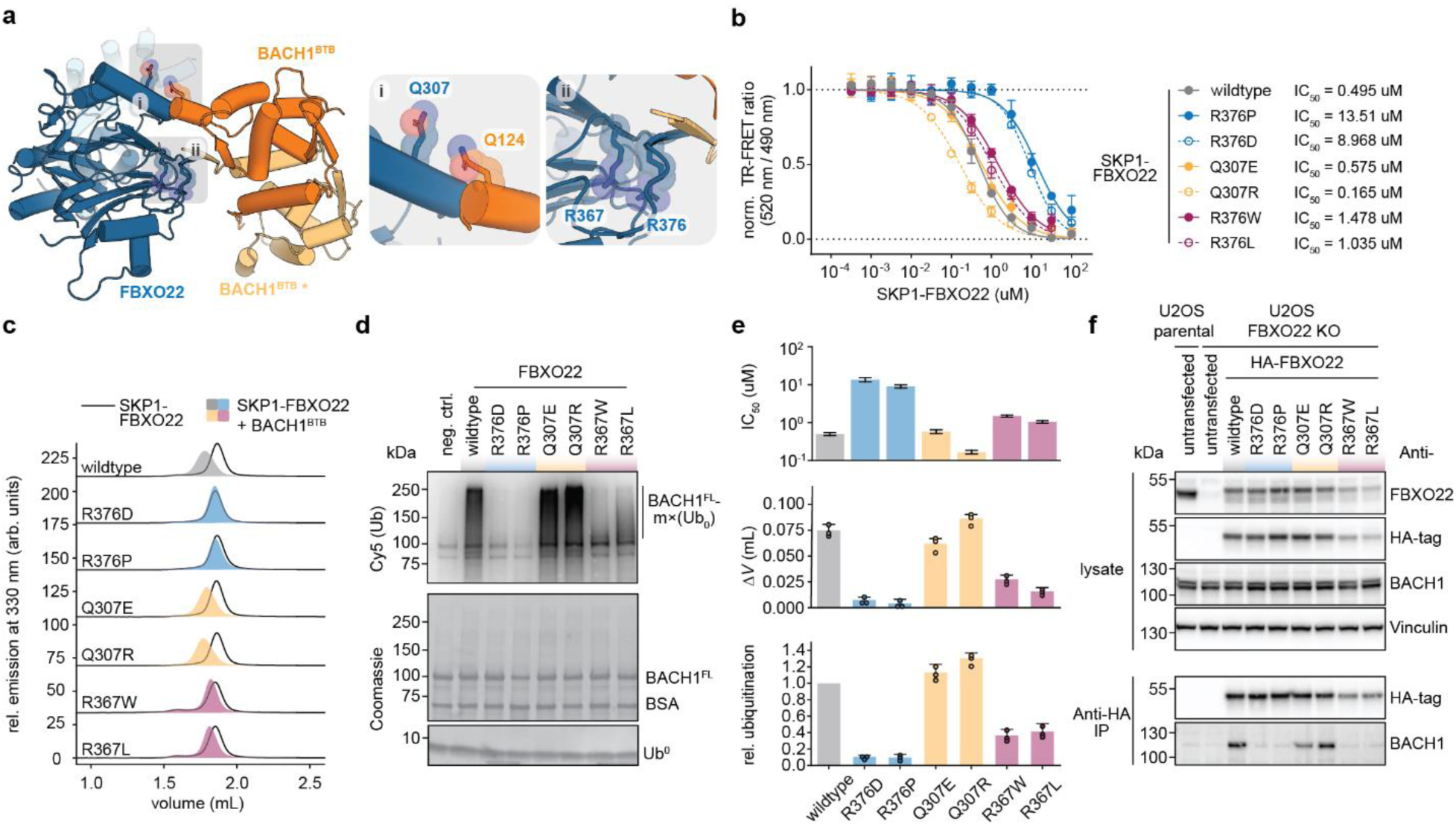
Impact of cancer-associated mutations in FBXO22 on the binding and ubiquitylation of BACH1. **a** Location of cancer-associated mutations in FBXO22 reported in the TCGA mapped in the cryo-EM structure of the SKP1-FBXO22/BACH1^BTB^ complex: (i) FBXO22 Q307E and Q307R, (ii) FBXO22 R367W and R367L. **b, c** Evaluation of FBXO22 mutations on the interaction with BACH1^BTB^ in the TR-FRET competition assay (**b**) and the fSEC assay (**c**). The FBXO22 mutants R376D and R376P were included as negative controls. **d** In vitro ubiquitylation of BACH1 by SCF^FBXO22^ wildtype and mutants. Top panel shows the fluorescence scan and the lower panel the Coomassie staining of the SDS-PAGE gel. **e** The correlation between the TR-FRET, fSEC and in vitro ubiquitylation assays confirms that modulating the BACH1 interaction through FBXO22 mutations correlates with altered BACH1 ubiquitylation. **f** Co-immunoprecipitation experiments of FBXO22 wildtype and mutants with BACH1. HA-tagged FBXO22 constructs were transiently expressed in U2OS FBXO22 knockout cells and tested for their interaction with BACH1 by HA-pulldown followed by western blot detection. Untransfected parental and FBXO22 KO U2OS cells were used as controls.

To investigate the impact of these mutations on the ubiquitylation of BACH1, we probed the impact of these cancer-associated mutations on the interaction of BACH1^BTB^ with SKP1-FBXO22. Importantly, the mutations did not perturb the overall structure of SKP1-FBXO22 as evidenced by SEC-MALS and circular dichroism (CD) spectroscopy (Supplementary Figure 10a-e). While the Q307E and Q307R mutations had no effect on the thermal stability of SKP1-FBXO22, similar to the R376P and R376D mutations, the R367W and R367L mutations substantially destabilized the protein (Supplementary Figure 10b-d), supporting the notion that β15 is a more stable secondary structure than β16 (Figure 4). In our TR-FRET competition assay, the Q307E mutant behaved similar as the wildtype protein (IC_50_=575 nM for Q307E versus IC_50_=495 nM for wildtype) (Figure 6b). Unexpectedly, the Q307R mutant exhibited competition at lower concentrations than wildtype (IC_50_=165 nM for Q307E), indicating that the Q307R mutation increases the affinity for BACH1^BTB^, potentially due to the formation of additional interactions between BACH1 residue Q124 and the mutated residue R307 (Figure 6b). This preserved ability of the SKP1-FBXO2(Q307E/R) mutants to form complexes with BACH1^BTB^ was also observable in fSEC experiments (Figure 6c). By contrast, we found the SKP1-FBXO22(R367W/L) mutants to compromise binding of wildtype BACH1^BTB^. In the TR-FRET assay, the potency to compete with wildtype SKP1-FBXO22 for BACH1^BTB^ binding was substantially reduced (IC_50_=1478 nM for R367W versus IC_50_=1035 nM for R367L) (Figure 6b). This finding was further confirmed by the impaired ability of these two mutants to form complexes with BACH1^BTB^ in fSEC experiments (Figure 6c).

Next, we asked whether these mutations in FBXO22 which affect the interaction with BACH1 are reflected in altered BACH1 ubiquitylation in vitro (Figure 6d). As expected, SCF^FBXO22^ was not able to mediate BACH1 ubiquitylation when carrying the R376P or R376D mutations in FBXO22, while the Q307E and Q307R mutants showed robust ubiquitylation as the wildtype FBXO22. In line with the increase affinity for BACH1^BTB^ observed in our TR-FRET assay (Figure 6b), the Q307R mutant also showed a stronger ubiquitylation of BACH1 than wildtype FBXO22. By contrast, BACH1 ubiquitylation was substantially reduced for the R367W and R367L mutants, but not entirely lost. Overall, the effects of the mutations in FBXO22 on the binding to BACH1 correlated well with altered in vitro ubiquitylation of BACH1 (Figure 6e)

Finally, we tested the effects of the FBXO22 mutations on binding of full-length BACH1 in a cellular context (Figure 6d). For this, we generated a FBXO22 knockout U2OS cell line using CRISPR/Cas9 and transiently expressed HA-tagged FBXO22 wildtype or the R367P, R367D, Q307E, Q307R, R367W, and R367L mutants. Protein immunoblotting analysis confirmed the expression of all constructs at levels similar to endogenous FBXO22 in wildtype U2OS cells, except for the R367W and R367L mutants which showed lower expression, likely due to the impaired stability. Crucially, co-immunoprecipitation experiments could largely reproduce our observations with recombinant proteins. In fact, while BACH1 co-immunoprecipitated at similar levels with wildtype FBXO22 and the Q307E and Q307R mutants, co-immunoprecipitation with FBXO22 mutants R376P, R376D, R367W, and R367L was severely impaired. This highlights how these residues are relevant for the interaction between FBXO22 and BACH1 in the cellular context.

In summary, our in vitro and cellular binding data, together with the concordant in vitro ubiquitylation data reveal that cancer-associated mutations in FBXO22 directly affect levels of BACH1 ubiquitylation.

### BTB domain destabilization redirects BACH1 from FBXO22 to FBXL17

Other E3 ligases have been linked to the degradation of BACH1 under oxidative stress, among them SCF^FBXL17 27^. Unlike FBXO22, FBXL17 binds monomeric BTB domains^13, 14^ and thus requires BTB dimer destabilization (Figure 7a). We thus speculated that FBXO22 is a more potent SCF receptor module for recognizing BACH1 than FBXL17. To test this, we generated recombinant SKP1-FBXL17 and probed its ability to bind the BACH1 BTB domain by NMR and fSEC (Figure 7b and c). FBXL17 showed only very weak interaction with BACH1^BTB^ compared to FBXO22 indicating a much higher affinity of BACH1 for FBXO22 than for FBXL17. Likewise, SCF^FBXO22^ more potently ubiquitylated SCF^FBXL17^ in vitro (Supplementary Figure 13a). Strikingly, we observed that destabilizing mutations in the BTB domain redirect the interaction of BACH1^BTB^ from FBXO22 to FBXL17 (Figure 7c). While the F9, Y11A and S13A mutants failed to interact with FBXO22, they readily interacted with FBXL17 in fSEC experiments. This is in line with the previously described role of FBXL17 in dimerization quality control (DQC) of BTB domains^13, 14^ and suggests that FBXL17 recognizes BACH1 upon BTB domain destabilization.

**Figure 7:**
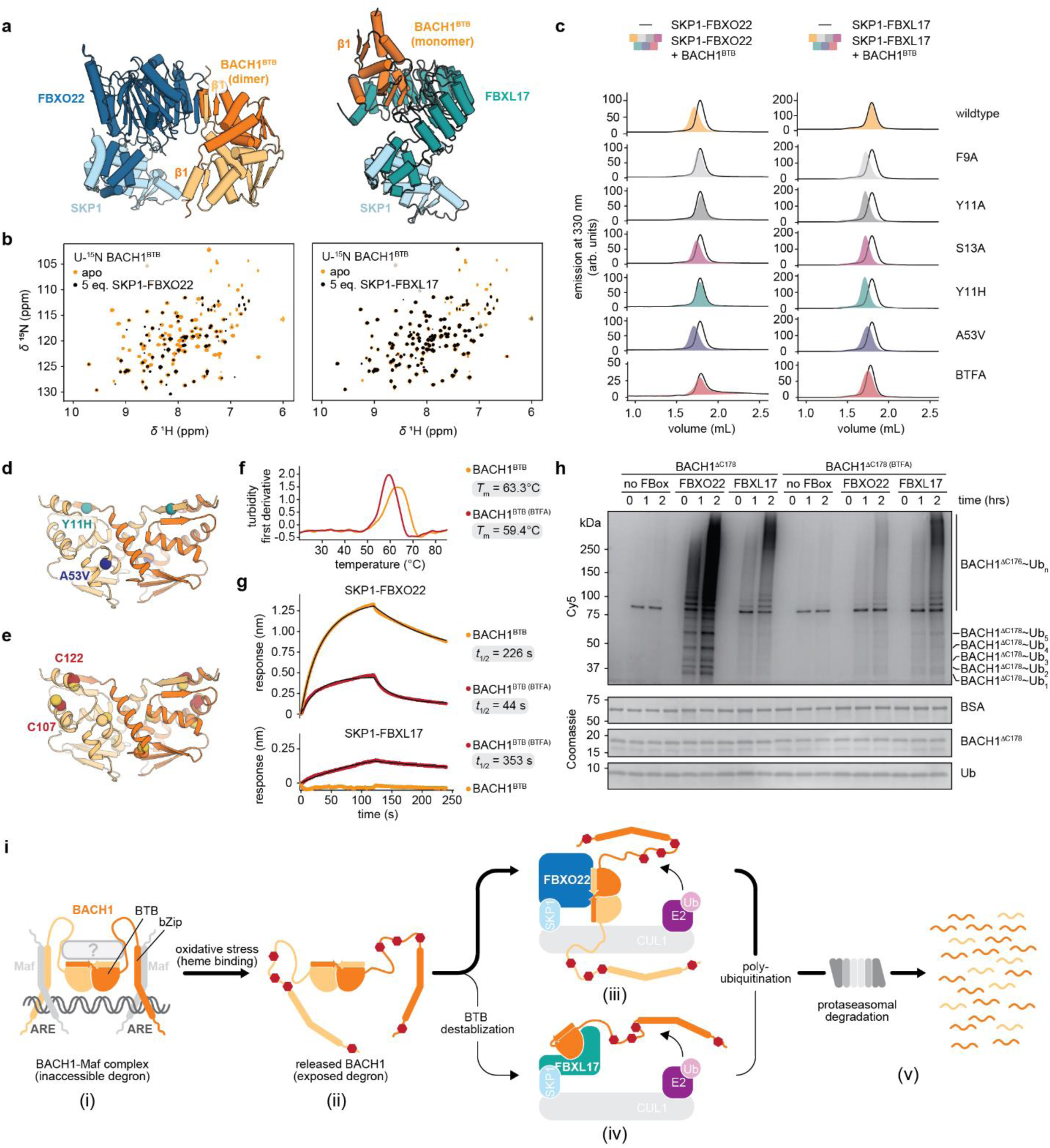
BTB destabilization redirects BACH1 binding to FBXL17. **a** Comparison of the BACH1 binding modes in the cryo-EM structure of the SKP1-FBXO22/BACH1^BTB^ complex and a homology model of the SKP1-FBXL17/BACH1^BTB^ complex. The homology model was generated using the SWISS-MODEL server based on the X-ray crystal structure of the SKP1-FBXO22/KEAP1^BTB(F64A,S172A)^ complex (PDB: 6W66)^14, 77, 78^. **b** Overlayed (^1^H,^15^N)-TROSY NMR spectra of (U-^15^N)-BACH1^BTB^ in the apo state (orange) or in the presence of 5-fold molar excess of SKP1-FBXO22 or SKP1-FBXL17 (black). **c** Complex formation of wildtype, mutant, and cysteine-modified BACH1^BTB^ constructs with SKP1-FBXO22 (left) or SKP1-FBXL17 (right) monitored by fSEC. **d, e** Cancer associated mutants Y11H and A53V (**d**) and reactive cysteine residues C107 and C122 (**e**) mapped on to the structure of the BACH1 BTB domain (AlphaFold2 model). **f** Cysteine-modification at C107 and C122 destabilizes the BACH1^BTB^ domain by ∼4°C as assessed by nano-DSF. **g** Quantitative assessment of the binding kinetics of the interaction between SKP1-FBXO22 or SKP1-FBXL17 and unmodified as well as modified BACH1^BTB^. Complex lifetimes *t*_1/2_ were determined from the inverse dissociation rate (see Supplementary table 2). **h** In vitro assay showing the effect of cysteine-modification in the BACH1^BTB^ domain by BTFA on the ubiquitylation of BACH1^ΔC178^ by SCF^FBXO22^ or SCF^FBXL17^. **i** Mechanism for heme-induced ubiquitylation of BACH1 by SCF^FBXO22^ and SCF^FBXL17^ leading to degradation by the proteasome. In the context of BACH1-Maf complexes bound to ARE sites (i), the degron in the BTB domain is sterically shielded, either conformationally or through a binding partner (grey box). Heme-binding under oxidative stress disrupts the complex, releases BACH1, and exposes the degron in the BTB domain (ii). While BACH1 with a dimeric BTB domain is recognized by SCF^FBXO22^ (iii), BACH1 with a destabilized BTB domain, e.g. through cysteine nitrosylation, is redirected to SCF^FBXL17^ (iv). This dual regulation by complementary ligases ensures efficient degradation of BACH1 under oxidative conditions via the proteaseome (v).

Recently, a Y11H mutation described in TCGA was shown to stabilize BACH1 in cancer cells under oxidative stress similar as the Y11F mutation (Figure 7d)^10^. Mutating the same residue to phenylalanine (Y11F) was sufficient to disrupt the BACH1-FBXO22 interaction, stabilize BACH1 in lung cancer cells, and was shown to induce metastasis in a xenograft model^10^. To assess the impact of the Y11H mutation on the BACH1-FBXO22 and BACH1-FBXL17 interaction, we expressed and purified a BACH1^BTB^ Y11H mutant (Supplementary Figure 2a). We further included the cancer associated BACH1^BTB^ A53V mutant with a mutation in the dimer interface of the BTB domain (Figure 7d). Equivalent mutations in the KEAP1 BTB domain were shown to interact strongly with FBXL17^14^. The proteins were still folded and able to form a dimer as demonstrated by SEC-MALS (Supplementary Figure 2a). However, the thermal stability of the Y11H mutant was severely reduced compared to the wildtype protein (Δ*T*_m_∼10°C), similar to the Y11A mutant, while the A53V mutant was as stable as the wildtype (Supplementary Figure 2b-d). While the BACH1^BTB^ Y11H mutant did not form a complex with SKP1-FBXO22 in fSEC experiments, it formed a stable complex with SKP1-FBXL17 (Figure 7b). By contrast, the A53V mutant showed a similar interaction with SKP1-FBXO22 as the wildtype BTB domain, but also formed a complex with SKP1-FBXL17.

Most notably, we found that cysteine modifications destabilized BACH1^BTB^ and induced a ligase shift, similarly to what we observed for genetic perturbations. In an effort to label native cysteines in BACH1^BTB^ with Bromo-1,1,1-trifluoroacetone (BTFA) for ^19^F NMR spectroscopy studies (Supplementary Figure 13b and c, we noticed loss of binding of modified BACH1^BTB^ to SKP1-FBXO22 in fSEC experiments. By contrast, complex formation with SKP1-FBXL17 was observed (Figure 7b). Importantly, modification with BTFA decreased the thermal stability of the BTB domain but retained its dimeric state (Supplementary Figure 13d-f). Mass spectrometry paired with NMR spectroscopy identified C107 and C122 in BACH1^BTB^ as the cysteines that were subject to modification by BTFA (Supplementary Figure 13d and e), with both residues being close to the FBXO22 degron (Figure 7f, Supplementary Figure 13f). We used BLI for a qualitative analysis of complex stability of modified and unmodified BACH1^BTB^ with SKP1-FBXO22 and SKP1-FBXL17 (Figure 7f). Strikingly, the cysteine modifications decreased the complex lifetime of SKP1-FBXO22/BACH1^BTB^ by one order of magnitude. Surprisingly, the complex lifetime of SKP1-FBXL17/BACH1^BTB^ increased from non-detectable to a range similar as for unmodified BACH1^BTB^ with SKP1-FBXO22. To test the effect of cysteine modifications on BACH1 ubiquitylation, we modified the construct BACH1^ΔC178^ with BTFA (Figure 7g). In line with our binding experiments, while unmodified BACH1 was more robustly ubiquitylated by SCF^FBXO22^, BTFA-modified BACH1 was more robustly ubiquitylated by SCF^FBXL17^.

Intriguingly, S-nitrosylating and alkylating agents have recently been described to induce the degradation of BACH1^28^. We thus tested by NMR whether C107 and C122 are modified by other alkylating or S-nitrosylating agents (Supplementary Figure 14a and b). Indeed, the spectra obtained by incubating BACH1^BTB^ with iodoacetamide (IAA) or S-nitrosylglutathione (GSNO) strongly resembled that of the BTFA modified BACH1^BTB^ and suggest that C107 and C122 represent reactive sites for alkylation or nitroslyation. The strong similarity between the NMR spectra of BTFA-modified BACH1^BTB^ and S-nitrosylated BACH1^BTB^ suggests that BTFA modification of cysteines is a suitable mimic of cysteine S-nitrosylation. It is thus conceivable that S-nitrosylation under oxidative stress or alkylation by antioxidants induces BACH1 degradation through ubiquitylation in a SCF^FBXL17^ dependent manner.

Overall, these data reveal that the recognition of BACH1 either as a dimer by FBXO22 or as a monomer by FBXL17 is determined by the stability of the BACH1 BTB domain. We find that the BACH1 BTB domain can be destabilized through mutations in cancer or through cysteine modifications. This destabilization induces a ligase switch in which BACH1 is redirected from FBXO22 to FBXL17.

## Discussion

In this study, we describe the molecular mechanism by which the E3 ligase SCF^FBXO22^ recognizes and ubiquitylates BACH1. FBXO22 recognizes BACH1 with high affinity through a three-dimensional degron formed by the BTB dimer interface. Our mutagenesis studies exhibit a very steep structure activity relationship of the FBXO22-dependent degron in BACH1. Consequently, other BTB proteins with low sequence similarity to BACH1 in the degron site are not recognized by FBXO22. This highlights FBXO22 as a specific receptor of the BACH1 BTB domain, in contrast to FBXL17 which acts as a universal receptor of BTB domains in the context of DQC^13, 14^.

The biophysical data and the cryo-EM structure presented here reveal that the interaction between FBXO22 and BACH1 is mediated through an extensive intermolecular β-sheet involving the FIST-C loop of FBXO22. In the absence of BACH1, this loop fluctuates between a disordered and a folded state, of which only the latter can bind the BACH1 BTB domain. Such conformational selection mechanisms have often been found at key regulatory signal transduction nodes^29, 30^. In the case of FBXO22, the loop potentially acts as a switch that kinetically controls the interaction with BACH1. Given that ubiquitylation occurs in the disordered region of BACH1, the FBXO22-BACH1 interaction requires a sufficiently slow dissociation rate to sample conformers of BACH1 in the SCF^FBXO22^/Ub∼E2/BACH1 complex which are compatible with ubiquitin transfer. The structural switch in the FIST-C loop may reduce the association rate to compensate the slow dissociation rate and maintain a nanomolar binding affinity of the FBXO22-BACH1 interaction. A fast association would increase the affinity and might lead to promiscuous BACH1 ubiquitylation and subsequent proteasomal degradation.

The high binding affinity between BACH1 and FBXO22 raises questions about the mechanism regulating their interaction. While in cells, the BACH1-FBXO22 interaction can be induced by the treatment with hemin^7, 10, 11^, we did not observe any effect of hemin on the binding kinetics and affinity of the FBXO22-BACH1 interaction or on BACH1 ubiquitylation using purified proteins. This argues against the hypothesis that heme may act as a molecular glue for the FBXO22-BACH1 interaction as described for other natural compounds or cofactors that induce ligase-substrate interactions^31, 32^. It is generally questionable whether further increasing the intrinsically high affinity between BACH1 and FBXO22, for example through post-translational modifications (PTMs) as described for other FBXO22 substrates^33^, can augment BACH1 degradation. One possibility is that heme may indirectly induce BACH1 degradation by releasing a factor that prevents the BACH1-FBXO22 interaction under normal physiological conditions. Heme was shown to displace BACH1 from ARE sites and induce BACH1 nuclear export^7, 8^. In this model, the BACH1 degron may be masked through interactions with other transcriptional co-factors such as small musculoaponeurotic fibrosarcoma (Maf) proteins or yet unknown factors when bound at AREs. Disruption of BACH1-Maf complexes and release from DNA may expose the BTB domain to enable binding of FBXO22 (Figure 7i). Elucidating which factors protect BACH1 from binding to FBXO22 and how this mechanism can be modulated may open opportunities for inducing the targeted degradation of BACH1 by small molecules for therapeutic purposes.

Recent studies have indicated that BACH1 stabilization in lung cancer cells stimulates a BACH1-dependent prometastatic gene program^10, 34^. However, several cancer types also show BACH1 downregulation^25^, potentially to activate NRF2 signaling^5^. We therefore hypothesized that cancer cells may acquire mutations in FBXO22 that affect the interaction with BACH1 and by this modulate BACH1 levels. Indeed, we identified two FBXO22 mutations in the TCGA database, R367W and R367L, which reduced the interaction with BACH1 and reduced its ubiquitylation. Interestingly, we also revealed that the cancer-related Q307R mutation in FBXO22 increased the interaction with BACH1 and caused stronger ubiquitylation. These examples suggest that cancer cells can acquire mutations that contribute to either the stabilization or the degradation of BACH1.

We found that genetic destabilization of the BACH1 BTB domain, including cancer-associated mutations, impaired the binding of FBXO22 and redirected BACH1 to FBXL17. This may indicate that in cancer cells SCF^FBXL17^ dominates the regulation of BACH1 instead of SCF^FBXO22^. Most notably, we discovered C107 and C122 as reactive cysteines residues that are readily modified by alkylating agents, resulting in the destabilization of the BACH1 BTB domain and the same ligase switch that we also observed upon mutagenesis. Importantly, we showed that the same cysteine residues can be S-nitrosylated. NO plays an important role in oxidative stress and S-nitrosylation of KEAP1 was shown to activate NRF2 signaling^35, 36^. Recent studies also suggest that NO-donors induce BACH1 degradation, an effect that is augmented by alkylating agents found in garlic^28^. However, the mechanism is yet unclear. Our data may suggest that C107 and C122 in the BACH1 BTB domain serve as sensors of oxidative stress, similar as in KEAP1^37^, and that S-nitrosylation or potentially oxidation by reactive oxygen species at these cysteines under oxidative stress induces BACH1 degradation via SCF^FBXL17^, resulting in the activation of NRF2 signaling. Notably, these cysteine residues may present suitable sites for covalent engagement by electrophilic small molecules to destabilize BACH1 and induce degradation by FBXL17 as a potential therapeutic approach.

Our findings highlight that two complementary E3 ligases ubiquitylate BACH1 based on the conformation as well as modification status of its BTB domain (Figure 7i). When BACH1 is released from DNA upon heme-binding, it is recognized by SCF^FBXO22^ through a native BTB dimer. However, a subset of S-nitrosylated or potentially otherwise modified BACH1 may evade recognition by SCF^FBXO22^ and, instead, gets recognized by SCF^FBXL17^ as a monomer^13, 14^. Notably, the ring-by-ring (RBR) E3 ligase HOIL-1 has also been implicated in BACH1 degradation under oxidative stress^11^. Unlike FBXO22 and FBXL17, HOIL-1 binds to the BACH1 disordered region in a heme-dependent manner. However, its exact function for BACH1 regulation remains to be determined.

In conclusion, our structural studies, together with detailed biochemical examinations of ubiquitylation reveal how the dual regulation by two complementary SCF-type E3 ligases post-transcriptionally controls BACH1 levels. These findings expand our understanding of the regulatory circuits underlying the oxidative stress response. Moreover, they may spur strategies to target BACH1 for degradation for the development of therapies for oxidative stress related disorders or cancer^24, 25, 34^.

## Acknowledgements

We thank Johann Wirsching for experimental support in the early stages of the project. We thank Anne Granger, William Forrester, Martin Schröder, and Jessica Desogus for feedback on the manuscript. We further thank the NBR Innovation Postdoctoral program for support of BG.

The results shown here are in part based upon data generated by the TCGA Research Network: https://www.cancer.gov/tcga.

## Author contributions

Conceptualization: BG

Data curation: BG, CF, PP, DA, MK, CF, CS

Investigation: BG, MK, PP, CS, DA, ND, JMR, SK, ZT, MM, DF, CM, FF, LT, BK, AH, SG, GR, ES, MI, CF

Visualization: BG

Writing – original draft: BG

Writing – original, review and editing: BG, CF, MI, MM, DF

## Materials and methods

### Antibodies

The following antibodies were used in this study.

| Antibody | Target | Source | Identifier | Dilution |
| --- | --- | --- | --- | --- |
| FBXO22 Polyclonal antibody | FBXO22 | Proteintech | #13606-1-AP | 1:1000 |
| BACH1 Polyclonal antibody | BACH1 | Proteintech | #14018-I-AP | 1:1000 |
| Vinculin (E1E9V) XP Rabbit mAb | Vinculin | Cell Signaling technologies | #13901 | 1:1000 |
| HA-Tag (C29F4) Rabbit mAb | HA tag | Cell Signaling technologies | #3724 | 1:1000 |
| Peroxidase AffiniPure Goat Anti-Rabbit IgG | Secondary antibody | Jackson ImmunoResearch | #111-035-144 | 1:2500 |
| Mouse anti His-Tag | His-tag | BioLegend | 652501 | 1:1000 |
| IRDye 680RD Streptavidin conjugate | biotin | LI-COR | 926-68079 | 1:1000 |
| IRDye 800CW Donkey anti-Mouse IgG | Secondary antibody | LI-COR | 925-32212 | 1:5000 |

### Plasmids & cloning

All plasmids used in this work were ordered from GeneArt.

SKP1-FBXO22 was cloned into a bicistronic flashbac vector for the simultaneous expression of SKP1 and FBXO22 using the *flash*BAC^TM^ system (Oxford Expression Technologies). While SKP1 was encoded in untagged form (full-length SKP1, residues 2-160), FBXO22 was encoded with an N-terminal truncation 11 (residues 12-403) as well as an N-terminal deca-histidine (His_10_) tag followed by a TEV-cleavage site. CUL1-RBX1 was also encoded a bicistronic flashback vector encoding full-length RBX1 (residues 1-108) and full-length CUL1 (residues 2-776) with an N-terminal TwinStrep-tag followed by a TEV cleavage site. In analogy, Avi-tagged SKP1-FBXO22 (^Avi^SKP1-FBXO22) and SKP1-FBXL17 (^Avi^SKP1-FBXL17) were cloned into a bicistronic flashbac vector with an N-terminally Avi-tagged SKP1 protein. FBXL17, comprising residues 310-701, was N-terminally tagged with a His_10_ tag followed by a maltose binding protein (MBP) tag and a TEV cleavage site.

Nedd8, Ubiquitin (Ub) and a lysine-deficient Ubiquitin variant (Ub^0^), in which all lysine residues were replaced by arginine, were cloned as fusion proteins with an N-terminal Hexa-histidine (His_6_)-tag followed by an HRV3C cleavage site and an artificial cysteine residue in a pJ201 vector modified with T7 promotor, T7 terminator, lac operon and lacI from pET41a(+) as regulatory elements. Nedd8 contained an additional ZZ-tag on the N-terminus.

Full-length BACH1 (BACH1^FL^, residues 2-736) was cloned into a flashback vector for expression using the *flash*BAC^TM^ system (Oxford Expression Technologies). The protein was fused to an N-terminal deca-histidine (His_10_) tag followed by an IgG binding domain of protein A (ZZ-tag)^38^ and an HRV3C-cleavage site. BACH1^BTB^ constructs (wildtype and mutants) and the extended BACH1 constructs BACH1^ΔC178^ (residues 7-177), BACH1^ΔC241^ (residues 2-240), BACH1^ΔC506^ (residues 2-505) were encoded as fusion proteins with an N-terminal His_10_-tag followed by a TEV cleavage site in a pJ201 vector modified with T7 promotor, T7 terminator, lac operon and lacI from pET41a(+) as regulatory elements.

The BACH2^BTB^ (residues 8-131), KEAP1^BTB^ (residues 48-180), and ZBTB16^BTB^ (1-130) domain constructs were designed in analogy to the BACH1^BTB^ construct.

A detailed overview of the protein sequences is available in Supplementary Table 1.

### Protein expression & purification

#### SKP1-FBXO22

The binary complex of wildtype SKP1-FBXO22 and the FBXO22 R376P, R376D, Q307E, Q307R, R367W and R367L mutants were expressed in *Spodoptera frugiperda* 9 I (*Sf*9) cells. Baculoviruses for expression of SKP1-FBXO22 constructs were generated in *Sf*9 cells using the *flash*BAC^TM^ system (Oxford Expression Technologies). Protein expression was performed by infecting *Sf*9 cultures grown to a cell density of 1.6×10^6^ cells/mL in Sf-900^TM^ III SFM medium (Gibco) with baculoviruses for 72 hours at 27 °C. Cells were harvested via centrifugation at 5000 x g for 15 min at 4 °C using a Sorvall BIOS 16 centrifuge (Thermo Scientific). Pellets were flash frozen in liquid nitrogen and stored at -80 °C.

Cells cultured with 1 L medium were lysed with a glass douncer using 50 mL of ice cold 25 mM HEPES pH 8.3, 300 mM NaCl, 20 mM imidazole, 1 mM TCEP supplemented with benzonase and a protease inhibitor cocktail tablet (Roche). The lysate was subsequently cleared by centrifugation at 87,207 x g for 1 hr at 4 °C using a Sorvall LYNX 6000 centrifuge and a A27-8 x 50 fixed angle rotor (both Thermo Scientific). All subsequent purification steps were carried out in a cold room at 4 °C. The clear lysate was subjected to an IMAC purification. For this, the supernatant was loaded onto TALON resin, washed with wash buffer (25 mM HEPES pH 8.3, 300 mM NaCl, 20 mM imidazole, 1 mM TCEP), and eluted using elution buffer (25 mM HEPES pH 8.3, 300 mM NaCl, 300 mM imidazole, 1 mM TCEP). The IMAC eluate was dialyzed overnight against dialysis buffer (25 mM HEPES pH 8.3, 300 mM NaCl, 20 mM imidazole, 1 mM TCEP) in the presence of 5 mol-% His_10_-taged TEV protease. The tag-free protein was afterwards separated from the protease and the cleaved tag using a reverse IMAC procedure. For this, the dialyzed sample was loaded onto TALON resin and washed with wash buffer. The flowthrough and was fractions were concentrated and further purified via size exclusion chromatography (SEC) using 25 mM HEPES pH 7.5, 150 mM NaCl, 1 mM TCEP and a HiLoad 16/60 Superdex 200 pg column (Cytiva). The pure protein fractions were pooled, flash frozen in liquid nitrogen, and stored at -80 °C.

#### ^Avi^SKP1-FBXL17

SKP1-FBXL17 was expressed and purified in analogy to SKP1-FBXO22.

#### Nedd8∼CUL1-RBX1

CUL1-RBX1 was expressed in *Sf*9 cells using the flashback system as described for SKP1-FBXO22. Cells cultured with 1 L medium were lysed with a glass douncer using 50 mL of ice cold 50 mM HEPES pH 7.0, 300 mM NaCl, 1 mM TCEP, 5 % (v/v) glycerol supplemented with benzonase and a protease inhibitor cocktail tablet (Roche). The lysate was subsequently cleared by centrifugation at 87,207 x g for 2 hrs at 4 °C using a Sorvall LYNX 6000 centrifuge and a A27-8 x 50 fixed angle rotor (both Thermo Scientific). The clear lysate was loaded onto a 5 mL StrepTrapHP column using an ÄKTA pure chromatography system (Cytiva), washed with wash buffer (50 mM HEPES pH 7.0, 300 mM NaCl, 1 mM TCEP, 5 % (v/v) glycerol), and eluted using elution buffer (50 mM HEPES pH 7.0, 300 mM NaCl, 1 mM TCEP, 5 % (v/v) glycerol, 2.5 mM Desthiobiotin). The eluate was dialyzed overnight against dialysis buffer (50 mM HEPES pH 70, 300 mM NaCl, 20 mM imidazole, 1 mM TCEP) in the presence of 5 mol-% His_10_-taged TEV protease. The sample was then concentrated and further purified via size exclusion chromatography (SEC) using 50 mM HEPES pH 7.0, 150 mM NaCl, 1 mM TCEP, 5 % (v/v) glycerol and a Superose6 increase 10/300 GL column (Cytiva). The pure protein fractions were pooled, flash frozen in liquid nitrogen, and stored at -80 °C.

Active Nedd8∼CUL1-RBX1 was obtained by incubating a mix of 20 µM CUL1-RBX1, 50 µM Nedd8, 1 µM NAE1/UBA3 and 1 µM UBE2M in 50 mM Tris pH 7.5, 100 mM NaCl, 2.5 mM MgCl2, 0.6 mM ATP, 0.5 mM TCEP, 5 % (v/v) glycerol for 1 hr at room temperature. The mixture was subsequently purified via size exclusion chromatography (SEC) using 50 mM HEPES pH 7.0, 150 mM NaCl, 1 mM TCEP, 5 % (v/v) glycerol and a Superose6 increase 10/300 GL column (Cytiva). The pure protein fractions were pooled, flash frozen in liquid nitrogen, and stored at -80 °C.

#### Ub and Ub^0^

The plasmids encoding Ub or Ub^0^ were transformed into *Escherichia coli* BL21 cells via the heat-shock method. cells were grown at 37 °C, 200 rpm in Luria Bertani (LB) medium supplemented with 30 ug/mL kanamycin to an OD_600_ of 0.8. Expression was induced with 1 mM IPTG for 4 hrs at 37 °C. Cells were harvested via centrifugation at 5000 x g for 15 min at 4 °C using a Sorvall BIOS 16 centrifuge (Thermo Scientific). Pellets were flash frozen in liquid nitrogen and stored at -80 °C.

All purification and refolding steps were carried out at 4 °C. Cell pellets were resuspended in 20 mM Tris pH 8.0, 100 mM NaCl supplemented with benzonase, homogenized using a POLYTRON PT1300 D handheld disperser (Kinematica), and subsequently lysed under cooling with ice using a M110P Microfluidizer (Microfluidics). The lysate was subsequently cleared by centrifugation at 40,000 x g for 30 min at 4 °C using a Sorvall LYNX 6000 centrifuge and a A27-8 x 50 fixed angle rotor (both Thermo Scientific). The pellet was resuspended in 20 mM Tris pH 8.0, 100 mM NaCl and centrifuged again at 40,000 x g for 20 min at 4 °C using a Sorvall LYNX 6000 centrifuge and a A27-8 x 50 fixed angle rotor (both Thermo Scientific). Inclusion bodies were then solubilized in 50 mM Tris pH 8.0, 7.5 M guanidine, 1 mM EDTA, 50 mM DTT for 30 min at room temperature and non-solubilized material was removed via centrifugation at 40,000 x g for 20 min at 4 °C using a Sorvall LYNX 6000 centrifuge and a A27-8 x 50 fixed angle rotor (both Thermo Scientific).

For refolding, 10 ml solubilized inclusion body solution was diluted with 40 mL 50 mM Tris pH 8.0, 7.5 M guanidine, 1 mM EDTA, the rapidly diluted in 5 L of precooled 0.5 M Tris pH 8.0 and incubated for 1 hour. The solution was filtered through 1.2/0.5 µM Optsiflow filters (Millipore) and concentrated in a Pellicon TFF system to 400 mL using a Biomax-5 membrane. The solution was dialyzed against 4 L of 100 mM NaCl, 1 mM DTT overnight. The sample was then concentrated and incubated with 2 mol-% of HRV3C protease overnight. The cleaved His_6_-tag was removed using a reverse Nickel-NTA IMAC purification. For this, the sample was loaded onto TALON resin, and washed with wash buffer (20 mM Tris pH 8.0, 300 mM NaCl, 1 mM TCEP). The tag-free protein was obtained in the flowthrough and wash fractions, concentrated, and subsequently purified via SEC in 20 mM NaH_2_PO_4_ pH 7.2, 100 mM NaCl, 1 mM TCEP using a HiLoad Superdex 75 26/600 pg column (Cytiva).

For Cy5-labeling, Ub or Ub^0^ were incubated at 4°C overnight with a 10-fold molar excess of Cy5-maleimide (Sigmal Aldrich). The labeled protein was afterwards separated from free dye by SEC in 50 mM HEPES pH 7.0, 150 mM NaCl, 1 mM TCEP using a Superdex 30 Increase 10/300 GL column. The labeling efficiency (85%) was determined photometrically using the Nanodrop (ThermoFisher).

#### Nedd8

Need8 was expressed in *E*. *coli* in analogy to Ub and Ub^0^. All purification and refolding steps were carried out at 4 °C. The cell pellet was resuspended in 50 mM Tris pH 8.0, 300 mM NaCl, 4 mM DTT supplemented with benzonase, homogenized using a POLYTRON PT1300 D handheld disperser (Kinematica), and subsequently lysed under cooling with ice using a M110P Microfluidizer (Microfluidics). The lysate was subsequently cleared by centrifugation at 40,000 x g for 60 min at 4 °C using a Sorvall LYNX 6000 centrifuge and a A27-8 x 50 fixed angle rotor (both Thermo Scientific). The cleared lysate was then loaded onto Ni-NTA (QIAGEN) equilibrated in 50 mM Tris pH 8.0, 300 mM NaCl, 20 mM imidazole, 4 mM DTT. After washing with 10 CV, the protein was eluted in 5 CV using 50 mM Tris pH 8.0, 300 mM NaCl, 500 mM imidazole, 4 mM DTT. The fusion was subsequently cleaved off by incubation in the presence of 1 mol-% HRV3C protease overnight at 4°C and tag-free Nedd8 was obtained via SEC in 50 mM HEPES pH 7.4, 200 mM NaCl, 0.5 mM TCEP, 10% (v/v) glycerol using a HiLoad Superdex 75 26/600 pg column (Cytiva).

#### BACH1

Full-length BACH1 (BACH1^FL^, residues 2-736) was expressed in *Trichoplusia ni* High Five (Hi5) cells using the *flash*BAC^TM^ system as described for SKP1-FBXO22 above. The lysis, IMAC purification, tag-cleavage followed the exact same protocol as described for SKP1-FBXO22 with the exception that His_10_-taged HRV3C protease was used for tag removal. The tag-free BACH1^FL^ protein was finally purified via size exclusion chromatography (SEC) using 25 mM HEPES pH 7.5, 150 mM NaCl, 1 mM TCEP and a Superose6 increase 10/300 GL column (Cytiva). The pure protein fractions were pooled, flash frozen in liquid nitrogen, and stored at -80°C.

BACH1^BTB^ constructs (wildtype and mutants) and the extended BACH1 constructs (BACH1^ΔC177^, BACH1^ΔC240^, BACH1^ΔC505^) were expressed in *E*. *coli* BL21 cells. The plasmids were transformed via the heat-shock method and cells were then grown in Terrific Broth medium supplemented with 50 ug/mL Kanamycin to an OD_600_ of 2.0, cooled to 18 °C, induced with 0.1 mM IPTG, and incubated overnight. Cells were harvested via centrifugation at 5000 x g for 15 min at 4 °C using a Sorvall BIOS 16 centrifuge (Thermo Scientific). Pellets were flash frozen in liquid nitrogen and stored at -80 °C.

For protein purification, the cell pellets were suspended in lysis buffer (25 mM HEPES pH 8.3, 300 mM NaCl, 20 mM imidazole, 1 mM TCEP), homogenized using a POLYTRON PT1300 D handheld disperser (Kinematica), and subsequently lysed under cooling with ice using a M110P Microfluidizer (Microfluidics). The lysate was subsequently cleared by centrifugation at 87,207 x g for 1 hr at 4 °C using a Sorvall LYNX 6000 centrifuge and a A27-8 x 50 fixed angle rotor (both Thermo Scientific). All subsequent purification steps were carried out in a cold room at 4 °C in analogy to SKP1-FBXO22.

#### Sample preparation for single particle cryo-EM analysis

The complex of SKP1-FBXO22 bound to BACH1^BTB^ for single particle cryo-EM analysis was generated by mixing SKP1-FBXO22 with a 1.5-fold molar excess of BACH1^BTB^ dimer and incubating the mixture at 4 °C for 1 hr. The SKP1-FBXO22/ BACH1^BTB^ complex was subsequently isolated via size exclusion chromatography (SEC) using 25 mM HEPES pH 7.5, 150 mM NaCl, 1 mM TCEP and a Superdex 200 increase 10/300 GL column (Cytiva). The sample was then concentrated to 16 mg/mL and immediately used for cryo-EM specimen preparation.

For cryo-EM analysis of SKP1-FBXO22 alone, the protein was thawed and purified via size exclusion chromatography (SEC) using 25 mM HEPES pH 7.5, 150 mM NaCl, 1 mM TCEP and a Superdex 200 increase 10/300 GL column (Cytiva) to remove potential aggregates. The sample was then concentrated to 17 mg/mL and used for cryo-EM specimen preparation.

#### Isotope labeling for NMR spectroscopy

^15^N and ^13^C labeled BACH1 constructs were obtained by overexpression in M9 minimal medium supplemented with 1 g/L ^15^NHCl as the sole nitrogen source and ^13^C-glucose as the sole carbon source. Deuteration was achieved using D_2_O based M9 minimal medium. For ^13^C-labeling of the Ile-δ_1_, Met-ε, Val-γ_1_/γ_2_, and Leu-δ_1_/δ_2_ and Ala-β methyl groups, the D_2_O based M9 medium was supplemented with with α-ketobutyrate (3,3-^2^H-4-^13^CH_3_; 50 mg/L), methionine (methyl-^13^CH_3_; 100 mg/L) and α-ketoisovalerate (3-^2^H- 3-(methyl-^13^CH_3_)-4-^13^CH_3_; 100 mg/L) 1 h prior to induction^39^.

A list of all used protein constructs can be found in Supplementary table 1.

### Protein biotinylation

N-terminally Avi-tagged SKP1 was coexpressed with His_10_-TEV-FBXO22(12-403) or His_10_-TEV-MBP-FBXL17(310-701) and purified as described for untagged variants to yield ^Avi^SKP1-FBXO22 or ^Avi^SKP1-FBXL17, respectively. Biotinylation reactions were carried out *in vitro* by incubating 50 µM ^Avi^SKP1-FBXO22/FBXL17 with 5 μM BirA enzyme and 200 µM D-Biotin in 50 mM HEPES pH 7.5, 300 mM NaCl, 5 mM MgCl2, 1 mM TCEP and 20 mM ATP overnight at 4°C. Complete biotinylation was confirmed via LC-MS analysis and biotinylated ^Avi^SKP1-FBXO22 or ^Avi^SKP1-FBXL17 was purified by SEC using 25 mM HEPES pH 7.5, 150 mM NaCl, 1 mM TCEP and a Superdex 200 increase 10/300 GL column (Cytiva). Protein containing fractions were concentrated to 1.5 mg/mL, aliquoted, flash frozen in liquid nitrogen and stored at -80°C.

### Cysteine modification of BACH1 constructs

BACH1^BTB^ or BACH1^ΔC178^ were transferred into reducing agent-free buffer via SEC using a Superdex 75 increase 10/300 GL column (Cytiva) and 25 mM HEPES pH 7.5, 150 mM NaCl. Proteins were incubated at 50 µM in the presence of 10x excess of Bromo-1,1,1-trifluoroacetate (BTFA, Sigma Aldrich) overnight at 4°C. Proteins were then separated from free BTFA via SEC into 25 mM HEPES pH 7.5, 150 mM NaCl, 1 mM TCEP. Labeling was validated via LC-MS.

### Mass photometry

All mass photometry data were collected on a Refeyn OneMP mass photometer with a 10.8 × 2.9 μm (128 × 35 pixels) field of view^40^. Landing assay measurements were carried out in silicone gaskets (3 mm × 1 mm, GBL103250, Grace Bio-Labs) on microscopy slides. Protein solutions (3 μl) were added to the gaskets containing 17 μl buffer and images were acquired for 60 s at 100 Hz. Landing assays were analyzed using DiscoverMP (Refeyn Ltd) to extract particle contrasts.

Calibration curves for mass determination were generated using stock solutions of 200 nM bovine serum albumin (66 kDa monomer, 132 kDa dimer) and 200 nM thyroglobulin (660 kDa) in phosphate buffered saline (PBS). For sample measurements, protein stock solutions of SKP1-FBXO22 and BACH1^FL^ were prepared at a concentration of 200 nM in phosphate buffered saline (PBS). For the SKP1-FBXO22/BACH1^FL^ sample, a stock solution of 200 nM SKP1-FBXO22 was mixed with 200 nM BACH1^FL^ to yield a 1:1:2 stoichiometry of SKP1, FBXO22 and BACH1^FL^. After addition of 3 µL of the sample stocks to the 17 µL PBS drop on the gasket, the protein concentrations were 30 nM in PBS. All samples were measured at room temperature.

### Surface plasmon resonance

SPR experiments were performed using a Biacore T-200 instrument (GE Healthcare) in 50 mM HEPES pH 7.5, 150 mM NaCl, 1 mM TCEP, 0.01% Tween20 buffer at 15 °C. Biotinylated ^Avi^SKP1-FBXO22(12-403) was immobilized on a high-affinity streptavidin (SA) sensor chip (Cytiva) surface at a density of 200 RU. BACH1^BTB^ was serially diluted in the running buffer and injected individually. Cycles were run with at a a flow rate of 30 μL/min with 60 s contact time and 420 s dissociation using a stabilization period of 30 s and syringe buffer wash steps between injections. Data analysis was carried out using the Biacore Evaluation Software (GE Healthcare). All data processed by double referencing to the reference channel and blank injection and the resulting sensorgrams were fitted with a kinetic model assuming a 1:1 binding mode for *K*_D_ estimation.

Kinetic data obtained by SPR are summarized in Supplementary table 2.

### Biolayer interferometry

BLI experiments were performed at room temperature on a BLItz instrument (ForteBio) using Octet® Streptavidin Biosensor tips (Sartorius) and 4 µL sample drop holder. Measurements of the SKP1-FBXO22 interaction with BACH1^BTB^ were carried out in 25 mM HEPES pH 7.5, 150 mM NaCl, 0.01% (v/v) Tween20 and measurements of the SKP1-FBXO22 interaction with BACH1^FL^ were carried out in 25 mM HEPES pH 7.5, 300 mM NaCl, 0.01% (v/v) Tween20. Heme-dependent binding experiments were carried out in buffer containing 5 µM hemin. Prior to the measurements, the analyte, BACH1^BTB^ and BACH1^FL^, was incubated for 30 min in the hemin supplemented buffer.

In each cycle, the tip was equilibrated in buffer for 30 sec and biotinylated ^Avi^SKP1-FBXO22 was immobilized at a concentration of 25 µg/mL for 120 secs followed by a 30 sec equilibration in buffer. The tip was then incubated in analyte solution for a 120 sec associate phase and the cycle was closed by a 120 sec dissociation phase in buffer. The data were analyzed using the BLItz Pro software (ForteBio).

Kinetic data obtained by BLI are summarized in Supplementary table 2.

### Time-resolved fluorescence resonance energy transfer assay

TR-FRET competition assays were measured in white 384-well plates with a total assay volume of 10 µL per well and 25 mM HEPES pH 7.5, 150 mM NaCl, 1 mM TCEP, 0.01% Tween20, 0.1 mg/mL BSA as assay buffer. The TR-FRET mixtures contained 10 µM His_10_-TEV-BACH1^BTB^, 100 nM biotinylated ^Avi^SKP1-FBXO22, 2 nM MAb Anti 6HIS-Tb cryptate HTRF reagent (Cisbio), 25 nM Streptavidin Alexa Fluor™ 488 conjugate (Invitrogen) and increasing concentrations of tag-free BACH1 or SKP1-FBXO22 constructs, respectively. The mixtures were allowed to equilibrate by incubation at RT for 30 min. The fluorescence at

520 nm and 490 nm was detected using PheraStar fluorescence reader (BMG Labtech) with the LanthaScreen Optic module, Flash lamp Excitation source, and 30 flashes per well. The TR-FRET ratio (520 nm / 490 nm) was normalized to a negative control (no competitor) as 1 and a positive control (no His_10_-TEV-BACH1^BTB^ and biotinylated AviSKP1-FBXO22) as 0. IC_50_ values were obtained in GraphPad by fitting the data with a symmetrical sigmoidal curve using a fixed upper limit of 1, a fixed lower limit of 0 as the only constraints.

Fit parameters obtained by TR-FRET are summarized in Supplementary tables 3-5.

### Differential scanning fluorimetry

Differential scanning fluorimetry was carried out using a Prometheus Panta system (Nanotemper). For melting temperature analysis of SKP1-FBXO22 constructs, the protein was used at a concentration of 5 µM in 25 mM HEPES pH 7.5, 150 mM NaCl, 1 mM TCEP in Prometheus NT.48 Series nanoDSF Grade Standard Capillaries. Samples were heated from 25 to 90 °C at a rate of 1 °C/min while monitoring the fluorescence at 330 and 350 nm. The fluorescence ratio between 350 and 330 nm (F350/F330) was used for melting temperature determination. For aggregation analysis of BACH1^BTB^ constructs, samples were used at 10 µM concentrations using Prometheus NT.48 Series nanoDSF Grade High Sensitivity Capillaries. Samples were heated from 25 to 85 °C at a rate of 1 °C/min. Data were analyzed using the Prometheus software. Melting and aggregation points were determined using the Boltzmann and the derivative method.

Melting temperatures obtained by nano-DSF are summarized in Supplementary table 6.

### Analytical and fluorescence-detected size exclusion chromatography

Analytical (aSEC) and fluorescence-detected size exclusion chromatography (fSEC) experiments were carried out on an Agilent 1200 series Gradient HPLC system equipped with an Agilent 1260 Infinity Fluorescence Detector (G1321B) and an Agilent 1260 Infinity Diode Array Detectors DAD G1315C. All fSEC experiments were carried out at 25 °C.

For binding studies, SKP1-FBXO22 constructs were used at 5 µM and BACH1 constructs at 10 µM final concentration in 25 mM HEPES pH 7.5, 150 mM NaCl, 1 mM TCEP. Samples were injected onto a Superdex 200 increase 5/150 GL column (Cytiva) using an Agilent 1260 Infinity Standard Autosampler (G1329B) with injection volumes set to 10 µL. Tryptophan fluorescence was monitored using an excitation wavelength of 295 nm and an emission wavelength of 330 nm. Protein absorption was detected at 280 nm and 260 nm.

Hemin-BACH1 binding studies were performed at 1 µM BACH1^FL^ or BACH1^CP0^ in the absence or presence of 5 µM hemin using a Superose 6 increase 5/150 GL column (Cytiva). Protein was detected at an absorption wavelength of 280 nm while hemin was detected at 390 and 412 nm.

### Multi-angle light scattering

Size exclusion chromatography coupled to multi-angle light scattering (SEC-MALS) was carried out on an Agilent 1260 Infinity II Multidetector system equipped with an Agilent 1260 Infinity II vialsampler (G7129A) connected to an Eclipse and miniDAWN system (Wyatt Technology).

Samples were used at concentrations between 1.5-3.0 mg/mL and a total of 30 µg were injected onto Superdex 200 increase 5/150 GL or Superose 6 increase 5/150 GL columns equilibrated in 25 mM HEPES pH 7.5, 150 mM NaCl, 1 mM TCEP, 0.01% NaN_3_. All samples were centrifuged at 16,000 x g, 4 °C for at least 30 min using an Eppendorf 5415R centrifuge prior to injection.

Data were processed using ASTRA (Wyatt Technology) assuming a Zimm model^41^. The molecular weight, *M_W_*, was obtained from the reduced Rayleigh ratio extrapolated to zero, *R*(0) according to equations **1**.

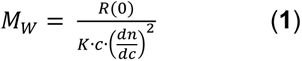

Here, *c* is the concentration of the analyte and (*dn*/*dc*) is the refractive index increment, for which a standard value of 0.185 mL/g was chosen^42^. *K* is a wavelength and solvent refractive index dependent optical constant. The protein extinction coefficients 280 nm were calculated from the respective amino acid sequences using the ProtParam tool^43^.

All molecular weights determined by SEC-MALS are summarized in Supplementary table 6.

### Circular dichroism spectroscopy

Far UV CD spectra were recorded at 25°C on a JASCO J-815 spectropolarimeter using MACRO CELL QS quartz glass High Performance cuvettes with 1 mm pathlength. Proteins were used at a concentration between 0.1-0.2 mg/mL in 1.25 mM HEPES pH 7.5, 7.5 mM NaCl. Spectra were recorded between 190-250 nm in 1 nm intervals. Spectra were obtained from the averaging of three consecutive accumulations per measurement. The measured ellipticity *θ_λ_* was converted to the mean residue ellipticity *θ*_mrw,_*_λ_* using equation **2**:

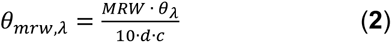

Here, MRW is the mean residue weight, *d* is the pathlength (in cm), and *c* is the protein concentration (in g/mL). The MRW is calculated from the molecular weight *M_W_* of the protein (in Da) and the number of amino acids *N* in the protein as MRW=*M_W_*/(*N*-1)^44^.

### In vitro ubiquitylation assay

In vitro ubiquitylation of BACH1 was performed by mixing 100 nM Nedd8∼CUL1-RBX1 with 100 nM SKP1-FBXO22 or SKP1-FBXL17, 1 µM BACH1 (or deletion constructs), 500 nM E2 (UbcH5c, BostonBiochem), 50 µM E1 (UBA1, BostonBiochem), and 25 µM Cy5-labeled Ub or Ub^0^, respectively, in 25 mM Tris pH 7.5, 250 mM NaCl, 5 mM MgCl_2_, 0.2 mM CaCl_2_, 0.5 mM TCEP, 12.5% glycerol, 0.1% Triton X-100, 0.1 mg/mL BSA. The mixture was allowed to equilibrate for 30 min on ice before adding 1 mM ATP to initiate the ubiquitylation reaction. Reactions were quenched at indicated time points by adding 4x Lämmli buffer with 50 mM DTT. Ubiquitylation was assessed by 4-20% SDS-PAGE followed by Cy5 fluorescence detection using an Amersham ImageQuant 800 (Cytiva) imager. Gels were subsequently stained with InstantBlue Coomassie Stain (Abcam) and scanned with the Gel Doc EZ system (BioRad).

Heme-dependent ubiquitylation assays were carried out with 0.5 µM His-tagged BACH1^FL^ or BACH1^CP0^ using 20 µM ubiquitin (Ub^0^ doped with 10% biotinylated Ub). BACH1 ubiquitylation was detected by WB. Biotinylated Ub was detected using a IRDye 680RD Streptavidin conjugate (LI-COR). His-tagged BACH1^FL^ or BACH1^CP0^ were detected using a mouse anti His-Tag primary antibody (BioLegend) combined with a IRDye 800CW Donkey anti-Mouse IgG (LI-COR) as secondary antibody. IR fluorescence was detected using an Odyssey DLx Imaging system (BioRad).

### Liquid chromatography coupled to mass spectrometry

Liquid chromatography coupled to mass spectrometry (LC-MS) experiments were carried out on an ACQUITY PREMIER HPLC system equipped with a ACQUITY UPLC® Protein BEH C4 Column (2.1x100 mm) coupled to a Xevo G3-XS QTof MS(ESI) mass spectrometer (Waters Corporation). Eluent A contained 0.05% TFA in H_2_O while eluent B contained 0.05% TFA in isopropanol. Samples were injected at 3 µL volumes between 0.5-1.0 mg/mL concentration and separated at a flow rate of 0.5 mL/min using a gradient program starting with5% eluent B for, followed by a gradient from 5-60% B for 8.0 min, and 60-98% B for

8.0 min. MS data were deconvoluted using the maximum entropy (MaxEnt) method and analyzed using the MassLynx software (Waters Corporation).

### Generation of U2OS FBXO22 knockout cells

#### Cell culture

U2OS cells were cultured in DMEM, high glucose, GlutaMAX^TM^ Supplement (Thermo Fisher Scientific #31966021) supplemented with 10% FBS (GE Healthcare) and 100 U/ml penicillin-streptomycin (100x stock, Thermo Fisher Scientific #15140122). Cells were passaged every 3-5 days by dilution of 1:3 to 1:6, after releasing the attached cells using TrypLE (Thermo #12605036).

HEK293T cells were cultured with DMEM, high glucose, GlutaMAX™ Supplement, pyruvate (Thermo Fisher Scientific #31966021) supplemented with 10% heat inactivated-FCS (Bioconcept, Cat#2-01F36-I), 10 mM Hepes (Thermo Fisher Scientific #15630080). Cells were passaged every 2-3 days, by diluting the cells 1:10 to 1:15, after releasing the attached cells using TrypLE (Thermo Fisher Scientific #12605036).

For lentiviral particle production, 200,000 HEK293T cells/well were seeded in Poly-D-Lysin (Sigma #P6407) coated 24-well plates one day prior to transfection. The cells were transfected with 0.05 µg of DNA of interest with 0.2 µg of ready-to-use lentiviral packaging plasmid mix (Cellecta #CPC-K2A) using TransIT®-293 Transfection Reagent (Mirus Bio LLC #MIR2700) according to the manufacturer’s protocol. Subsequently, the media was changed 24 hours after transfection and lentivirus-containing supernatants were collected after 72 hours and filtered using a cellulose acetate membrane, pore size 0.45 μm, diam. 30 mm (Sigma #Z741871).

#### Knockout cell line generation

Constitutive Cas9 expressing U2OS cells were generated by lentiviral delivery of the Cas9 protein gene in pNGx-LV-c004 and selected with 10 µg/ml blasticidin S HCl (Thermo Fisher Scientific #A1113902) as previously described^45^. Cas9 expression in U2OS clones were verified with TaqMan Assay (Forward primer: 5’- AGAGCTGCACGCAATCCT-3’, Reverse primer: 5’- CGCGGTTATCCTTCAGAAA-3’, Probe: 5’- Fam-CGACGACAGGAAGACTTCTACCCA- BHQ-1 -3’). A single U2OS Cas9 cell clone was then transduced with FBXO22 targeting sgRNA using pNGx-LV-g003 backbone (sgRNA sequence: 5’- TGGTGAGGAATGGAGCCGGT-3’)^45^. Lentiviral particles were generated as previously described. 90,000 U2OS cells/well were seeded in 6-well with 8 ug/ml polybrene (Millipore #TR-1003-G) addition to the growth medium and infected with 4ul of lentiviral particle containing supernatant. 24-hours after infections, cells were selected with 1.5ug/ml puromycin (Thermo Fisher Scientific #A11138-03) for three days. 90% of infection rate detected by RFP content measurement by FACS analysis on BD Fortessa.

Knockout efficiency was confirmed by protein detection by western blotting.

### Co-immunoprecipitation assays

#### Cell culture and transfection conditions

U2OS parental and U2OS FBXO22 KO cells were maintained in McCoy’s 5A medium (Bioconcept 1-18F01-I) supplemented with 10% fetal bovine serum (SIGMA F7524), 2 mM GlutaMAX Supplement (Thermo Fisher Scientific 35050-038).

For transient expression of FBXO22 transgenes, gene fragments coding for wild-type and mutant HA-tagged FBXO22 proteins were synthesized and cloned into the pXP1510 plasmid (Twist Bioscience). U2OS FBXO22 KO cells were seeded to 60% confluency on plates coated with 0.1% gelatin (Sigma G13393) and the plasmids were transfected using X-tremeGENE™ 9 DNA Transfection Reagent (Roche XTG9-RO) according to the manufacturer’s protocol.

Cells were collected 48 hrs after transfection and lysed in NP-40 extraction buffer (50 mM Tris-HCl pH 7.5, 1% NP-40, 120 mM NaCl, 20 mM NaF, 1 mM EDTA, 6 mM EGTA, 15 mM sodium pyrophosphate, 1 mM benzamidine, 0.5 mM PMSF, 1mM sodium orthovanadate) including cOmplete™, Mini Protease Inhibitor Cocktail (Roche 11836153001).

#### Co-Immunoprecipitation

Pierce™ Anti-HA magnetic beads (Thermo Fisher Scientific 88836) were used for immunoprecipitation of recombinant HA-tagged proteins and co-immunoprecipitation of their interacting proteins. Briefly, 40 µL pre-washed magnetic beads were incubated with 100 µg cell lysates for 2 hrs at 4°C with rotation. Beads were subsequently washed four times with NP-40 extraction buffer using a magnetic separation rack and proteins were eluted in SDS-PAGE sample buffer by heating the sample at 95 °C for 5 min.

#### Western blots

Samples were analyzed by Western blotting using specific primary antibodies indicated above. After blocking in 1x PBS, 0.1% Tween20, 5% milk buffer for 30 min, PVDF membranes were incubated with primary antibodies O/N at 4°C followed by the secondary antibody for 2 hrs at RT.

### Hydrogen-deuterium exchange mass spectrometry

SKP1-FBXO22 wt and R376P mutant stock solutions were prepared in 25 mM Tris, 150 mM NaCl at pH 7.5, BACH1 stock solution was prepared in the same buffer as SKP1-FBXO22. HDX-MS sample solutions were prepared by mixing 50 µL of 44 µM SKP1-FBXO22 plus 50 µL sample buffer (±118 µM BACH1). The solution was incubated for 30 min prior to initiation of HDX experiments. HDX labeling was performed in the stock buffer prepared with D_2_O pD 7.5. An exchange reaction was initiated by diluting 2 µL of SKP1-FBXO22 sample solution (±118 µM BACH1) with 27 µL deuterated buffer. The labeling reaction mixture was incubated at 18 °C for varying times (30 to 10,000 sec). These partially deuterated samples were then subjected to HDX-MS analysis as follows. The labeling reaction was quenched at a 1:1 ratio with a solution of 3.2 M GuHCl, 0.8% Formic Acid pH 2.1 at 2 °C. The quenched solution was then injected onto an integrated fluidics system containing an HDx-3 PAL liquid handling robot and a temperature-controlled chromatography chamber set at 1.5 °C (Trajan Scientific), a Dionex Ultimate 3000 UHPLC system, and a Q-Exactive HFX hybrid quadrupole Orbitrap mass spectrometer (Thermo Fisher Scientific). Mobile phase A: 0.1% Formic Acid in Water and mobile phase B: 0.1% formic acid in 95% Acetonitrile in Water. The protein is first loaded onto a mixed protease column (NovaBioAssays, FPXII:Pepsin (w/w 1:1) 2.1x30 mm, NBA2014002) at 100 µL/min for 3 minutes at 5 °C. The resulting peptides are trapped and desalted on a C18 trapping column (Acclaim 5 µm, PepMap 300, 1 mm x 15 mm, Fisher Scientific). Once desalting is complete the trap column is placed in line with a separation column (Hypersil GOLD 1.9 µm particle size, 50 x 1 mm, Fisher Scientific) and eluted with using a gradient of 10-35 %B over 7 min. MS experiments were acquired over a scan range of 350 to 1500 m/z on a Thermo Q-Exactive HFX.

### NMR spectroscopy

#### NMR assignments

For backbone resonance assignments, dimeric, uniformly ^2^H,^13^C,^15^N-labeled BACH1^BTB^ was prepared in 20 mM d_18_-HEPES, 150 mM NaCl, 1 mM d_16_-TCEP, 5% D_2_O, pH 6.5 and concentrated to 500 µM in the presence of 5% d_8_-glycerol. All NMR spectra were measured in 3 mm NMR tubes with sample volumes of 170 µl at an initial protein concentration of 500 µM. All experiments were collected at 23°C on a Bruker Avance III HD 800 MHz equipped with four radio-frequency channels for generating the ^1^H, ^2^H, ^13^C and ^15^N pulses and a 5 mm ^1^H{^13^C,^15^N}-triple resonance cryogenic probe with shielded xyz-gradient coils. Backbone assignments were obtained using 2D (^1^H,^15^N)-HSQC^46^/(^1^H,^15^N)-TROSY^47^ and 3D ^2^H-decoupled HNCA, HNCACB and HN(CO)CACB experiments^48^, complemented with a 3D ^15^N-edited NOESY (120 ms mixing time)^49^. 2D (^1^H,^15^N)-TROSY spectra of four single residue mutants of BACH1^BTB^, namely F9A, E12R, A53V and F125A, where used to support backbone resonance assignments. The NMR data were processed with the software Topspin 3.6 (Bruker, Switzerland) and analyzed with CCPNMR^50, 51^.

Overall, we achieved backbone N+H assignments for 111 of 120 expected residues (92.5%). Moreover, we obtained 91.9% of the expected Cα assignments and 70.8% Cβ assignments. Unfavorable relaxation and peak overlap prevented us to assign residues S14, V36, I38, F39, V40, R49, T93, A94, K95. All assignment data have been deposited in the BMRB under the ID 52341.

#### Binding experiments

All experiments were carried out at 296 K. For NMR experiments with ^15^N-labeled protein, (^1^H,^15^N)-TROSY NMR spectra of 50 µM U-^15^N-BACH1^BTB^ were measured on a Bruker Avance III HD 800 MHz spectrometer in the absence and presence of 100 µM SKP1-FBXO22 or SKP1-FBXL17, respectively. For NMR experiments with ^13^C-methyl labeled protein, (^1^H,^13^C)-HMQC NMR spectra of 50 µM U-(^2^H,^15^N), (ILVMA)- ^13^CH_3_ labeled BACH1^BTB^ were measured in the absence and presence of 100 µM SKP1-FBXO22.

Titration experiments were conducted by measuring (^1^H,^13^C)-HMQC NMR spectra 20 µM µM (U-^2^H,^15^N), (ILVMA)-^13^CH_3_ labeled BACH1^BTB^ on a Bruker Avance III HD 800 MHz spectrometer at increasing SKP1-FBXO22 concentrations. Samples for each concentration step were set up in separate tubes to avoid dilution effects upon adding ligands to the apo protein.

Hemin binding to the BACH1 BTB domain was tested by measuring (^1^H,^15^N)-TROSY NMR spectra of 75 µM (U-^15^N)-BACH1^BTB^ in 25 mM HEPES pH 7.5, 150 mM NaCl, 10% D2O, 30 µM DSS in the absence or presence of 100 µM hemin on a Bruker Avance III HD 800 MHz spectrometer. Hemin was added from a 3.75 mM d^6^-DMSO stock.

#### Cysteine modification in BACH1^BTB^

All experiments were carried out at 296 K. (^1^H,^15^N)-SOFAST-HSQC NMR spectra of 100 µM U-^15^N-BACH1^BTB^ in 25 mM HEPES pH 7.5, 150 mM NaCl, 10% D_2_O were measured on a Bruker Avance III HD 800 MHz spectrometer in the absence and presence of 1 mM BTFA, 1 mM iodoacetamide or 1 mM S-nitrosylgluathione (both 100 mM DMSO stocks). Proteins were incubated for at least 5 hours with the modifying agents at 4°C in the dark before measurements.

#### {^1^H},^15^N-het*NOE* experiments

{^1^H},^15^N steady-state nuclear Overhauser effect measurements ({^1^H},^15^N-*NOE*) were performed at 296 K on a Bruker Avance III HD 800 MHz spectrometer using a standard Bruker pulse sequence. Separate 2D ^1^H,^15^N spectra of 150 µM U-^15^N-BACH1^BTB^ were acquired with and without continuous ^1^H saturation, respectively. The het*NOE* values were determined by taking the ratio of peak volumes from the two spectra, het*NOE*=*I*_sat_*/I*_0_, where *I*_sat_ and *I*_0_ are the peak heights with and without ^1^H saturation. Peak heights and errors were obtained using CCPNMR version 3^50, 51^. Errors were obtained from the peak height errors of the individual peaks by error propagation.

#### Hydrogen-deuterium exchange experiments

Hydrogen-deuterium exchange (HDX) experiments were conducted at 296 K on a Bruker Avance III HD 800 MHz spectrometer using 150 µM U-^15^N-BACH1^BTB^. Reference spectra of U-^15^N-BACH1^BTB^ were measured in 20 mM HEPES pH 7.5, 150 mM NaCl, 1 mM TCEP, 10% D_2_O. For HDX measurements, the protein was transferred to 20 mM HEPES pH 7.5, 150 mM NaCl, 1 mM TCEP, 80% D_2_O and consecutive (^1^H,^15^N)-SOFAST-HSQC spectra were measured for 60 hrs. The peak heights at a given time point were determined using CCPNMR version 3^50, 51^ and the HDX (in %) was calculated by dividing through the peak heights in the reference spectrum, i.e. at 10% D_2_O, thus yielding HDX values between 20% (maximum HDX) and 90% (minimum HDX).

### Cryo-electron microscopy

#### Cryo-EM specimen preparation and data collection

Concentrated SKP1/FBXO22 sample to 16 mg/ml and SKP1-FBXO22/BTB dimer to 20 mg/ml were used to prepare cryo-EM grids. Vitrified specimens were prepared by applying 5 µL of protein complexes onto glow discharged µLtrafoil R1.2/1.3 Au 200 mesh. Cryo-EM grids were flash frozen in liquid ethane using a Vitrobot Mark IV (Thermo Fisher Scientific, Waltham, MA, USA). Data were collected on a Titan Krios microscope (Thermo Fisher Scientific, Waltham, MA, USA) equipped with a Cs-corrector hardware operated at an accelerating voltage of 300 kV with a 50 μm C2 aperture at an indicated magnification of 96 000× in nanoprobe mode and a spot size of 5 for SKP1-FBXO22 dataset and indicated magnification of 75000x in nanoprobe mode for SKP1-FBXO22/BACH1^BTB^ complex sample. A Falcon4i Direct Electron Detector camera operated in Electron-Event representation (EER) mode electron was used to acquire dose-fractionated images of the SKP1-FBXO22 using 0.656 Å/pixel and 0.84 Å/pixel for SKP1-FBXO22/BACH1^BTB^ complex samples. All Movies were recorded with an exposure time of 3.07 s for the SKP1-FBXO22 sample and 3.34 s for the SKP1-FBXO22/BACH1^BTB^ complex sample. A total exposure of

50 e^−^/Å^2^ with an exposure rate of 10.13-11.52 e^−^/pixel/second was fractionated into 50 subframes. Defocus was varied in the range between −0.6 to −1.6 μm. Beam-image shift was used to acquire data from 9 surrounding holes and 4 acquisitions per hole after which the stage was moved to the next collection area using Thermo Fisher Scientific EPU software. This allowed for a higher throughput data collection, corresponding to an acquisition rate of more than 500 micrographs/hour.

### Cryo-EM data processing and 3D reconstruction

#### Data processing of SKP1-FBXO22

Image processing was done using Relion 4^52^ and CryoSPARC™^53^ image processing software. After motion-correction (using Relion’s implementation of MOTIONCOR^54^, CTF estimation was done using patch CTF estimation in CryoSPARC. particles were picked using a blob picker algorithm implemented in CryoSPARC. After picking particles and performing few rounds of 2D classification, selected 2D classes were used to repick particles using template picker algorithm implemented in CryoSPARC. A total of 6062563 particles were picked from 8720 micrographs of SKP1-FBXO22 dataset and extracted using 240 px box. Following two rounds of 2D classification and one round of *ab-initio* in CryoSPARC, the best class was chosen as a reference for further refinement. At the end non-uniform refinement was carried out using 717992 particles yielding a reconstruction with a global resolution of 3.4 Å. The map resolution was determined based on the gold-standard 0.143 criterion.

#### Data processing of the SKP1-FBXO22/BACH1^BTB^ complex

18037 micrographs were used to pick particles in CryoSPARC using blob picking algorithm. Particles were extracted using 200 px box. Representative 2D classes of the sample were used for template picking in CryoSPARC. After performing few rounds of 2D classification, ab initio reconstruction and hetero refinement, the good class from hetero refinement in CryoSPARC was used to perform non-uniform refinement job. At the end it yielded a final map with a global resolution of 3.2 Å based on the gold-standard 0.143 criterion

### Cryo-EM map interpretation

Atomic models of the SKP1-FBXO22 heterodimer and the SKP1-FBXO22/BACH1^BTB^ complex were generated with AlphaFold v2.0 multimer^55, 56^ and fitted as rigid bodies into the respective cryo-EM density maps in coot^57^. Unresolved regions were manually deleted in the models in coot. The models were real space refined in PHENIX^58^ and validated using MolProbity^59^.

All cryo-EM data are summarized in Supplementary table 8.

### Molecular dynamics simulations

The SKP1-FBXO22 complex was prepared for simulation with and without BACH1. The cryo-EM structure of the full SKP1-FBXO22/BACH1^BTB^ complex was used as a starting point. The conformations of some loops that were not resolved in that structure but could be resolved in the SKP1-FBXO22 cryo-EM structure were combined with the full SKP1-FBXO22-BACH1 structure using Schrodinger Maestro’s homology modelling tools to make a chimeric homology model^60^. To create the model without BACH1, BACH1 was simply removed, and the resulting model saved separately. Both models were then prepared using MOE protein preparation tools^61^. MD systems for both models were set up using Amber Tools 22^62, 63^, the Amber ff19SB forcefield^64^, and the TIP3P solvent model ^65^. Buried water positions were calculated using 3D-RISM^66^ with the SANDER interface. These were used in an orthorhombic water box that also contained a

### 0.15 M concentration of NaCl salt

Both systems were then minimized with progressively weaker restraints. This began with 400 steps of minimization, applying a harmonic force restraint of 1.0 kcal·mol^-1^·Å^-2^, of which the first 50 iterations used the steepest descent method and the rest conjugate gradients. A second minimization of 10 000 steps was performed with a harmonic force restraint of 0.1 kcal·mol^-1^·Å^-2^, of which the first 5000 steps used the steepest descent method and the rest conjugate gradients. A final minimization of 10 000 steps was performed without restraints, of which the first 5000 steps once again used the steepest descent method and the rest conjugate gradients. Equilibration was performed by first heating up the system to 100 K with Langevin dynamics^67^ (collision frequency of 1 ps^-1^) in a constant volume 5 ps simulation and applying a 5 kcal·mol^-1^·Å^-2^ harmonic force restraint. Bonds involving hydrogen were constrained using the SHAKE algorithm^68^, which allowed for 2 fs timesteps. The system was then heated from 100 K to 310 K at constant pressure in 500 ps also with a 5 kcal·mol^-1^·Å^-2^ harmonic force restraint. Production runs were performed without positional restraints and using hydrogen mass repartitioning^69^ to facilitate 4 fs timesteps. Both systems were simulated in triplicates for 1 μs using PMEMD.CUDA on graphics processing units (GPUs)^70–72^.

The simulations were analyzed in three ways. First, we analyzed the root-mean-square-deviation (RMSD) of the backbone atom positions of the system with respect to their average positions in the simulation. Next, we determined the root-mean-square-fluctuation (RMSF) of backbone atom positions of residues also with respect to their average positions. Finally, we analyzed the time-resolved presence of secondary structures as defined by the DSSP algorithm^73^. All analyses were run with CPPTRAJ^74^.

### Retrieval of data from The Cancer Genome Atlas

Cancer-associated mutations in BACH1 and FBXO22 were retrieved from The Cancer Genome Atlas (TCGA) database (https://www.cancer.gov/tcga). For BACH1, we identified the mutations Y11H (UUID: 2c28b2e9-ac86-59f7-8533-dda6008e7f40) and A53V (UUID f42de62e-f16c-5607-88fb-03a2433e200e) as potentially relevant for the interaction with FBXO22 or FBXL17. For FBXO22, we identified the mutations Q307E (UUID: a0a1dbc9-4bf0-5ecf-837d-5b883e54581d), Q307R (UUID: 9ddb8d67-4896-5fcd-89a1-b70b8ae6c2f6), R367W (UUID: d9fc3434-35f8-5ad9-970a-b3facf7ec033), and R367L (UUID: 6e147f7b-277b-5804-80da-545592ff8252) as potentially modulating the interaction with BACH1.

## Data and code availability

Atomic coordinates have been deposited to the Protein Data Bank under accession codes 8S7D and 8S7E. Electron microscopy-derived density maps were deposited in the Electron Microscopy Data Bank under accession codes 19766 and 19768. Backbone NMR assignments have been deposited to the Biological Magnetic Resonance Bank under accession code 52341. Molecular dynamics simulation data were deposited in a Zenodo repository under the following link: https://zenodo.org/doi/10.5281/zenodo.10723643.

## Materials availability

This study did not generate new unique reagents.

