## Supplementary information for "Structural basis of dual BACH1 regulation by SCF^FBXO22^ and SCF^FBXL17^"

**Supplementary table 1: Recombinant protein constructs used in this study.**

| Name | UniProt ID / Fusion tag / amino acid sequence |
|---|---|
| **$^{Avi}$SKP1-FBXO22** | P63208/Q8NEZ5 — Avi-lin on SKP1, His$_{10}$-TEV-lin on FBXO22<br>Avi-lin-SKP1(2-163):<br>MGLNDIFEAQKIEWHEGGSPSIKLQSSDGEIFEVDVEIAKQSVTIKTMLEDLGMDDEGDDDPVPLP<br>NVNAAILKKVIQWCTHHKDDPPPPEDDENKEKRTDDIPVWDQEFLKVDQGTLFELILAANYLDIKGL<br>LDVTCKTVANMIKGKTPEEIRKTFNIKNDFTEEEEAQVRKENQWCEEK<br><br>His$_{10}$-TEV-lin-FBXO22(12-403):<br>MASHHHHHHHHHHGSENLYFQGGSGSSVDPRSTFVLSNLAEVVERVLTFLPAKALLRVACVCRL<br>WRECVRRVLRTHRSVTWISAGLAEAGHLEGHCLVRVVAEELENVRILPHTVLYMADSETFISLEEC<br>RGHKRARKRTSMETALALEKLFPKQCQVLGIVTPGIVVTPMGSGSNRPQEIEIGESGFALLFPQIEG<br>IKIQPFHFIKDPKNLTLERHQLTEVGLLDNPELRVVLVFGYNCCKVGASNYLQQVVSTFSDMNIILA<br>GGQVDNLSSLTSEKNPLDIDASGVVGLSFSGHRIQSATVLLNEDVSDEKTAEAAMQRLKAANIPEH<br>NTIGFMFACVGRGFQYYRAKGNVEADAFRKFFPSVPLFGFFGNGEIGCDRIVTGNFILRKCNEVKD<br>DDLFHSYTTIMALIHLGSSK |
| **SKP1-FBXO22** | P63208/Q8NEZ5 — His$_{10}$-TEV-lin on FBXO22<br>SKP1(1-163):<br>MPSIKLQSSDGEIFEVDVEIAKQSVTIKTMLEDLGMDDEGDDDPVPLPNVNAAILKKVIQWCTHHKD<br>DPPPPEDDENKEKRTDDIPVWDQEFLKVDQGTLFELILAANYLDIKGLLDVTCKTVANMIKGKTPEE<br>IRKTFNIKNDFTEEEEAQVRKENQWCEEK<br><br>His$_{10}$-TEV-lin-FBXO22(12-403):<br>MASHHHHHHHHHHGSENLYFQGGSGSSVDPRSTFVLSNLAEVVERVLTFLPAKALLRVACVCRL<br>WRECVRRVLRTHRSVTWISAGLAEAGHLEGHCLVRVVAEELENVRILPHTVLYMADSETFISLEEC<br>RGHKRARKRTSMETALALEKLFPKQCQVLGIVTPGIVVTPMGSGSNRPQEIEIGESGFALLFPQIEG<br>IKIQPFHFIKDPKNLTLERHQLTEVGLLDNPELRVVLVFGYNCCKVGASNYLQQVVSTFSDMNIILA<br>GGQVDNLSSLTSEKNPLDIDASGVVGLSFSGHRIQSATVLLNEDVSDEKTAEAAMQRLKAANIPEH<br>NTIGFMFACVGRGFQYYRAKGNVEADAFRKFFPSVPLFGFFGNGEIGCDRIVTGNFILRKCNEVKD<br>DDLFHSYTTIMALIHLGSSK |
| **SKP1-FBXO22 R376D** | P63208/Q8NEZ5 — His$_{10}$-TEV-lin on FBXO22<br>SKP1(1-163):<br>MPSIKLQSSDGEIFEVDVEIAKQSVTIKTMLEDLGMDDEGDDDPVPLPNVNAAILKKVIQWCTHHKD<br>DPPPPEDDENKEKRTDDIPVWDQEFLKVDQGTLFELILAANYLDIKGLLDVTCKTVANMIKGKTPEE<br>IRKTFNIKNDFTEEEEAQVRKENQWCEEK<br><br>His$_{10}$-TEV-lin-FBXO22(12-403_**R376D**):<br>MASHHHHHHHHHHGSENLYFQGGSGSSVDPRSTFVLSNLAEVVERVLTFLPAKALLRVACVCRL<br>WRECVRRVLRTHRSVTWISAGLAEAGHLEGHCLVRVVAEELENVRILPHTVLYMADSETFISLEEC<br>RGHKRARKRTSMETALALEKLFPKQCQVLGIVTPGIVVTPMGSGSNRPQEIEIGESGFALLFPQIEG<br>IKIQPFHFIKDPKNLTLERHQLTEVGLLDNPELRVVLVFGYNCCKVGASNYLQQVVSTFSDMNIILA<br>GGQVDNLSSLTSEKNPLDIDASGVVGLSFSGHRIQSATVLLNEDVSDEKTAEAAMQRLKAANIPEH<br>NTIGFMFACVGRGFQYYRAKGNVEADAFRKFFPSVPLFGFFGNGEIGCDRIVTGNFIL**D**KCNEVKD<br>DDLFHSYTTIMALIHLGSSK |
| **SKP1-FBXO22 R376P** | P63208/Q8NEZ5 — His$_{10}$-TEV-lin on FBXO22<br>SKP1(1-163):<br>MPSIKLQSSDGEIFEVDVEIAKQSVTIKTMLEDLGMDDEGDDDPVPLPNVNAAILKKVIQWCTHHKD<br>DPPPPEDDENKEKRTDDIPVWDQEFLKVDQGTLFELILAANYLDIKGLLDVTCKTVANMIKGKTPEE<br>IRKTFNIKNDFTEEEEAQVRKENQWCEEK<br><br>His$_{10}$-TEV-lin-FBXO22(12-403_**R376P**):<br>MASHHHHHHHHHHGSENLYFQGGSGSSVDPRSTFVLSNLAEVVERVLTFLPAKALLRVACVCRL<br>WRECVRRVLRTHRSVTWISAGLAEAGHLEGHCLVRVVAEELENVRILPHTVLYMADSETFISLEEC<br>RGHKRARKRTSMETALALEKLFPKQCQVLGIVTPGIVVTPMGSGSNRPQEIEIGESGFALLFPQIEG<br>IKIQPFHFIKDPKNLTLERHQLTEVGLLDNPELRVVLVFGYNCCKVGASNYLQQVVSTFSDMNIILA<br>GGQVDNLSSLTSEKNPLDIDASGVVGLSFSGHRIQSATVLLNEDVSDEKTAEAAMQRLKAANIPEH<br>NTIGFMFACVGRGFQYYRAKGNVEADAFRKFFPSVPLFGFFGNGEIGCDRIVTGNFIL**P**KCNEVKD<br>DDLFHSYTTIMALIHLGSSK |
| **SKP1-FBXO22 Q307E** | P63208/Q8NEZ5 — His$_{10}$-TEV-lin on FBXO22<br>SKP1(1-163):<br>MPSIKLQSSDGEIFEVDVEIAKQSVTIKTMLEDLGMDDEGDDDPVPLPNVNAAILKKVIQWCTHHKD<br>DPPPPEDDENKEKRTDDIPVWDQEFLKVDQGTLFELILAANYLDIKGLLDVTCKTVANMIKGKTPEE<br>IRKTFNIKNDFTEEEEAQVRKENQWCEEK |

| | |
|---|---|
| | His$_{10}$-TEV-lin-FBXO22(12-403_**Q307E**):<br>MASHHHHHHHHHHGSENLYFQGGSGSSVDPRSTFVLSNLAEVVERVLTFLPAKALLRVACVCRL<br>WRECVRRVLRTHRSVTWISAGLAEAGHLEGHCLVRVVAEELENVRILPHTVLYMADSETFISLEEC<br>RGHKRARKRTSMETALALEKLFPKQCQVLGIVTPGIVVTPMGSGSNRPQEIEIGESGFALLFPQIEG<br>IKIQPFHFIKDPKNLTLERHQLTEVGLLDNPELRVVLVFGYNCCKVGASNYLQQVVSTFSDMNIILA<br>GGQVDNLSSLTSEKNPLDIDASGVVGLSFSGHRIQSATVLLNEDVSDEKTAEAAM**E**RLKAANIPEH<br>NTIGFMFACVGRGFQYYRAKGNVEADAFRKFFPSVPLFGFFGNGEIGCDRIVTGNFILRKCNEVKD<br>DDLFHSYTTIMALIHLGSSK |
| **SKP1-FBXO22 Q307R** | P63208/Q8NEZ5 \| His$_{10}$-TEV-lin on FBXO22<br>SKP1(1-163):<br>MPSIKLQSSDGEIFEVDVEIAKQSVTIKTMLEDLGMDDEGDDDPVPLPNVNAAILKKVIQWCTHHKD<br>DPPPPEDDENKEKRTDDIPVWDQEFLKVDQGTLFELILAANYLDIKGLLDVTCKTVANMIKGKTPEE<br>IRKTFNIKNDFTEEEEAQVRKENQWCEEK<br><br>His$_{10}$-TEV-lin-FBXO22(12-403_**Q307R**):<br>MASHHHHHHHHHHGSENLYFQGGSGSSVDPRSTFVLSNLAEVVERVLTFLPAKALLRVACVCRL<br>WRECVRRVLRTHRSVTWISAGLAEAGHLEGHCLVRVVAEELENVRILPHTVLYMADSETFISLEEC<br>RGHKRARKRTSMETALALEKLFPKQCQVLGIVTPGIVVTPMGSGSNRPQEIEIGESGFALLFPQIEG<br>IKIQPFHFIKDPKNLTLERHQLTEVGLLDNPELRVVLVFGYNCCKVGASNYLQQVVSTFSDMNIILA<br>GGQVDNLSSLTSEKNPLDIDASGVVGLSFSGHRIQSATVLLNEDVSDEKTAEAAM**R**RLKAANIPEH<br>NTIGFMFACVGRGFQYYRAKGNVEADAFRKFFPSVPLFGFFGNGEIGCDRIVTGNFILRKCNEVKD<br>DDLFHSYTTIMALIHLGSSK |
| **SKP1-FBXO22 R367W** | P63208/Q8NEZ5 \| His$_{10}$-TEV-lin on FBXO22<br>SKP1(1-163):<br>MPSIKLQSSDGEIFEVDVEIAKQSVTIKTMLEDLGMDDEGDDDPVPLPNVNAAILKKVIQWCTHHKD<br>DPPPPEDDENKEKRTDDIPVWDQEFLKVDQGTLFELILAANYLDIKGLLDVTCKTVANMIKGKTPEE<br>IRKTFNIKNDFTEEEEAQVRKENQWCEEK<br><br>His$_{10}$-TEV-lin-FBXO22(12-403_**R367W**):<br>MASHHHHHHHHHHGSENLYFQGGSGSSVDPRSTFVLSNLAEVVERVLTFLPAKALLRVACVCRL<br>WRECVRRVLRTHRSVTWISAGLAEAGHLEGHCLVRVVAEELENVRILPHTVLYMADSETFISLEEC<br>RGHKRARKRTSMETALALEKLFPKQCQVLGIVTPGIVVTPMGSGSNRPQEIEIGESGFALLFPQIEG<br>IKIQPFHFIKDPKNLTLERHQLTEVGLLDNPELRVVLVFGYNCCKVGASNYLQQVVSTFSDMNIILA<br>GGQVDNLSSLTSEKNPLDIDASGVVGLSFSGHRIQSATVLLNEDVSDEKTAEAAMQRLKAANIPEH<br>NTIGFMFACVGRGFQYYRAKGNVEADAFRKFFPSVPLFGFFGNGEIGCD**W**IVTGNFILRKCNEVK<br>DDDLFHSYTTIMALIHLGSSK |
| **SKP1-FBXO22 R367L** | P63208/Q8NEZ5 \| His$_{10}$-TEV-lin on FBXO22<br>SKP1(1-163):<br>MPSIKLQSSDGEIFEVDVEIAKQSVTIKTMLEDLGMDDEGDDDPVPLPNVNAAILKKVIQWCTHHKD<br>DPPPPEDDENKEKRTDDIPVWDQEFLKVDQGTLFELILAANYLDIKGLLDVTCKTVANMIKGKTPEE<br>IRKTFNIKNDFTEEEEAQVRKENQWCEEK<br><br>His$_{10}$-TEV-lin-FBXO22(12-403_**R367L**):<br>MASHHHHHHHHHHGSENLYFQGGSGSSVDPRSTFVLSNLAEVVERVLTFLPAKALLRVACVCRL<br>WRECVRRVLRTHRSVTWISAGLAEAGHLEGHCLVRVVAEELENVRILPHTVLYMADSETFISLEEC<br>RGHKRARKRTSMETALALEKLFPKQCQVLGIVTPGIVVTPMGSGSNRPQEIEIGESGFALLFPQIEG<br>IKIQPFHFIKDPKNLTLERHQLTEVGLLDNPELRVVLVFGYNCCKVGASNYLQQVVSTFSDMNIILA<br>GGQVDNLSSLTSEKNPLDIDASGVVGLSFSGHRIQSATVLLNEDVSDEKTAEAAMQRLKAANIPEH<br>NTIGFMFACVGRGFQYYRAKGNVEADAFRKFFPSVPLFGFFGNGEIGCD**L**IVTGNFILRKCNEVKD<br>DDLFHSYTTIMALIHLGSSK |
| **$^{Avi}$SKP1-FBXL17** | P63208/Q9UF56 \| Avi-lin on SKP1, His$_{10}$-MBP-lin-TEV on FBXL17<br>Avi- lin-SKP1:<br>MGLNDIFEAQKIEWHEGGSPSIKLQSSDGEIFEVDVEIAKQSVTIKTMLEDLGMDDEGDDDPVPLP<br>NVNAAILKKVIQWCTHHKDDPPPPEDDENKEKRTDDIPVWDQEFLKVDQGTLFELILAANYLDIKGL<br>LDVTCKTVANMIKGKTPEEIRKTFNIKNDFTEEEEAQVRKENQWCEEK<br><br>His$_{10}$-MBP-lin-TEV-FBXL17(310-701):<br>MASHHHHHHHHHHGSKIEEGKLVIWINGDKGYNGLAEVGKKFEKDTGIKVTVEHPDKLEEKFPQV<br>AATGDGPDIIFWAHDRFGGYAQSGLLAEITPDKAFQDKLYPFTWDAVRYNGKLIAYPIAVEALSLIY<br>NKDLLPNPPKTWEEIPALDKELKAKGKSALMFNLQEPYFTWPLIAADGGYAFKYENGKYDIKDVGV<br>DNAGAKAGLTFLVDLIKNKHMNADTDYSIAEAAFNKGETAMTINGPWAWSNIDTSKVNYGVTVLPT<br>FKGQPSKPFVGVLSAGINAASPNKELAKEFLENYLLTDEGLEAVNKDKPLGAVALKSYEEELAKDP<br>RIAATMENAQKGEIMPNIPQMSAFWYAVRTAVINAASGRQTVDEALKDAQTRITKGGGGSENLYF<br>QSCHREPPPETPDINQLPPSILLKIFSNLSLDERCLSASLVCKYWRDLCLDFQFWKQLDLSSRQQV<br>TDELLEKIASRSQNIIEINISDCRSMSDNGVCVLAFKCPGLLRYTAYRCKQLSDTSIIAVASHCPLLQ<br>KVHVGNQDKLTDEGLKQLGSKCRELKDIHFGQCYKISDEGMIVIAKGCLKLQRIYMQENKLVTDQS |

| | | |
|---|---|---|
| | VKAFAEHCPELQYVGFMGCSVTSKGVIHLTKLRNLSSLDLRHITELDNETVMEIVKRCKNLSSLNLC<br>LNWIINDRCVEVIAKEGQNLKELYLVSCKITDYALIAIGRYSMTIETVDVGWCKEITDQGATLIAQSSK<br>SLRYLGLMRCDKVNEVTVEQLVQQYPHITFSTVLQDCKRTLERAYQMGWTPNMSAASS | |
| **BACH1$^{FL}$** | O14867 | His$_{10}$-ZZ-lin-HRV3C |
| | His$_{10}$-ZZ-lin-HRV3C-BACH1(2-736):<br>MASHHHHHHHHHHAQHDEAVDNKFNKEQQNAFYEILHLPNLNEEQRNAFIQSLKDDPSQSANLL<br>AEAKKLNDAQAPKVDNKFNKEQQNAFYEILHLPNLNEEQRNAFIQSLKDDPSQSANLLAEAKKLND<br>AQAPKVDANGGGGSGGGGSLEVLFQGPSLSENSVFAYESSVHSTNVLLSLNDQRKKDVLCDVTI<br>FVEGQRFRAHRSVLAACSSYFHSRIVGQADGELNITLPEEVTVKGFEPLIQFAYTAKLILSKENVDE<br>VCKCVEFLSVHNIEESCFQFLKFKFLDSTADQQECPRKKCFSSHCQKTDLKLSLLDQRDLETDEVE<br>EFLENKNVQTPQCKLRRYQGNAKASPPLQDSASQTYESMCLEKDAALALPSLCPKYRKFQKAFG<br>TDRVRTGESSVKDIHASVQPNERSENECLGGVPECRDLQVMLKCDESKLAMEPEETKKDPASQC<br>PTEKSEVTPFPHNSSIDPHGLYSLSLLHTYDQYGDLNFAGMQNTTVLTEKPLSGTDVQEKTFGES<br>QDLPLKSDLGTREDSSVASSDRSSVEREVAEHLAKGFWSDICSTDTPCQMQLSPAVAKDGSEQIS<br>QKRSECPWLGIRISESPEPGQRTFTTLSSVNCPFISTLSTEGCSSNLEIGNDDYVSEPQQEPCPYA<br>CVISLGDDSETDTEGDSESCSAREQECEVKLPFNAQRIISLSRNDFQSLLKMHKLTPEQLDCIHDIR<br>RRSKNRIAAQRCRKRKLDCIQNLESEIEKLQSEKESLLKERDHILSTLGETKQNLTGLCQKVCKEAA<br>LSQEQIQILAKYSAADCPLSFLISEKDKSTPDGELALPSIFSLSDRPPAVLPPCARGNSEPGYARGQ<br>ESQQMSTATSEQAGPAEQCRQSGGISDFCQQMTDKCTTDE | |
| **BACH1$^{\Delta CP}$** | O14867 | His$_{10}$-ZZ-lin-HRV3C |
| | His$_{10}$-ZZ-lin-HRV3C-BACH1(2-736<br>_C224A_C299A_C435A_C461A_C492A_C646A):<br>MASHHHHHHHHHHAQHDEAVDNKFNKEQQNAFYEILHLPNLNEEQRNAFIQSLKDDPSQSANLL<br>AEAKKLNDAQAPKVDNKFNKEQQNAFYEILHLPNLNEEQRNAFIQSLKDDPSQSANLLAEAKKLND<br>AQAPKVDANGGGGSGGGGSLEVLFQGPSLSENSVFAYESSVHSTNVLLSLNDQRKKDVLCDVTI<br>FVEGQRFRAHRSVLAACSSYFHSRIVGQADGELNITLPEEVTVKGFEPLIQFAYTAKLILSKENVDE<br>VCKCVEFLSVHNIEESCFQFLKFKFLDSTADQQECPRKKCFSSHCQKTDLKLSLLDQRDLETDEVE<br>EFLENKNVQTPQCKLRRYQGNAKASPPLQDSASQTYESMCLEKDAALALPSLCPKYRKFQKAFG<br>TDRVRTGESSVKDIHASVQPNERSENECLGGVPECRDLQVMLKCDESKLAMEPEETKKDPASQC<br>PTEKSEVTPFPHNSSIDPHGLYSLSLLHTYDQYGDLNFAGMQNTTVLTEKPLSGTDVQEKTFGES<br>QDLPLKSDLGTREDSSVASSDRSSVEREVAEHLAKGFWSDICSTDTPCQMQLSPAVAKDGSEQIS<br>QKRSECPWLGIRISESPEPGQRTFTTLSSVNCPFISTLSTEGCSSNLEIGNDDYVSEPQQEPCPYA<br>CVISLGDDSETDTEGDSESCSAREQECEVKLPFNAQRIISLSRNDFQSLLKMHKLTPEQLDCIHDIR<br>RRSKNRIAAQRCRKRKLDCIQNLESEIEKLQSEKESLLKERDHILSTLGETKQNLTGLCQKVCKEAA<br>LSQEQIQILAKYSAADCPLSFLISEKDKSTPDGELALPSIFSLSDRPPAVLPPCARGNSEPGYARGQ<br>ESQQMSTATSEQAGPAEQCRQSGGISDFCQQMTDKCTTDE | |
| **BACH1$^{BTB}$** | O14867 | His$_{10}$-TEV |
| | His$_{6}$-TEV-BACH1(7-128)<br>MHHHHHHSSGVDLGTENLYFQSMSVFAYESSVHSTNVLLSLNDQRKKDVLCDVTIFVEGQRFRA<br>HRSVLAACSSYFHSRIVGQADGELNITLPEEVTVKGFEPLIQFAYTAKLILSKENVDEVCKCVEFLSV<br>HNIEESCFQFLKF | |
| **BACH1$^{BTB}$ F9A** | His$_{6}$-TEV-BACH1(7-128_**F9A**)<br>MHHHHHHSSGVDLGTENLYFQSMSV**A**AYESSVHSTNVLLSLNDQRKKDVLCDVTIFVEGQRFRA<br>HRSVLAACSSYFHSRIVGQADGELNITLPEEVTVKGFEPLIQFAYTAKLILSKENVDEVCKCVEFLSV<br>HNIEESCFQFLKF | |
| **BACH1$^{BTB}$ Y11A** | His$_{6}$-TEV-BACH1(7-128_**Y11A**)<br>MHHHHHHSSGVDLGTENLYFQSMSVFA**A**ESSVHSTNVLLSLNDQRKKDVLCDVTIFVEGQRFRA<br>HRSVLAACSSYFHSRIVGQADGELNITLPEEVTVKGFEPLIQFAYTAKLILSKENVDEVCKCVEFLSV<br>HNIEESCFQFLKF | |
| **BACH1$^{BTB}$ Y11F** | His$_{6}$-TEV-BACH1(7-128_**Y11F**)<br>MHHHHHHSSGVDLGTENLYFQSMSVFA**F**ESSVHSTNVLLSLNDQRKKDVLCDVTIFVEGQRFRA<br>HRSVLAACSSYFHSRIVGQADGELNITLPEEVTVKGFEPLIQFAYTAKLILSKENVDEVCKCVEFLSV<br>HNIEESCFQFLKF | |
| **BACH1$^{BTB}$ Y11H** | His$_{6}$-TEV-BACH1(7-128_**Y11H**)<br>MHHHHHHSSGVDLGTENLYFQSMSVFA**H**ESSVHSTNVLLSLNDQRKKDVLCDVTIFVEGQRFRA<br>HRSVLAACSSYFHSRIVGQADGELNITLPEEVTVKGFEPLIQFAYTAKLILSKENVDEVCKCVEFLSV<br>HNIEESCFQFLKF | |
| **BACH1$^{BTB}$ E12R** | His$_{6}$-TEV-BACH1(7-128_**E12R**)<br>MHHHHHHSSGVDLGTENLYFQSMSVFAY**R**SSVHSTNVLLSLNDQRKKDVLCDVTIFVEGQRFRA<br>HRSVLAACSSYFHSRIVGQADGELNITLPEEVTVKGFEPLIQFAYTAKLILSKENVDEVCKCVEFLSV<br>HNIEESCFQFLKF | |
| **BACH1$^{BTB}$ S13A** | His$_{6}$-TEV-BACH1(7-128_**S13A**)<br>MHHHHHHSSGVDLGTENLYFQSMSVFAYEASVHSTNVLLSLNDQRKKDVLCDVTIFVEGQRFRA<br>HRSVLAACSSYFHSRIVGQADGELNITLPEEVTVKGFEPLIQFAYTAKLILSKENVDEVCKCVEFLSV<br>HNIEESCFQFLKF | |
| **BACH1$^{BTB}$ S13D** | His$_{6}$-TEV-BACH1(7-128_**S13D**) | |

| | |
|---|---|
| | MHHHHHHSSGVDLGTENLYFQSMSVFAYE**D**SVHSTNVLLSLNDQRKKDVLCDVTIFVEGQRFRAHRSVLAACSSYFHSRIVGQADGELNITLPEEVTVKGFEPLIQFAYTAKLILSKENVDEVCKCVEFLSVHNIEESCFQFLKF |
| **BACH1$^{\Delta BTB}$** | O14867 — His$_6$-ZZ-lin-HRV3C |
| | His$_6$-ZZ-lin-HRV3C-BACH1(174-736)<br>MKTHHHHHHGAQHDEAVDNKFNKEQQNAFYEILHLPNLNEEQRNAFIQSLKDDPSQSANLLAEAKKLNDAQAPKVDNKFNKEQQNAFYEILHLPNLNEEQRNAFIQSLKDDPSQSANLLAEAKKLNDAQAPKVDANGGGSGGGSGGGSLEVLFQGPLENKNVQTPQCKLRRYQGNAKASPPLQDSASQTYESMCLEKDAALALPSLCPKYRKFQKAFGTDRVRTGESSVKDIHASVQPNERSENECLGGVPECRDLQVMLKCDESKLAMEPEETKKDPASQCPTEKSEVTPFPHNSSIDPHGLYSLSLLHTYDQYGDLNFAGMQNTTVLTEKPLSGTDVQEKTFGESQDLPLKSDLGTREDSSVASSDRSSVEREVAEHLAKGFWSDICSTDTPCQMQLSPAVAKDGSEQISQKRSECPWLGIRISESPEPGQRTFTTLSSVNCPFISTLSTEGCSSNLEIGNDDYVSEPQQEPCPYACVISLGDDSETDTEGDSESCSAREQECEVKLPFNAQRIISLSRNDFQSLLKMHKLTPEQLDCIHDIRRRSKNRIAAQRCRKRKLDCIQNLESEIEKLQSEKESLLKERDHILSTLGETKQNLTGLCQKVCKEAALSQEQIQILAKYSAADCPLSFLISEKDKSTPDGELALPSIFSLSDRPPAVLPPCARGNSEPGYARGQESQQMSTATSEQAGPAEQCRQSGGISDFCQQMTDKCTTDE |
| **BACH1$^{\Delta C178}$** | O14867 — His$_6$-TEV- |
| | His$_6$-TEV-BACH1(7-177)<br>MHHHHHHSSGVDLGTENLYFQSMSVFAYESSVHSTNVLLSLNDQRKKDVLCDVTIFVEGQRFRAHRSVLAACSSYFHSRIVGQADGELNITLPEEVTVKGFEPLIQFAYTAKLILSKENVDEVCKCVEFLSVHNIEESCFQFLKFKFLDSTADQQECPRKKCFSSHCQKTDLKLSLLDQRDLETDEVEEFLENK |
| **BACH1$^{\Delta C241}$** | O14867 — His$_6$-TEV- |
| | His$_6$-TEV-BACH1(2-240)<br>MHHHHHHSSGVDLGTENLYFQSLSENSVFAYESSVHSTNVLLSLNDQRKKDVLCDVTIFVEGQRFRAHRSVLAACSSYFHSRIVGQADGELNITLPEEVTVKGFEPLIQFAYTAKLILSKENVDEVCKCVEFLSVHNIEESCFQFLKFKFLDSTADQQECPRKKCFSSHCQKTDLKLSLLDQRDLETDEVEEFLENKNVQTPQCKLRRYQGNAKASPPLQDSASQTYESMCLEKDAALALPSLCPKYRKFQKAFGTDRVR |
| **BACH1$^{\Delta C506}$** | O14867 — His$_6$-TEV- |
| | His$_6$-TEV-BACH1(2-505)<br>MHHHHHHSSGVDLGTENLYFQSLSENSVFAYESSVHSTNVLLSLNDQRKKDVLCDVTIFVEGQRFRAHRSVLAACSSYFHSRIVGQADGELNITLPEEVTVKGFEPLIQFAYTAKLILSKENVDEVCKCVEFLSVHNIEESCFQFLKFKFLDSTADQQECPRKKCFSSHCQKTDLKLSLLDQRDLETDEVEEFLENKNVQTPQCKLRRYQGNAKASPPLQDSASQTYESMCLEKDAALALPSLCPKYRKFQKAFGTDRVRTGESSVKDIHASVQPNERSENECLGGVPECRDLQVMLKCDESKLAMEPEETKKDPASQCPTEKSEVTPFPHNSSIDPHGLYSLSLLHTYDQYGDLNFAGMQNTTVLTEKPLSGTDVQEKTFGESQDLPLKSDLGTREDSSVASSDRSSVEREVAEHLAKGFWSDICSTDTPCQMQLSPAVAKDGSEQISQKRSECPWLGIRISESPEPGQRTFTTLSSVNCPFISTLSTEGCSSNLEIGNDDYVSEPQQEPCPYACVISLGDDSE |
| **BACH2$^{BTB}$** | Q9BYV9 — His$_{10}$-TEV- |
| | His$_{10}$-TEV-BACH2(8-131):<br>MKTHHHHHHHHHHGSENLYFQGDSPMYVYESTVHCTNILLGLNDQRKKDILCDVTLIVERKEFRAHRAVLAACSEYFWQALVGQTKNDLVVSLPEEVTARGFGPLLQFAYTAKLLLSRENIREVIRCAEFLRMHNLEDSCFSFLQT |
| **KEAP1$^{BTB}$** | Q14145 — His$_{10}$-TEV- |
| | His$_{10}$-TEV-KEAP1(48-180):<br>MKTHHHHHHHHHHGSENLYFQGNRTFSYTLEDHTKQAFGIMNELRLSQQLCDVTLQVKYQDAPAAQFMAHKVVLASSSPVFKAMFTNGLREQGMEVVSIEGIHPKVMERLIEFAYTASISMGEKCVLHVMNGAVMYQIDSVVRACSDFLVQQLD |
| **ZBTB16$^{BTB}$** | Q05516 — His$_{10}$-TEV- |
| | His$_{10}$-TEV-ZBTB16(1-130):<br>MKTHHHHHHHHHHGSENLYFQGMDLTKMGMIQLQNPSHPTGLLCKANQMRLAGTLCDVVIMVDSQEFHAHRTVLACTSKMFEILFHRNSQHYTLDFLSPKTFQQILEYAYTATLQAKAEDLDDLLYAAEILEIEYLEEQCLKMLETIQASDD |
| **CUL1-RBX1** | Q13616<br>P62877 — TwinStrep-TEV on CUL1 |
| | TwinStrep-TEV-CUL1(2-776):<br>MASAWSHPQFEKGGGSGGGSGGSAWSHPQFEKENLYFQSSTRSQNPHGLKQIGLDQIWDDLRAGIQQVYTRQSMAKSRYMELYTHVYNYCTSVHQSNQARGAGVPPSKSKKGQTPGGAQFVGLELYKRLKEFLKNYLTNLLKDGEDLMDESVLKFYTQQWEDYRFSSKVLNGICAYLNRHWVRRECDEGRKGIYEIYSLALVTWRDCLFRPLNKQVTNAVLKLIEKERNGETINTRLISGVVQSYVELGLNEDDAFAKGPTLTVYKESFESQFLADTERFYTRESTEFLQQNPVTEYMKKAEARLLEEQRRVQVYLHESTQDELARKCEQVLIEKHLEIFHTEFQNLLDADKNEDLGRMYNLVSRIQDGLGELKKLLETHIHNQGLAAIEKCGEAALNDPKMYVQTVLDVHKKYNALVMSAFNNDAGFVAALDKACGRFINNAVTKMAQSSSKSPELLARYCDSLLKKSSKNPEEAELEDTLNQVMVVFKYIEDKDVFQKFYAKMLAKRLVHQNSASD |

| | |
|---|---|
| | DAEASMISKLKQACGFEYTSKLQRMFQDIGVSKDLNEQFKKHLTNSEPLDLDFSIQVLSSGSWPF<br>QQSCTFALPSELERSYQRFTAFYASRHSGRKLTWLYQLSKGELVTNCFKNRYTLQASTFQMAILL<br>QYNTEDAYTVQQLTDSTQIKMDILAQVLQILLKSKLLVLEDENANVDEVELKPDTLIKLYLGYKNKKL<br>RVNINVPMKTEQKQEQETTHKNIEEDRKLLIQAAIVRIMKMRKVLKHQQLLGEVLTQLSSRFKPRVP<br>VIKKCIDILIEKEYLERVDGEKDTYSYLA<br><br>RBX1(1-108):<br>MAAAMDVDTPSGTNSGAGKKRFEVKKWNAVALWAWDIVVDNCAICRNHIMDLCIECQANQASAT<br>SEECTVAWGVCNHAFHFHCISRWLKTRQVCPLDNREWEFQKYGH |
| **Nedd8** | Q15843 ┆ ZZ-His$_6$-HRV3C<br>ZZ-His$_6$-S-HRV3C-Nedd8(2-76_**M1C**)<br>AQHDEAVDNKFNKEQQNAFYEILHLPNLNEEQRNAFIQSLKDDPSQSANLLAEAKKLNDAQAPKV<br>DNKFNKEQQNAFYEILHLPNLNEEQRNAFIQSLKDDPSQSANLLAEAKKLNDAQAPKVDANSTSG<br>SGHHHHHHSAGKETAAAKFERQHMDSPDLGTGNITSLYKKAGLEVLFQGPGS**C**LIKVKTLTGKEIE<br>IDIEPTDKVERIKERVEEKEGIPPQQQRLIYSGKQMNDEKTAADYKILGGSVLHLVLALRGG |
| **Cys-Ub** | P0CG47 ┆ His$_6$-lin-HRV3C-<br>His$_6$-lin-HRV3C-Ub(2-76_**M1C**):<br>MKTHHHHHHGGGGSGGGGSLEVLFQGPG**C**QIFVKTLTGKTITLEVEPSDTIENVKAKIQDKEGIPP<br>DQQRLIFAGKQLEDGRTLSDYNIQKESTLHLVLRLRGG |
| **Cys-Ub$^0$** | P0CG47 ┆ His$_6$-lin-HRV3C-<br>His$_6$-lin-HRV3C-Ub(2-76_**M1C**_K6R_K11R_K27R_K29R_K33R_K48R_K63R):<br>MKTHHHHHHGGGGSGGGGSLEVLFQGPG**C**QIFVRTLTGRTITLEVEPSDTIENVRARIQDREGIP<br>PDQQRLIFAGRQLEDGRTLSDYNIQRESTLHLVLR LRGG |

**Supplementary table 2: Kinetic binding parameter of the SKP1-FBXO22 interaction with BACH1 determined from SPR and BLI experiments.**

| Technique | Ligand | Analyte | $K_D$ (nM) | $k_a$ ($M^{-1}s^{-1}$) | $k_d$ ($s^{-1}$) |
|---|---|---|---|---|---|
| SPR | $^{Avi}$SKP1-FBXO22 | BACH1$^{BTB}$ | 57.7 | 51190 | 0.002953 |
| BLI | $^{Avi}$SKP1-FBXO22 | BACH1$^{BTB}$ | 123.8 | 27360 | 0.003387 |
| BLI | $^{Avi}$SKP1-FBXO22 | BACH1$^{BTB}$/hemin | 83.9 | 28330 | 0.002378 |
| BLI | $^{Avi}$SKP1-FBXO22 | BACH1$^{FL}$ | 44.3 | 47180 | 0.002088 |
| BLI | $^{Avi}$SKP1-FBXO22 | BACH1$^{FL}$/hemin | 28.6 | 59890 | 0.001714 |
| BLI* | $^{Avi}$SKP1-FBXO22 | BACH1$^{BTB}$ | 218.3 | 20230 | 0.004416 |
| BLI* | $^{Avi}$SKP1-FBXO22 | BACH1$^{BTB(BTFA)}$ | 5017 | 4437 | 0.0226 |
| BLI* | $^{Avi}$SKP1-FBXL17 | BACH1$^{BTB}$ | n/a | n/a | n/a |
| BLI* | $^{Avi}$SKP1-FBXL17 | BACH1$^{BTB(BTFA)}$ | 872.6 | 3249 | 0.002835 |

*single concentration for qualitative analysis

**Supplementary table 3: Fit parameter of the TR-FRET competition assay shown in Figure 1h using BACH1$^{BTB}$ wildtype (WT) and mutants as competitor.**

| log(inhibitor) vs. response -- Variable slope (four parameters) | **WT** | **F9A** | **Y11A** | **Y11F** | **Y11H** | **E12R** | **S13A** | **S13D** | **F125Y** | **F125A** |
|---|---|---|---|---|---|---|---|---|---|---|
| Best-fit values | | | | | | | | | | |
| Bottom | = 0.000 | = 0.000 | = 0.000 | = 0.000 | = 0.000 | = 0.000 | = 0.000 | = 0.000 | = 0.000 | = 0.000 |
| Top | = 1.000 | = 1.000 | = 1.000 | = 1.000 | = 1.000 | = 1.000 | = 1.000 | = 1.000 | = 1.000 | = 1.000 |
| $LogIC_{50}$ | -1.193 | -0.4944 | 0.4245 | -0.8564 | -0.06288 | -0.8246 | -0.7408 | 0.3915 | -0.1828 | 0.8728 |
| HillSlope | -1.009 | -1.245 | -1.525 | -1.243 | -2.043 | -0.8735 | -1.196 | -1.039 | -1.220 | -1.547 |
| $IC_{50}$ | 0.06419 | 0.3203 | 2.657 | 0.1392 | 0.8652 | 0.1498 | 0.1816 | 2.463 | 0.6564 | 7.461 |
| Span | = 1.000 | = 1.000 | = 1.000 | = 1.000 | = 1.000 | = 1.000 | = 1.000 | = 1.000 | = 1.000 | = 1.000 |
| 95% CI (profile likelihood) | | | | | | | | | | |
| $LogIC_{50}$ | -1.306 to -1.077 | -0.5810 to -0.4067 | 0.3462 to 0.5044 | -0.9912 to -0.7144 | -0.1267 to 0.001321 | -1.014 to -0.6247 | -0.8324 to -0.6465 | 0.3388 to 0.4445 | -0.2524 to -0.1124 | 0.7955 to 0.9515 |
| HillSlope | -1.277 to -0.8175 | -1.534 to -1.028 | -1.926 to -1.230 | -1.798 to -0.9126 | -2.583 to -1.629 | -1.257 to -0.6424 | -1.523 to -0.9671 | -1.171 to -0.9283 | -1.446 to -1.044 | -1.930 to -1.260 |
| $IC_{50}$ | 0.04948 to 0.08383 | 0.2624 to 0.3920 | 2.219 to 3.195 | 0.1020 to 0.1930 | 0.7470 to 1.003 | 0.09680 to 0.2373 | 0.1471 to 0.2257 | 2.182 to 2.783 | 0.5593 to 0.7720 | 6.244 to 8.943 |
| Goodness of Fit | | | | | | | | | | |
| Degrees of Freedom | 10 | 10 | 10 | 10 | 10 | 10 | 10 | 10 | 10 | 10 |
| R squared | 0.9912 | 0.9939 | 0.9918 | 0.9856 | 0.9948 | 0.9769 | 0.9932 | 0.9970 | 0.9957 | 0.9892 |
| Sum of Squares | 0.01958 | 0.01419 | 0.01407 | 0.03414 | 0.01209 | 0.04710 | 0.01535 | 0.004493 | 0.009012 | 0.01386 |
| Sy.x | 0.04425 | 0.03767 | 0.03751 | 0.05843 | 0.03477 | 0.06863 | 0.03917 | 0.02120 | 0.03002 | 0.03722 |
| Constraints | | | | | | | | | | |
| Bottom | Bottom = 0 | Bottom = 0 | Bottom = 0 | Bottom = 0 | Bottom = 0 | Bottom = 0 | Bottom = 0 | Bottom = 0 | Bottom = 0 | Bottom = 0 |
| Top | Top = 1 | Top = 1 | Top = 1 | Top = 1 | Top = 1 | Top = 1 | Top = 1 | Top = 1 | Top = 1 | Top = 1 |
| Number of points | | | | | | | | | | |
| # of X values | 12 | 12 | 12 | 12 | 12 | 12 | 12 | 12 | 12 | 12 |
| # Y values analyzed | 12 | 12 | 12 | 12 | 12 | 12 | 12 | 12 | 12 | 12 |

**Supplementary table 4: Fit parameter of the TR-FRET competition assay shown in Figure 1k using BACH2, KEAP1, and ZBTB16 BTB domains as competitor.**

| log(inhibitor) vs. response -- Variable slope (four parameters) | $BACH2^{BTB}$ | $KEAP1^{BTB}$ | $ZBTB16^{BTB}$ |
|---|---|---|---|
| Best-fit values | | | |
| Bottom | = 0.000 | = 0.000 | = 0.000 |
| Top | = 1.000 | = 1.000 | = 1.000 |
| LogIC50 | -1.072 | -0.8237 | -0.2069 |
| HillSlope | -1.151 | -1.104 | -1.323 |
| IC50 | 0.08468 | 0.1501 | 0.6211 |
| Span | = 1.000 | = 1.000 | = 1.000 |
| 95% CI (profile likelihood) | | | |
| LogIC50 | -1.127 to -1.018 | -0.8963 to -0.7506 | -0.2873 to -0.1279 |
| HillSlope | -1.316 to -1.015 | -1.308 to -0.9446 | -1.727 to -1.053 |
| IC50 | 0.07467 to 0.09594 | 0.1270 to 0.1776 | 0.5161 to 0.7449 |
| Goodness of Fit | | | |
| Degrees of Freedom | 10 | 10 | 10 |
| R squared | 0.9976 | 0.9959 | 0.9936 |
| Sum of Squares | 0.005246 | 0.008989 | 0.01305 |
| Sy.x | 0.02290 | 0.02998 | 0.03612 |
| Constraints | | | |
| Bottom | Bottom = 0 | Bottom = 0 | Bottom = 0 |
| Top | Top = 1 | Top = 1 | Top = 1 |
| Number of points | | | |
| # of X values | 12 | 12 | 12 |
| # Y values analyzed | 12 | 12 | 12 |

**Supplementary table 5: Fit parameter of the TR-FRET competition assay shown in Figure 3b and e using SKP1-FBXO22 wildtype (WT) and mutants as competitor.**

| log(inhibitor) vs. response -- Variable slope (four parameters) | **WT** | **R376P** | **R376D** | **Q307E** | **Q307R** | **R367W** | **R367L** |
|---|---|---|---|---|---|---|---|
| Best-fit values | | | | | | | |
| Bottom | = 0.000 | = 0.000 | = 0.000 | = 0.000 | = 0.000 | = 0.000 | = 0.000 |
| Top | = 1.000 | = 1.000 | = 1.000 | = 1.000 | = 1.000 | = 1.000 | = 1.000 |
| LogIC50 | -0.3041 | 1.131 | 0.9527 | -0.2403 | -0.7822 | 0.1696 | 0.01485 |
| HillSlope | -1.089 | -1.015 | -1.188 | -0.8795 | -0.8656 | -0.8642 | -0.8805 |
| IC50 | 0.4965 | 13.51 | 8.968 | 0.5750 | 0.1651 | 1.478 | 1.035 |
| Span | = 1.000 | = 1.000 | = 1.000 | = 1.000 | = 1.000 | = 1.000 | = 1.000 |
| 95% CI (profile likelihood) | | | | | | | |
| LogIC50 | -0.3835 to -0.2236 | 1.007 to 1.260 | 0.8566 to 1.051 | -0.3465 to -0.1323 | -0.8870 to -0.6755 | 0.1034 to 0.2362 | -0.06281 to 0.09321 |
| HillSlope | -1.307 to -0.9218 | -1.328 to -0.7943 | -1.531 to -0.9439 | -1.072 to -0.7346 | -1.050 to -0.7254 | -0.9808 to -0.7661 | -1.014 to -0.7707 |
| IC50 | 0.4135 to 0.5976 | 10.16 to 18.20 | 7.188 to 11.24 | 0.4503 to 0.7375 | 0.1297 to 0.2111 | 1.269 to 1.723 | 0.8653 to 1.239 |
| Goodness of Fit | | | | | | | |
| Degrees of Freedom | 10 | 10 | 10 | 10 | 10 | 9 | 9 |
| R squared | 0.9952 | 0.9783 | 0.9860 | 0.9922 | 0.9928 | 0.9959 | 0.9951 |
| Sum of Squares | 0.01059 | 0.02347 | 0.01696 | 0.01524 | 0.01461 | 0.005122 | 0.007109 |
| Sy.x | 0.03254 | 0.04845 | 0.04118 | 0.03903 | 0.03822 | 0.02386 | 0.02811 |
| Constraints | | | | | | | |
| Bottom | Bottom = 0 | Bottom = 0 | Bottom = 0 | Bottom = 0 | Bottom = 0 | Bottom = 0 | Bottom = 0 |
| Top | Top = 1 | Top = 1 | Top = 1 | Top = 1 | Top = 1 | Top = 1 | Top = 1 |
| Number of points | | | | | | | |
| # of X values | 12 | 12 | 12 | 12 | 12 | 11 | 11 |
| # Y values analyzed | 12 | 12 | 12 | 12 | 12 | 11 | 11 |

**Supplementary table 6: Overview of melting temperatures determined via nano DSF.**

| | | Fluorescence-based (350 nm / 33 nm) | | Turbidity-based | |
|---|---|---|---|---|---|
| **Protein** | **construct** | **Average $T_m$ (°C)** | **Standard deviation from n=3 experiments Average (°C)** | **Average $T_m$ (°C)** | **Standard deviation from n=3 experiments Average (°C)** |
| **SKP1-FBXO22** | Wildtype | 55.44 | 0.14 | 55.03 | 0.10 |
| | R376P | 54.20 | 0.11 | 54.15 | 0.12 |
| | R376D | 55.08 | 0.11 | 55.01 | 0.11 |
| | Q307E | 55.10 | 0.13 | 54.91 | 0.13 |
| | Q307R | 54.99 | 0.06 | 54.61 | 0.15 |
| | R367L | 49.41 | 0.11 | 50.17 | 0.06 |
| | R367W | 46.96 | 0.37 | 49.94 | 0.08 |
| **BACH1$^{BTB}$** | Wildtype | 58.23 | 0.53 | 63.34 | 0.69 |
| | F9A | 49.21 | 0.13 | 51.75 | 0.03 |
| | Y11A | 43.59 | 0.03 | 44.49 | 0.05 |
| | Y11F | 52.79 | 0.24 | 58.03 | 0.31 |
| | Y11H | 45.08 | 0.06 | 47.43 | 0.28 |
| | E12R | 51.74 | 0.06 | 57.07 | 0.07 |
| | S13A | 46.72 | 0.01 | 47.79 | 0.04 |
| | S13D | 48.29 | 0.03 | 49.37 | 0.05 |
| | A53V | 58.52 | 0.74 | 63.00 | 0.79 |
| | F125A | 56.48 | 0.23 | 63.81 | 0.11 |
| | F125Y | 58.41 | 0.49 | 61.81 | 0.49 |
| | BTFA | 58.04 | 0.14 | 59.44 | 0.30 |

**Supplementary table 7: Molecular weights and oligomeric states of protein constructs used in this study determined by SEC-MALS.**

| Protein | construct | Calculated mass (Da) | Experimental mass (Da) | Error (%) | Oligomeric state |
|---|---|---|---|---|---|
| **SKP1-FBXO22** | wildtype | 62184 | 61330 | 2.505 | Heterodimer |
| | R376P | 62125 | 58340 | 2.550 | Heterodimer |
| | R376D | 62143 | 57400 | 1.991 | Heterodimer |
| | Q307E | 62185 | 59340 | 1.250 | Heterodimer |
| | Q307R | 62212 | 58630 | 0.885 | Heterodimer |
| | R367L | 62141 | 59260 | 1.438 | Heterodimer |
| | R367W | 62214 | 60470 | 1.461 | Heterodimer |
| **SKP1-FBXL17** | Wildtype | 65191 | 65690 | 4.006 | Heterodimer |
| **BACH1** | FL | 81980 | 163100 | 0.517 | Homodimer |
| | ΔBTB | 62356 | 127700 | 2.643 | Homodimer |
| | ΔC177 | 19814 | 40590 | 3.260 | Homodimer |
| | ΔC240 | 27215 | 53110 | 3.553 | Homodimer |
| | ΔC505 | 56091 | 118400 | 3.248 | Homodimer |
| **BACH1$^{BTB}$** | wildtype | 14042 | 27420 | 5.068 | Homodimer |
| | F9A | 13966 | 27460 | 3.830 | Homodimer |
| | Y11A | 13950 | 30200 | 3.144 | Homodimer |
| | Y11F | 14026 | 28710 | 4.190 | Homodimer |
| | Y11H | 14016 | 31420 | 1.552 | Homodimer |
| | E12R | 14069 | 30220 | 1.008 | Homodimer |
| | S13A | 14026 | 27900 | 2.638 | Homodimer |
| | S13D | 14070 | 30070 | 2.848 | Homodimer |
| | A53V | 14070 | 27670 | 2.045 | Homodimer |
| | F125A | 13965 | 27280 | 3.046 | Homodimer |
| | F125Y | 14058 | 27930 | 2.661 | Homodimer |
| | BTFA | 14042 | 27500 | 1.300 | Homodimer |
| **BACH1** | BTB | 14331 | 30010 | 5.626 | Homodimer |
| **KEAP** | BTB | 14881 | 32720 | 4.363 | Homodimer |
| **ZBTB16** | BTB | 14948 | 28790 | 5.110 | Homodimer |
| **complex** | SKP1-FBXO22/ BACH1$^{FL}$ | | 189500 | 0.478 | Heterodimer + homodimer |

**Supplementary table 8: Cryo-electron microscopy data.**

| | SKP1-FBO22/BACH1$^{BTB}$ (EMDB-19766) (PDB-8S7D) | SKP1-FBXO22 (EMDB-19768) (PDB-8S7E) |
|---|---|---|
| **Data collection and processing** | | |
| Magnification | 75000 | 96000 |
| Voltage (kV) | 300 | 300 |
| Electron exposure ($e^-/Å^2$) | 50 | 50 |
| Defocus range (um) | 0.6-1.6 | 0.6-1.6 |
| Pixel size (Å) | 0.845 | 0.656 |
| Symmetry imposed | C1 | C1 |
| Initial particle images (no.) | 8434370 | 6062563 |
| Final particle images (no.) | 1715031 | 717992 |
| Map resolution (Å) | 3.2 | 3.4 |
| FSC threshold | 0.143 | 0.143 |
| Map resolution range (Å) | 3.0-3.8 | 3.0-3.8 |
| **Refinement** | | |
| Initial model used | AlphaFold2 prediction of SKP1-FBXO22 with BACH1$^{BTB}$ | AlphaFold2 prediction of SKP1-FBXO22 |
| Model resolution (Å) | 3.2 | 3.44 |
| FSC threshold | 0.143 | 0.143 |
| Model resolution range (Å) | 3.1-3.5 | 3.3-6.8 |
| Map sharpening B factor ($Å^2$) | 202 | 177s |
| Model composition | | |
| Non-hydrogen atoms | 5488 | 3664 |
| Protein residues | 691 | 466 |
| B factors ($Å^2$) | | |
| Protein (min/max/mean) | 0.00/178.14/83.60 | 69.94/126.43/96.78 |
| R.m.s. deviations | | |
| Bond lengths (Å) | 0.003 (0) | 0.003 (0) |
| Bond angles (°) | 0.569 (2) | 0.550 (10) |
| Validation | | |
| MolProbity score | 2.37 | 2.29 |
| Clashscore | 13.28 | 20.70 |
| Poor rotamers (%) | 4.7 | 0.25 |
| Ramachandran plot | | |
| Favored (%) | 96.54 | 92.19 |
| Allowed (%) | 3.46 | 7.81 |
| Disallowed (%) | 0.00 | 0.00 |

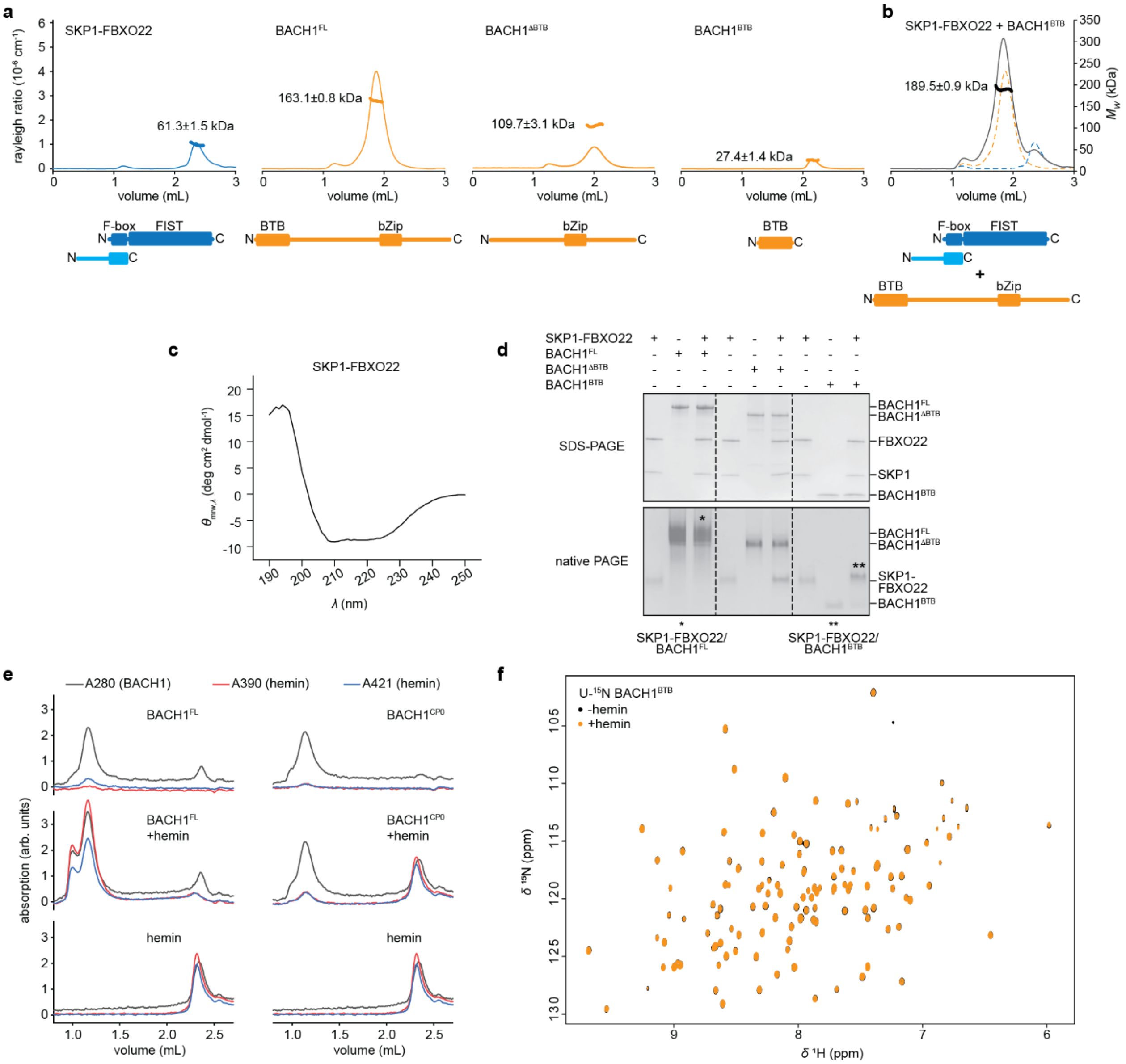

**Supplementary Figure 1: Characterization of SKP1-FBXO22 and BACH1 constructs.**

**a** SEC-MALS analysis confirms that recombinant SKP1-FBXO22 forms a heterodimer while recombinant BACH1$^{FL}$, BACH1$^{\Delta BTB}$, and BACH1$^{BTB}$ constructs form homodimers.

**b** SEC-MALS analysis of a mixture of BACH1$^{FL}$ with a fivefold molar excess of SKP1-FBXO22. The determined mass of 190 kDa is more consistent with a complex stoichiometry of a SKP1-FBXO22 bound to a BACH1$^{FL}$ dimer (227 kDa) than to a BACH1$^{FL}$ monomer (146 kDa).

**c** Circular dichroism (CD) spectrum of SKP1-FBXO22 indicating that the protein is correctly folded.

**d** Native PAGE (**c**) and fSEC analysis (**d**) of complex formation between SKP1-FBXO22 and BACH1 constructs. Proteins were used in equimolar ratios, i.e., one SKP1-FBXO22 heterodimer per BACH1 homodimer. Asterisks in (**c**) indicate co-migration of SKP1-FBXO22 and BACH1$^{FL}$ or BACH1$^{BTB}$.

**e** Analytical SEC to monitor hemin binding to BACH1. While BACH1 absorbs at 280 nm, hemin absorption is significant at 390 nm and 421 nm but also detectable at 280 nm. The shift of the hemin peak at 390 nm and 421 nm upon BACH1$^{FL}$ addition but not upon BACH1$^{CP0}$ addition confirms that hemin only binds to the CP motifs in the BACH1 C-terminus.

**f** ($^{1}H$,$^{15}N$)-TROSY NMR spectrum of 75 µM U-$^{15}N$ BACH1$^{BTB}$ recorded at 296 K in the absence and presence of 100 µM hemin.

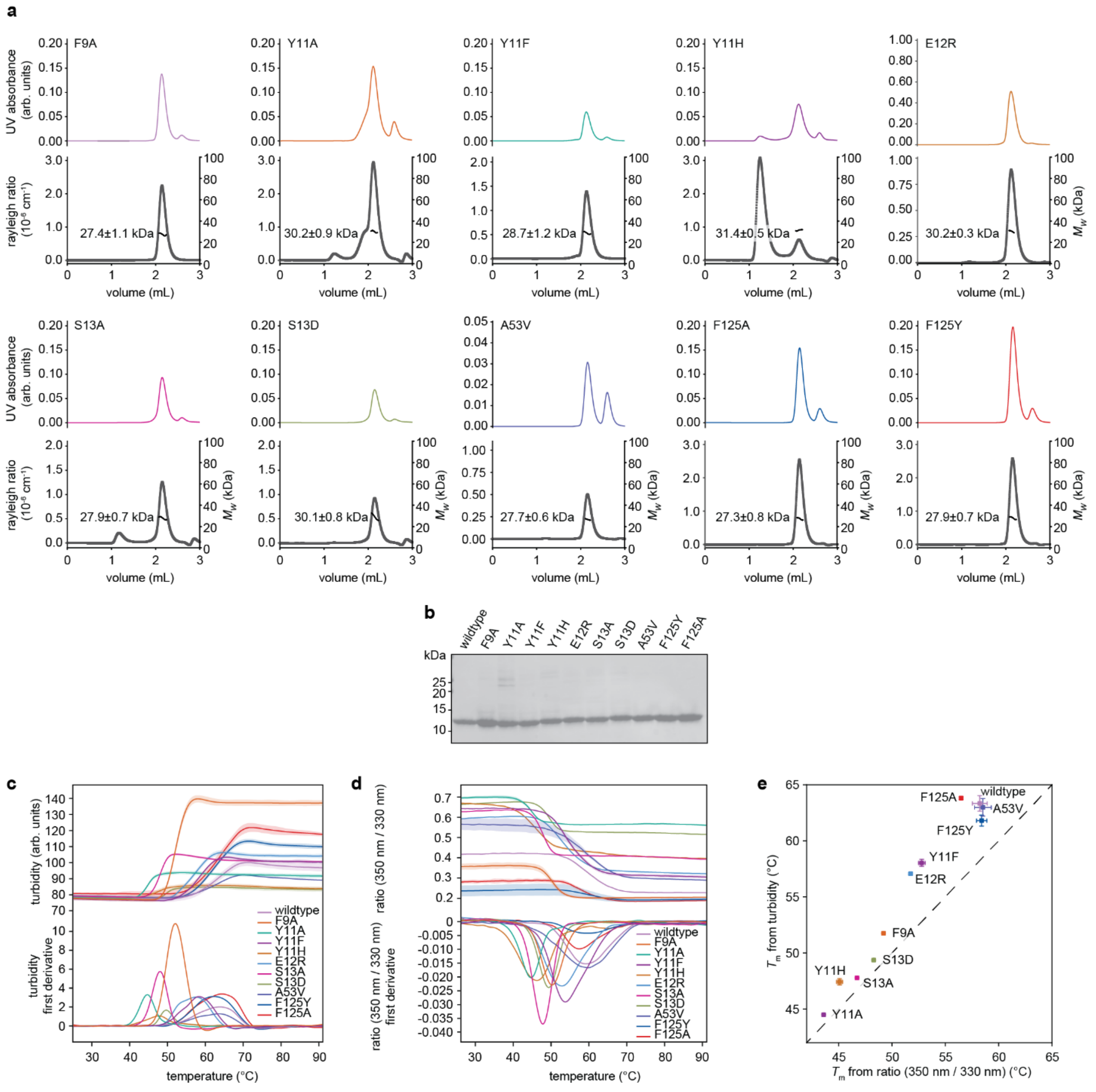

**Supplementary Figure 2: Characterization of BACH1$^{BTB}$ mutants.**

**a** SEC-MALS confirms the structural integrity of recombinantly produced BACH1$^{BTB}$ mutants F9A, Y11A, Y11F, E12R, S13A, S13D, A53C, F125A, F125Y (see Supplementary table 7 for determined molecular weights).

**b** Purity of recombinant BACH1$^{BTB}$ mutants demonstrated by SDS-PAGE followed by Coomassie staining.

**c, d, e** Nano differential scanning fluorimetry (nano DSF) analysis of BACH1$^{BTB}$ mutants. Melting temperatures, $T_m$, were determined from turbidity (**c**) and fluorescence (**d**) measurements (upper panel) using the Boltzmann and the derivative method (lower panel). Both methods yielded comparable melting temperatures for the measured mutants (**e**).

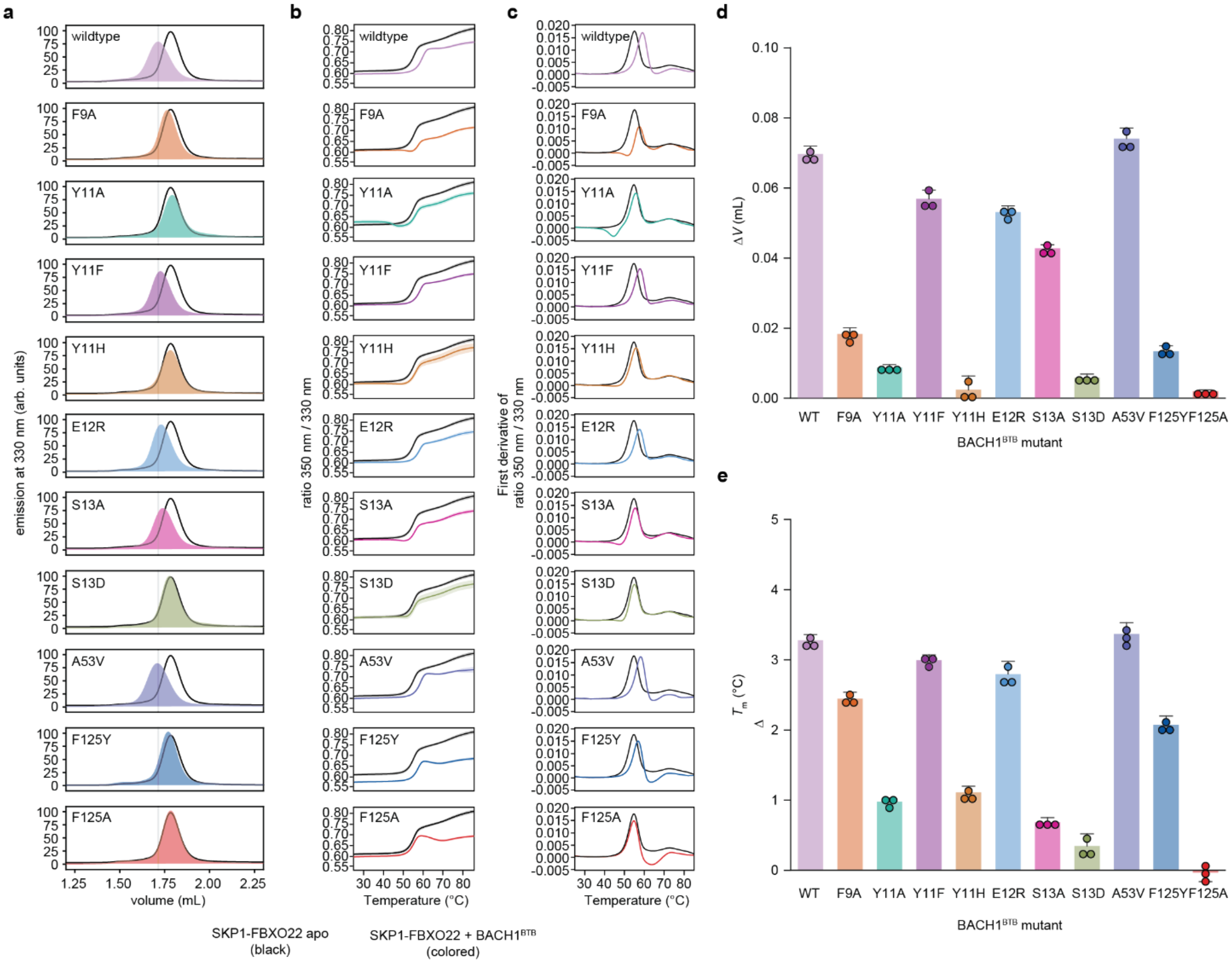

**Supplementary Figure 3: Fluorescence-detected size exclusion chromatography (fSEC) and nano differential scanning fluorimetry (nano DSF) analysis of the interaction between SKP1-FBXO22 and BACH1$^{BTB}$ mutants.**

**a** fSEC profiles of SKP1-FBXO22 and SKP1-FBXO22 in the presence of BACH1$^{BTB}$ wildtype or mutant constructs. Due to the absence of Trp residues in the BACH1 BTB domain, the fluorescence signal at 330 nm is solely from the SKP1-FBXO22 heterodimer.

**b** Nano DSF melting experiments of SKP1-FBXO22 apo (black) and in presence of BACH1$^{BTB}$ wildtype or mutant (colored). Due to the low abundance of aromatic side chains (no Trp) in the BACH1 BTB domain, the nano DSF signal at 350 nm and 33 nm is dominated by the fluorescence of SKP1-FBXO22. The solid lines represent the mean and the transparent error bands represent the standard deviation from n=3 experiments.

**c** Derivative of melting curves shown in (**b**) for melting temperature determination.

**d** fSEC shifts ($\Delta V$) from profiles in **a**. Values represent the mean and error bars represent the standard deviation from n=3 experiments.

**e** Melting temperatures ($T_m$) determined from nano DSF experiments in (**b**) and (**c**). Values represent the mean and error bars represent the standard deviation from n=3 experiments.

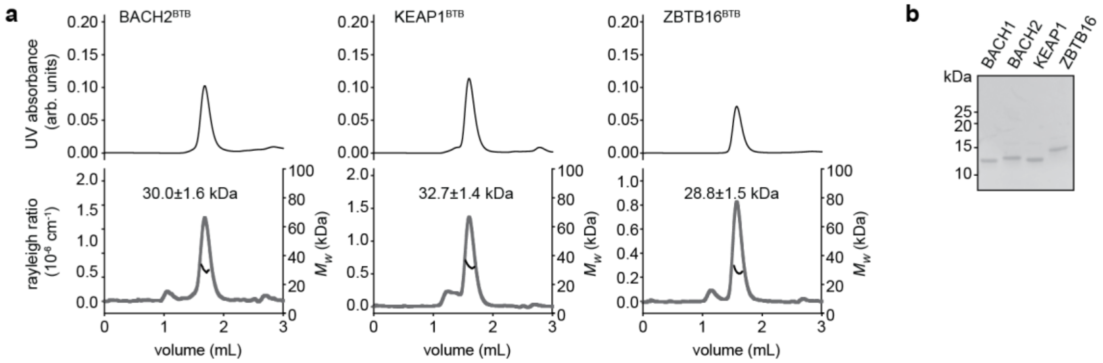

**Supplementary Figure 4: Characterization of the BACH2, KEAP1, and ZBTB16 BTB domains.**

**a** SEC-MALS confirms the structural integrity of the recombinantly produced BACH2, KEAP1, ZBTB16 BTB domains.

**b** Purity of recombinant BTB domains of BACH2, KEAP1, ZBTB16 demonstrated by SDS-PAGE followed by Coomassie staining.

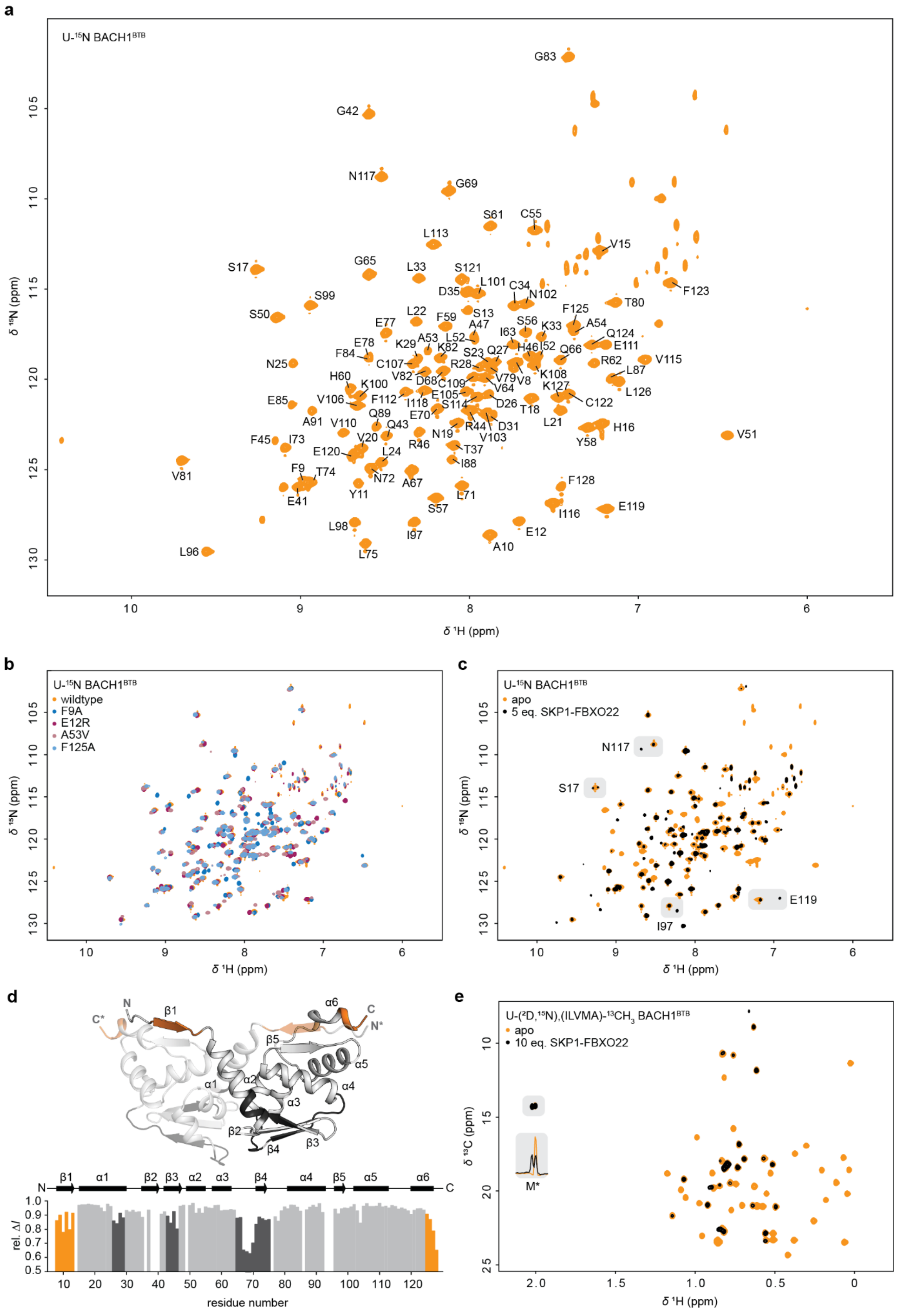

a
U-¹⁵N BACH1BTB
δ ¹⁵N (ppm)
δ ¹H (ppm)
b
U-¹⁵N BACH1BTB
wildtype
F9A
E12R
A53V
F125A
c
U-¹⁵N BACH1BTB
apo
5 eq. SKP1-FBXO22
N117
S17
E119
I97
d
residue number
rel. ΔI
e
U-(²D,¹⁵N),(ILVMA)-¹³CH3 BACH1BTB
apo
10 eq. SKP1-FBXO22
M*
δ ¹³C (ppm)

**Supplementary Figure 5: NMR spectroscopic analysis of BACH1BTB and its interaction with SKP1-FBXO22.**

**a** ($^{1}H$,$^{15}N$)-TROSY NMR spectrum of 50 µM U-$^{15}N$ BACH1$^{BTB}$ recorded at 296 K with backbone NMR assignment.

**b** Superimposed ($^{1}H$,$^{15}N$)-HSQC NMR spectra of U-$^{15}N$ BACH1$^{BTB}$ wildtype, and F9A, E12R, A53V, F125A mutants used to aid backbone NMR assignments of BACH1$^{BTB}$.

**c** Superimposed ($^{1}H$,$^{15}N$)-HSQC NMR spectra of U-$^{15}N$ BACH1$^{BTB}$ in the apo state (orange) and in complex with SKP1-FBXO22 (black).

**d** Line broadening analysis of spectra shown in (**c**). The differential intensity between the apo and FBXO22 bound state, $\Delta I$, is plotted against the amino acid sequence and mapped onto an AlphaFold2 structure of the BACH1 BTB domain[S1,S2]. A value of $\Delta I$=1 indicates full signal loss upon FBXO22 binding. Reduced $\Delta I$ values are detectable in loop regions (dark grey) as well as in the N-terminal β1 strand and the C-terminal α6 helix (orange).

**e** Superimposed ($^{1}H$,$^{13}C$)-HMQC NMR spectra of U-($^{2}D$,$^{15}N$), (ILVMA)-$^{13}CH_3$ labeled BACH1$^{BTB}$ in the apo state (orange) and in complex with SKP1-FBXO22 (black). The peak corresponding to the artificial N-terminal methionine residue (M*) is indicated with a grey box together as a $^{1}H$ 1D projection to highlight the peak splitting upon complex formation.

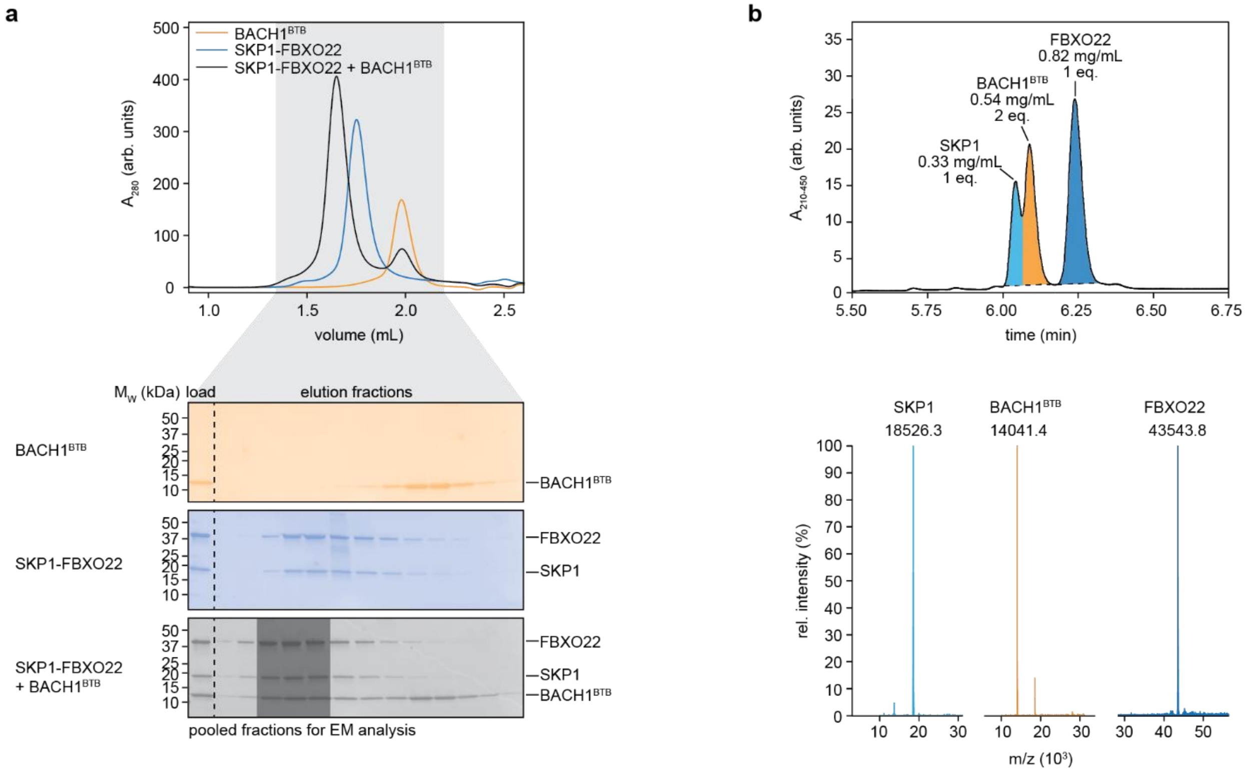

**Supplementary Figure 6: SKP1-FBXO22/BACH1$^{BTB}$ complex preparation for single particle cryo-electron microscopy analysis.**

**a** Preparation of the SKP1-FBXO22/BACH1$^{BTB}$ complex for single particle cryo-EM analysis via size exclusion chromatography (SEC). SKP1-FBXO22 was mixed with a 1.5-fold excess of BACH1$^{BTB}$ dimer, incubated, and the SKP1-FBXO22/BACH1$^{BTB}$ complex purified via SEC. The pooled fractions used for cryo-EM specimen preparation are highlighted with a shaded box.

**b** High pressure liquid chromatography coupled to mass spectrometry (HPLC-MS) of the pooled fractions from (**a**) confirmed the SKP1, FBXO22, BACH1$^{BTB}$ stoichiometry of 1:1:2 as previously seen for BACH1$^{FL}$ by MP and SEC-MALS.

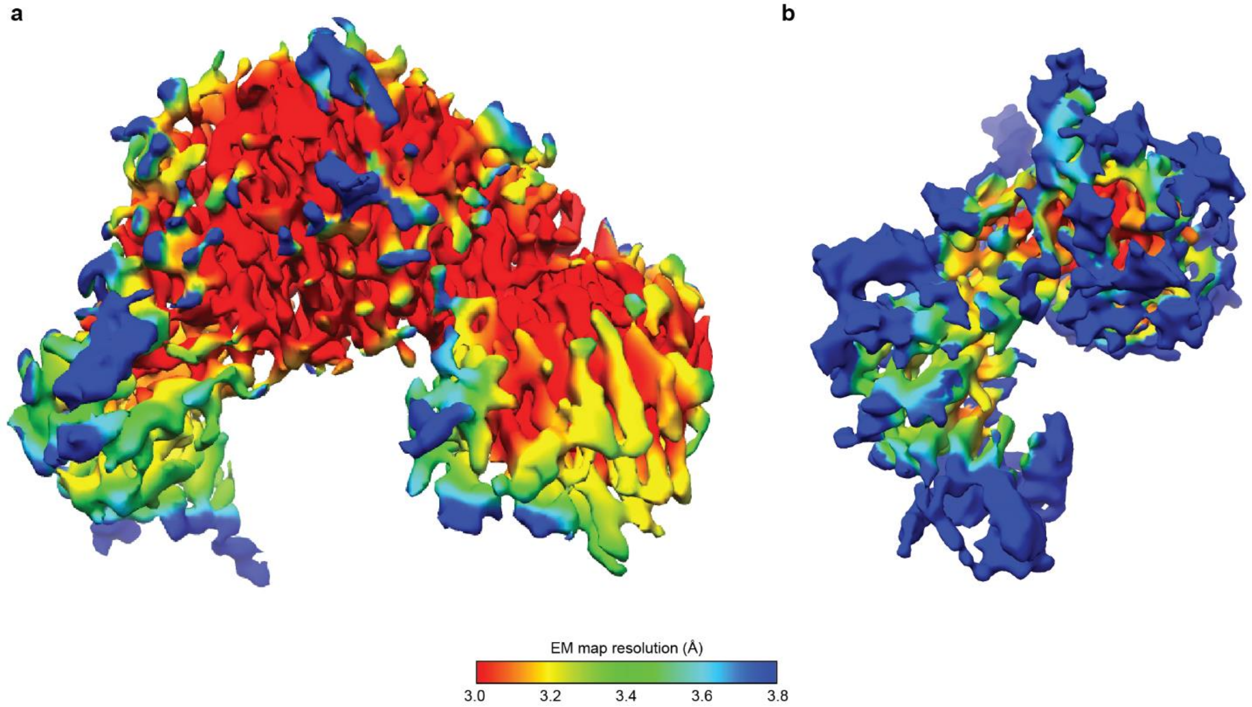

**Supplementary Figure 7: Cryo-EM resolution map of the SKP1-FBXO22 in the BACH1$^{BTB}$ bound and apo state.**

**a** EM resolution map of the SKP1-FBXO22/BACH1$^{BTB}$ complex. The average resolution is 3.2 Å.

**b** EM resolution map of SKP1-FBXO22. The average resolution is 3.4 Å.

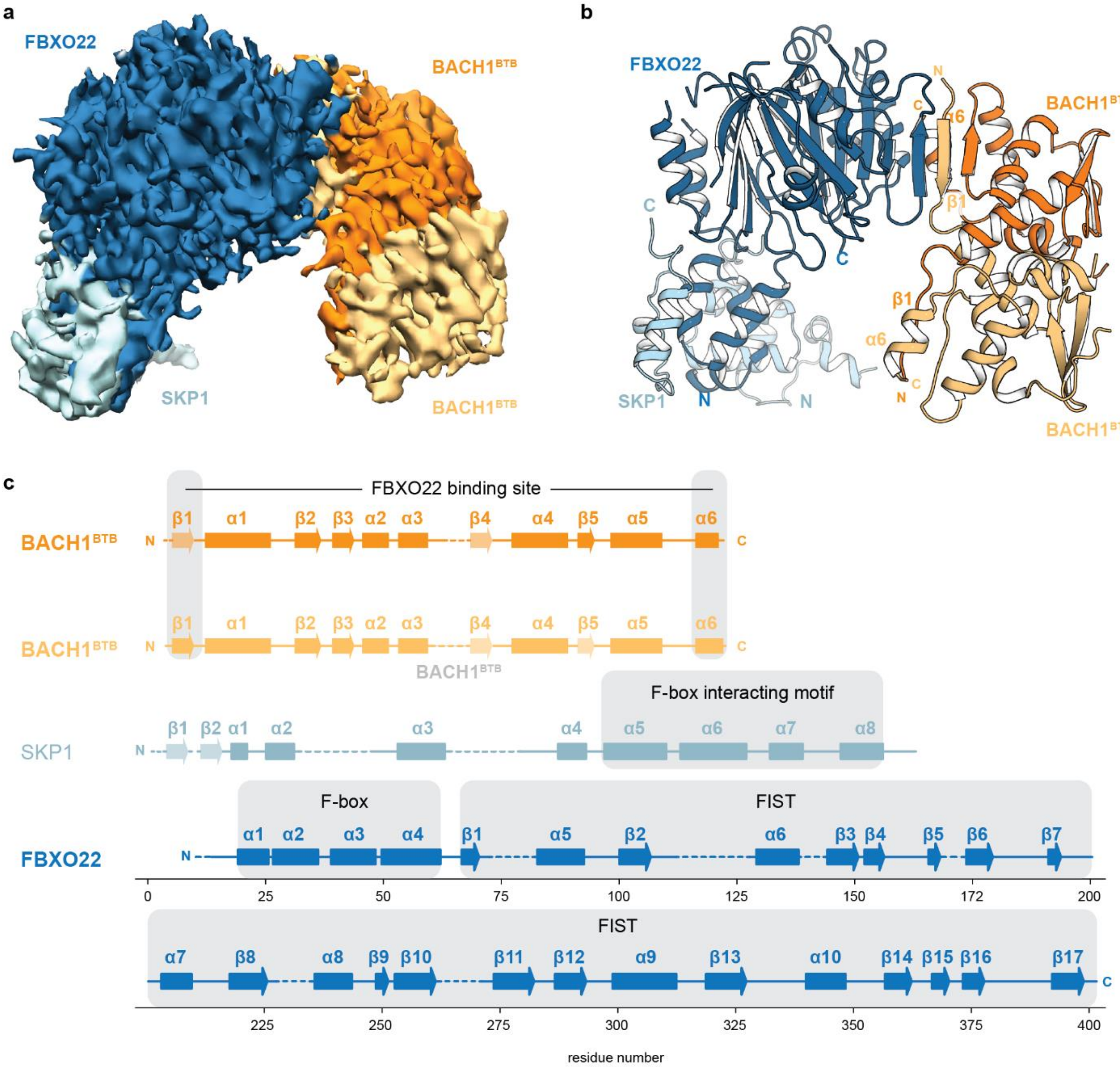

**Supplementary Figure 8: Cryo-EM structure of the SKP1-FBXO22/BACH1$^{BTB}$ complex.**

**a** Cryo-EM density map of the SKP1-FBXO22/BACH1$^{BTB}$ complex with the colored individual components.

**b** Structure of the SKP1-FBXO22/BACH1$^{BTB}$ complex obtained from the cryo-EM map shown in (**a**). The structure revealing the FBXO22 binding site in β1 of one subunit and helix α6 of the other subunit of the BACH1 BTB domain agrees well with the symmetry break suggested by the NMR peak splitting shown in Figure 2a and b.

**c** Secondary structure topology of SKP1, FBXO22 and BACH1BTB. Unresolved loops are shown as dashed lines. Secondary structure elements which are badly or not resolved but present in other available structures of SKP1 or BACH1 are transparent.

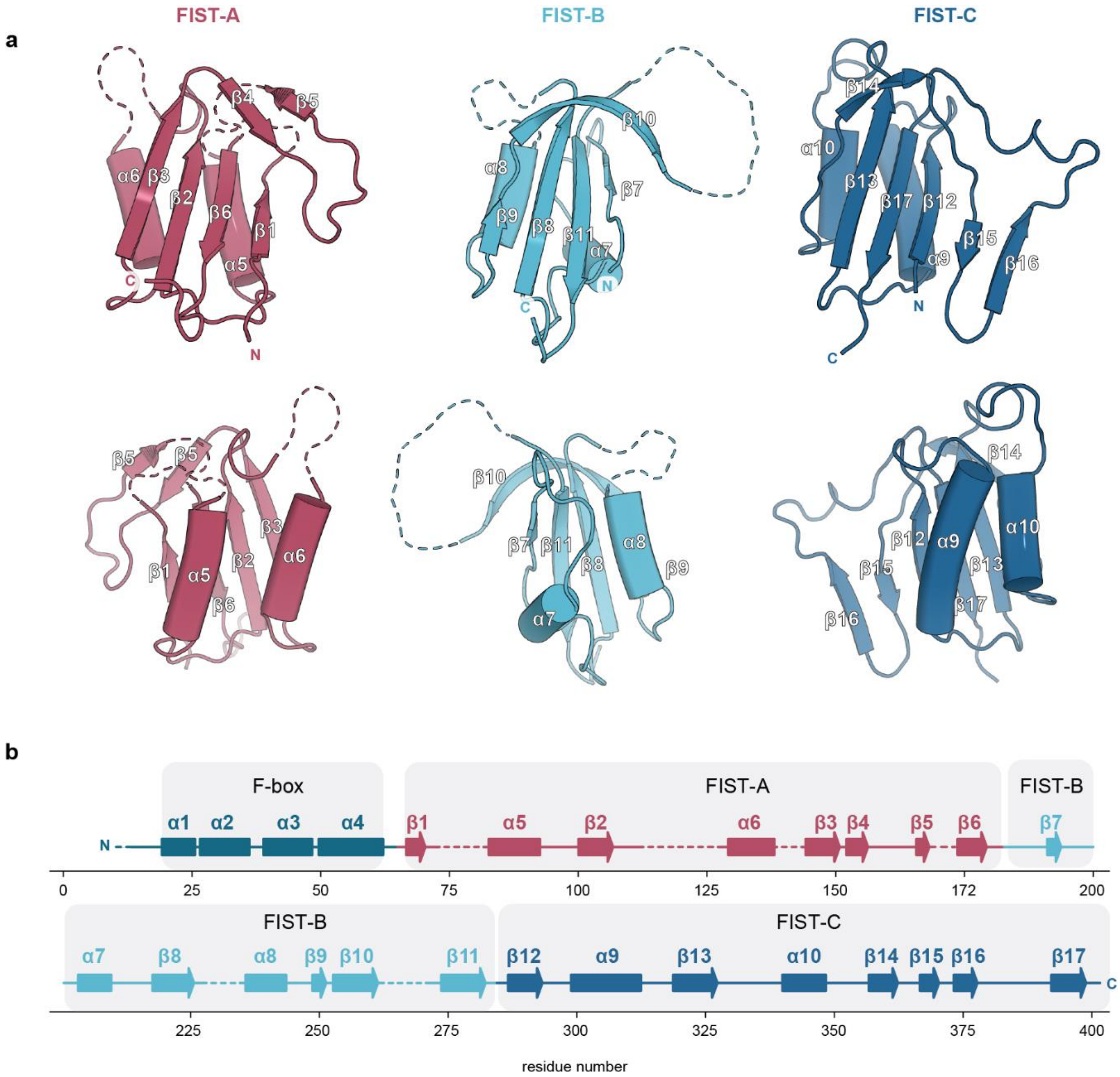

**Supplementary Figure 9: Domain architecture of the FBXO22 FIST domain.**

**a** Cartoon presentation of the β-α-β-α-β-loop-β fold of the FIST subdomains A, B, and C viewed from the core (top row) and the periphery of the FIST domain[S3].

**b** Secondary structure topology of FBXO22. Loops which are unresolved in the EM structures are shown as dashed lines.

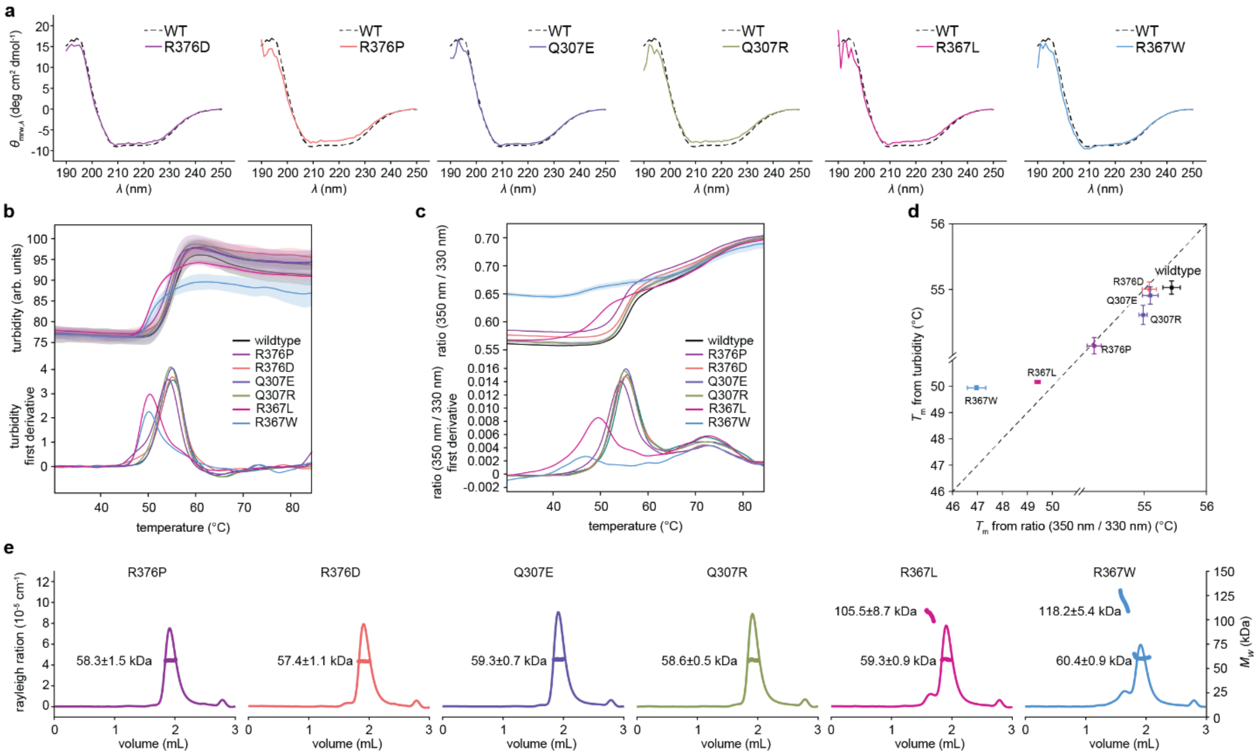

**Supplementary Figure 10: Characterization of SKP1-FBXO22 single point mutants.**

**a** Circular dichroism (CD) spectra of SKP1-FBXO22 wildtype and mutant constructs with single point mutations in FBXO22.

**b-d** Nano differential scanning fluorimetry (nano DSF) analysis of SKP1-FBXO22 mutants. Melting temperatures, $T_m$, were determined from turbidity (**b**) and fluorescence (**c**) measurements (upper panel) using the Boltzmann and the derivative method (lower panel). Both methods yielded comparable melting temperatures for the measured mutants (**d**).

**e** SEC-MALS analysis of SKP1-FBXO22 wildtype and mutants.

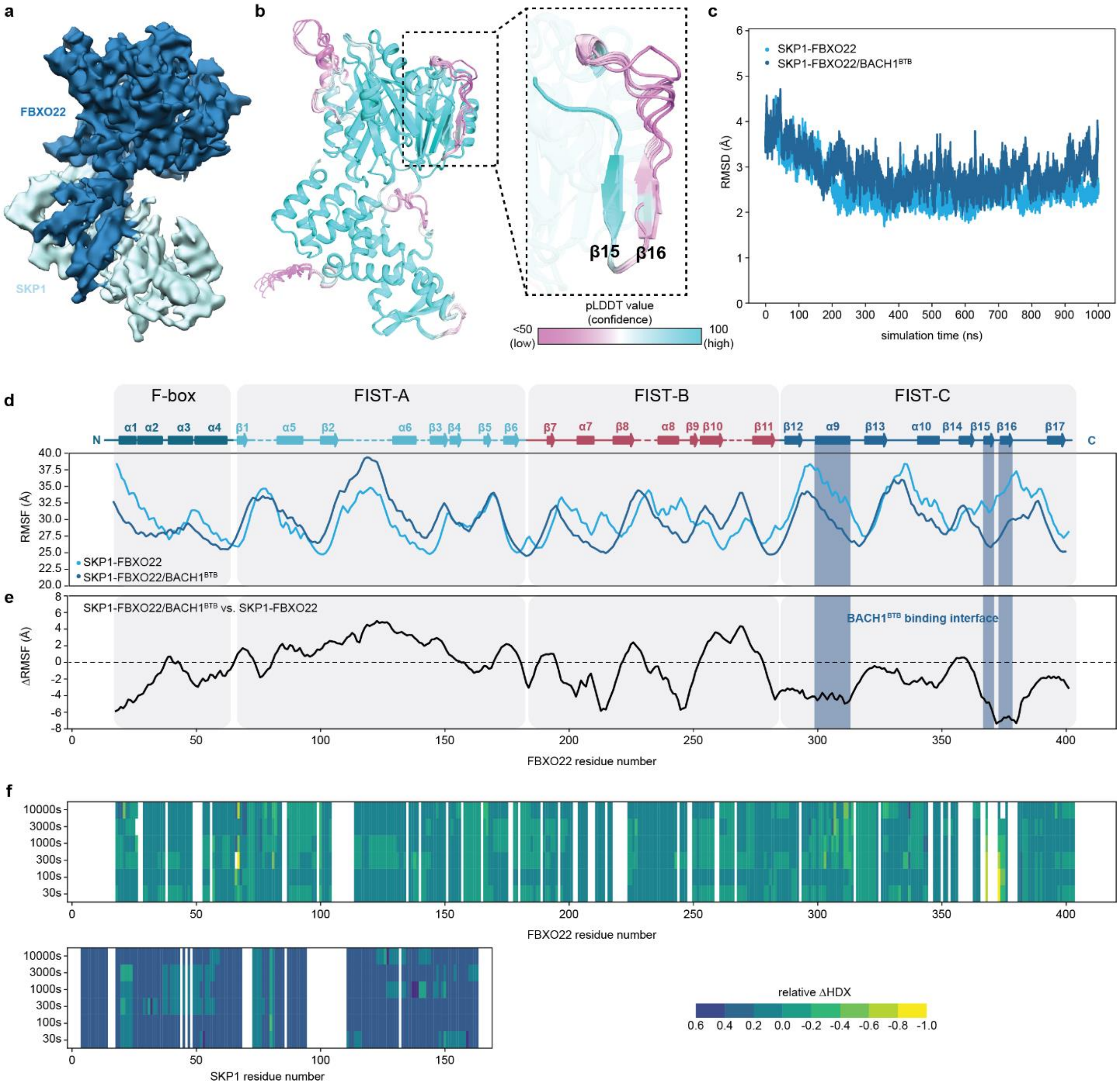

**Supplementary Figure 11: Analysis of the structural dynamics in SKP1-FBXO22.**

**a** Cryo-EM density of SKP1-FBXO22 with colored subunits.

**b** Six lowest energy AlphaFold2[S1,S2] models of SKP1-FBXO22. Low confidence (pLDDT) may suggest structural flexibility in β16 in the absence of BACH1[S4].

**c** Root mean square deviation (RMSD) plotted versus simulation time.

**d** Root-mean-square fluctuations (RMSF) during the 1 µs MD simulation plotted versus the FBXO22 sequence.

**e** Differential RMSF between SKP1-FBXO22 and SKP1-FBXO22/BACH1$^{BTB}$ shows significant changes in protein dynamics in the BACH1 binding site.

**f** Differential HDX-MS heat map showing the relative differential HDX between SKP1-FBXO22 apo versus SKP1-FBXO22/BACH1$^{BTB}$. Negative values indicate reduced flexibility upon BACH1$^{BTB}$ binding.

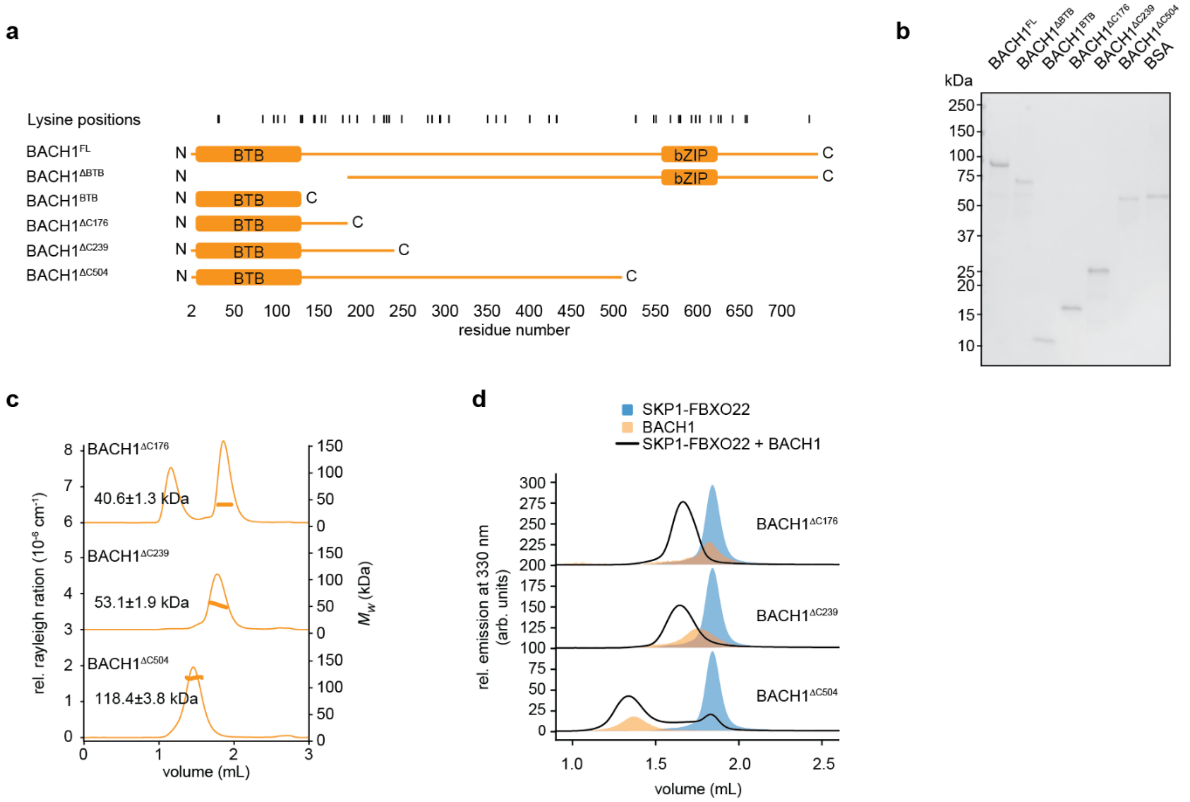

**Supplementary Figure 12: BACH1 deletion constructs used for the identification ubiquitylation sites in BACH1.**

**a** Overview of BACH1 deletion construct used in this study. Lysine positions are highlighted on top of the schematic.

**b** SDS-PAGE analysis of the BACH1 constructs shown in (a).

**c** SEC-MALS analysis of the BACH1 deletion constructs BACH1$^{\Delta C178}$, BACH1$^{\Delta C241}$, BACH1$^{\Delta C506}$ confirms their dimeric state.

**d** Complex formation of the BACH1 deletion constructs with SKP1-FBXO22 demonstrated by fSEC analysis.

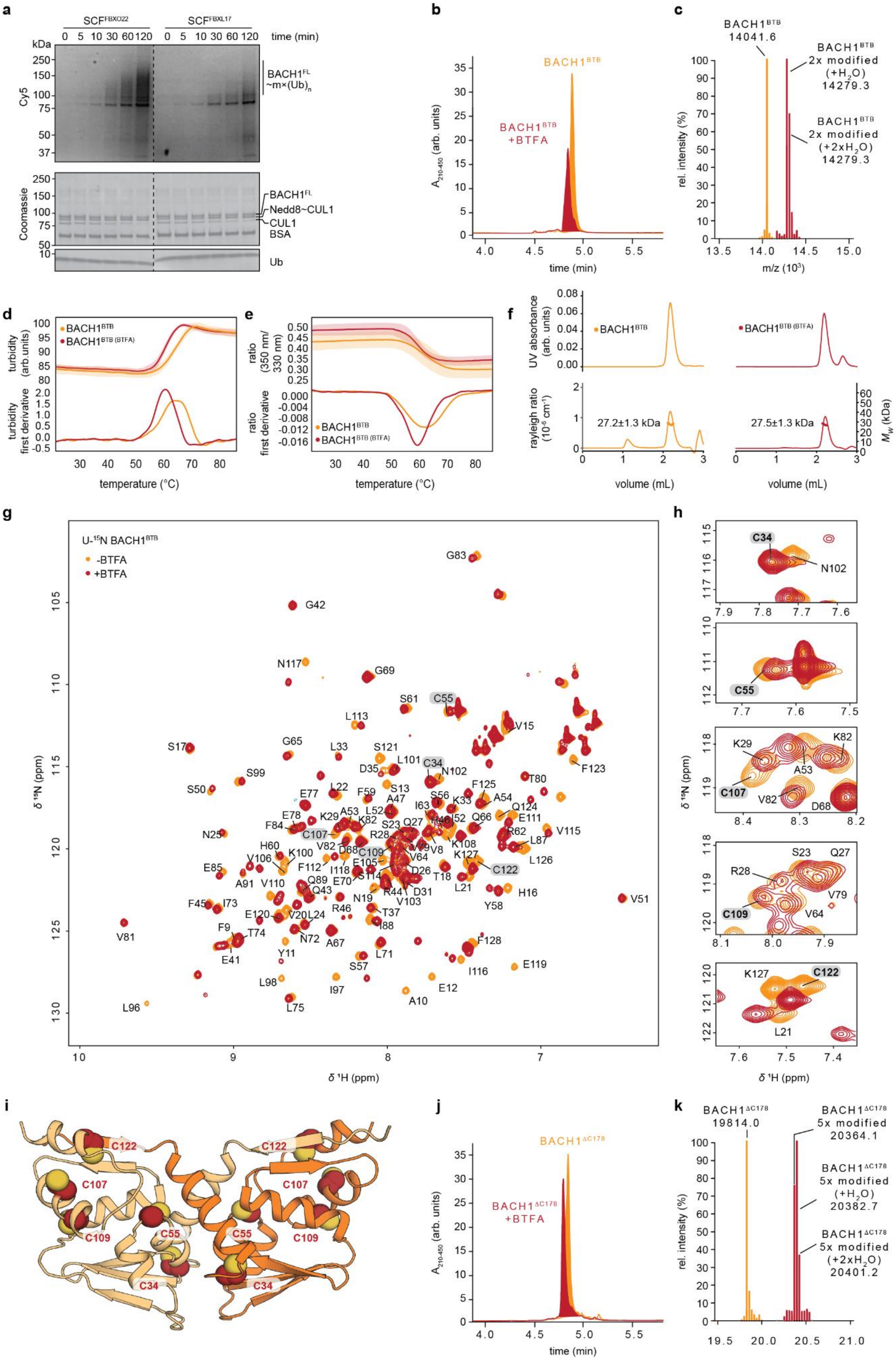
a
SCF^FBXO22
SCF^FBXL17
time (min)
kDa
Cy5
BACH1^FL ~m×(Ub)n
Coomassie
BACH1^FL
Nedd8~CUL1
CUL1
BSA
Ub
b
BACH1^BTB
BACH1^BTB +BTFA
A210-450 (arb. units)
time (min)
c
BACH1^BTB 14041.6
BACH1^BTB 2x modified (+H2O) 14279.3
BACH1^BTB 2x modified (+2xH2O) 14279.3
rel. intensity (%)
m/z (10^3)
d
turbidity (arb.units)
turbidity first derivative
temperature (°C)
e
ratio (350 nm/ 330 nm)
ratio first derivative
BACH1^BTB (BTFA)
f
UV absorbance (arb. units)
rayleigh ratio (10^-6 cm^-1)
27.2±1.3 kDa
27.5±1.3 kDa
Mw (kDa)
volume (mL)
g
U-15N BACH1^BTB
-BTFA
+BTFA
δ 15N (ppm)
δ 1H (ppm)
h
C34
N102
C55
C107
C109
C122
i
C122
C107
C109
C55
C34
j
BACH1^ΔC178
BACH1^ΔC178 +BTFA
k
BACH1^ΔC178 19814.0
BACH1^ΔC178 5x modified 20364.1
BACH1^ΔC178 5x modified (+H2O) 20382.7
BACH1^ΔC178 5x modified (+2xH2O) 20401.2

**Supplementary Figure 13: Cysteine modification of BACH1 constructs by Bromo-1,1,1-trifluoroacetate.**

**a** In vitro ubiquitination assay of BACH1$^{FL}$ by SCF$^{FBXO22}$ and SCF$^{FBXL17}$ over a time course of 2 hrs. Upper panel represents the fluorescence scan (Cy5) while the lower panel represents the Coomassie stained SDS-PAGE gel.

**b, c** LC-MS analysis of BACH1$^{BTB}$ and BTFA-labeled BACH1$^{BTB}$. The mass shift between BACH1$^{BTB}$ and BACH1$^{BTB(BTFA)}$ agrees with a twofold labeling with BTFA (110 Da per modification).

**d, e** Nano differential scanning fluorimetry (nano DSF) analysis of BACH1$^{BTB}$ modified with BTFA. Melting temperatures, $T_m$, were determined from turbidity (**d**) and fluorescence (**e**) measurements (upper panel) using the Boltzmann and the derivative method (lower panel).

**f** SEC-MALS analysis shows that the BACH1 BTB domain when modified with BTFA is still able to form dimers.

**g** Overlayed ($^{1}H$,$^{15}N$)-HSQC NMR spectra of (U-$^{15}N$)-BACH1$^{BTB}$ in the absence and presence of a 10x molar excess of Bromo-1,1,1-trifluoroacetate (BTFA). Substantial peak shifts can be observed for residues close to and within the degron sequence $^{9}$FAYES$^{12}$.

**h** Close up view of peaks in the BACH1$^{BTB}$ ($^{1}H$,$^{15}N$)-HSQC NMR spectra shown in (**d**) corresponding to cysteine residues C34, C55, C107, C109, and C122. Peak shifts for C107 and C122 but not for C34, C55, and C109 indicates selective modification at C107 and C122 yielding a twofold BTFA-modified BTB domain, in agreement with the LC-MS analysis.

**i** Cysteine residues shown as spheres in an AlphaFold2[S1,S2] model of the BACH1 BTB domain.

**j, k** LC-MS analysis of BTFA-labeled BACH1$^{\Delta C178}$ reveals a fivefold modification. This can be attributed to the modification of C107 and C122 in the BTB domain as well as C140, C145, and C150 in the disordered C-terminus. Of note, it cannot be excluded that modification of C140, C145, and C150 influences ubiquitination of BACH1$^{\Delta C178}$ at lysine residues.

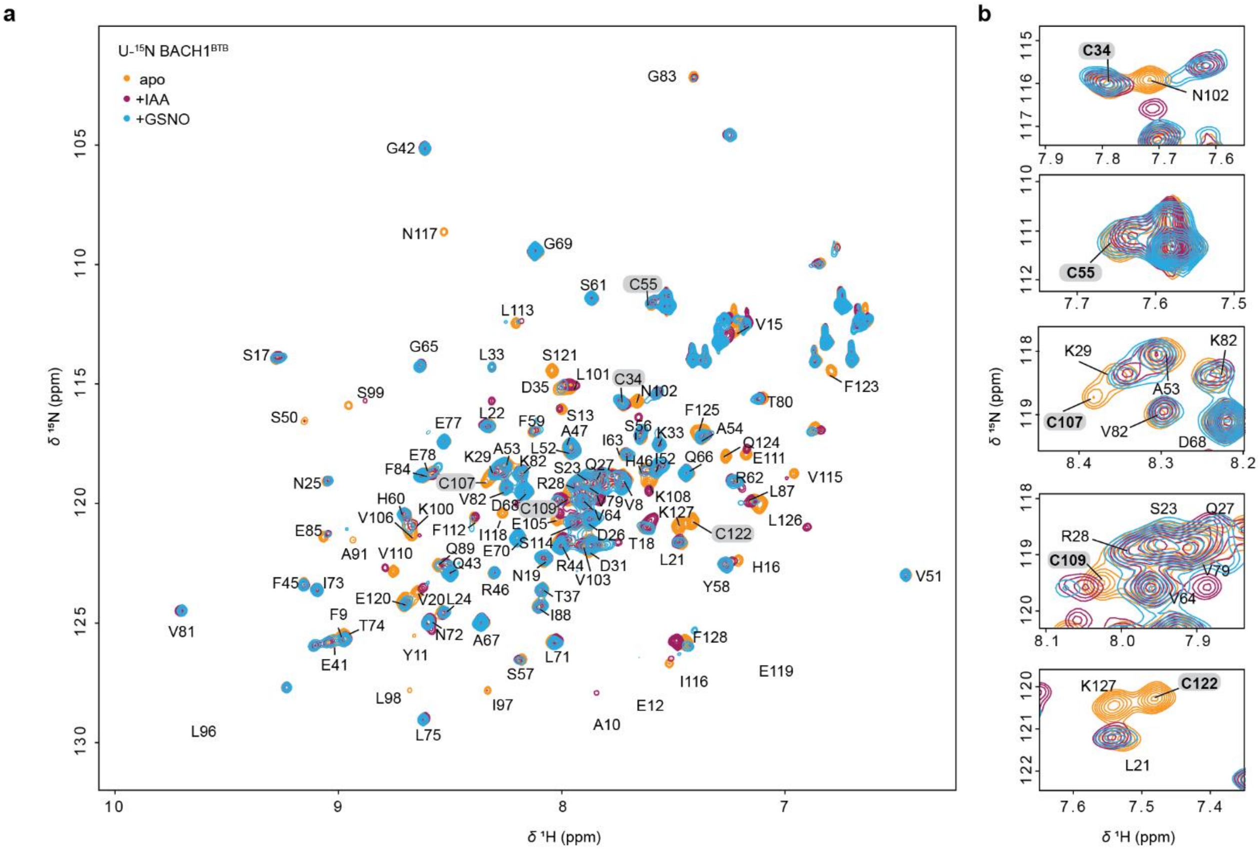

**Supplementary Figure 14: Cysteine modification by iodoacetamide and S-nitrosylglutathione monitored by NMR spectroscopy.**

**a** Overlayed ($^{1}H$,$^{15}N$)-HSQC NMR spectra of (U-$^{15}N$)-BACH1$^{BTB}$ in the absence and presence of a 10x molar excess of iodoacetamide (IAA) or S-nitrosylglutathione. The spectral changes strongly resemble those upon modification with BTFA.

**b** Close up view of peaks in the BACH1$^{BTB}$ HSQC spectra shown in (**a**) corresponding to cysteine residues C34, C55, C107, C109, and C122. The peak shifts for C107 and C122 but not for the remaining cysteine residues suggests that those are the sites for modification, like for BTFA (Supplementary Figure 13g).
